# Feasibility conditions of ecological models: Unfolding links between model parameters

**DOI:** 10.1101/2021.10.11.463747

**Authors:** Mohammad AlAdwani, Serguei Saavedra

## Abstract

Over more than 100 years, ecological research has been striving to derive internal and external conditions compatible with the coexistence of a given group of interacting species. To address this challenge, numerous studies have focused on developing ecological models and deriving the necessary conditions for species coexistence under equilibrium dynamics, namely feasibility. However, due to mathematical limitations, it has been impossible to derive analytic expressions if the isocline equations have five or more roots, which can be easily reached even in 2-species models. Here, we propose a general formalism to obtain the set of analytical conditions of feasibility for any polynomial population dynamics model of any dimension without the need to solve for the equilibrium locations. We additionally illustrate the application of our methodology by showing how it is possible to derive mathematical relationships between model parameters, while maintaining feasibility in modified Lotka-Volterra models with functional responses and higher-order interactions (model systems with at least five equilibrium points)—a task that is impossible to do with simulations. This work unlocks the opportunity to increase our systematic understanding of species coexistence.

## Introduction

Over more than 100 years, ecological research has been striving to derive the biotic and abiotic conditions compatible with the coexistence of a given group of interacting species (also known as an ecological system or community) (Tansley, 1920; Lotka, 1920; Volterra, 1926; Gause, 1932; Case, 2000). These conditions can provide the keys to understand the mechanisms responsible for the maintenance of biodiversity and the tolerance of ecological systems to external perturbations (Levins, 1968; Sugihara, 1994; Loreau and De Mazancourt, 2013; Kerr et al., 2002). Because of the complexity of this question, many efforts have been centered on developing phenomenological and mechanistic models to represent the dynamics of ecological systems and predict their behavior (MacArthur, 1970; Turchin, 2003; Svirezhev and Logofet, 1983; Vandermeer and Goldberg, 2013). However, even if we had knowledge about the exact equations governing the dynamics of interacting species, extracting and solving the set of conditions compatible with the coexistence of such species would remain a big mathematical challenge (Grilli et al., 2017; AlAdwani and Saavedra, 2020; Song et al., 2019). Indeed, most of the analytical work looking at these coexistence conditions has focused on relatively simple 2-species systems or strictly particular cases of higher-dimensional systems (Cox et al., 2010; Strogatz, 2015; Ong and Vandermeer, 2015; Barabás et al., 2018). In fact, even at the 2-species level, currently there is no general methodology that can provide us with a full analytical understanding about coexistence conditions for any arbitrary model (AlAdwani and Saavedra, 2020). Therefore, the majority of work has relied on numerical simulations (Valdovinos, 2019; Letten and Stouffer, 2019), which only provide a partial view of the dynamics conditioned by the choice of parameter values (AlAdwani and Saavedra, 2019).

Recent work has started to address the challenge above by focusing on the necessary conditions for species coexistence under equilibrium dynamics: feasibility (Hofbauer and Sigmund, 1998; Song et al., 2018). Mathematically, the feasibility of a generic *n*-species dynamical system *dN*_*i*_/*dt* = *N*_*i*_*f*_*i*_(**N**)/*q*_*i*_(**N**), where the *f*’s and *q*’s are multivariate polynomials in species abundances **N** = (*N*_1_, *N*_2_, …, *N*_*n*_)^*T*^, corresponds to the existence of at least one equilibrium point (i.e., *dN*_*i*_/*dt* = 0 ∀*i*) whose components are all real and positive (i.e., 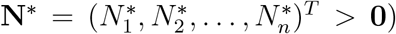. Feasibility conditions are typically represented by inequalities as a function of model parameters (Vandermeer, 1975; Barabás et al., 2018).

Traditionally, feasibility conditions have been attained by finding the isocline equations *f*_*i*_(**N**\*) = 0 ∀*i* and then solving for **N**\* before finding the conditions that satisfy **N**\* > **0** (Strogatz, 2015; Case, 2000; Vandermeer and Goldberg, 2013).

For example, let us focus on the linear Lotka-Volterra (LV) model of the form 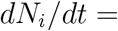 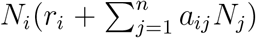, where *a*’s and *r*’s represent the interaction coefficients and the intrinsic growth rates, respectively. In the linear LV model, the isocline equations (for any dimension) can be written as **r** + **AN**\* = **0**, whose unique root is given by **N**\* = −**A**^−1^**r**. Therefore, feasibility conditions in this case are simply given by the inequality −**A**^−1^**r** > **0**. However, adding nonlinear functional responses or higher-order terms can increase exponentially the number of roots of the system (AlAdwani and Saavedra, 2019). Importantly, it can be shown from elimination theory (via Grobner basis) and Abel’s impossibility theorem that it is impossible to solve analytically for **N**\* when the number of roots of the system is five or more (Abel, 1824, 1826; Adams et al., 1994). Similarly, using numerical approaches, it has been demonstrated that the probability of feasibility (the probability of finding at least one equilibrium point whose components are all positive by randomly choosing parameter values) is an increasing function of the model’s complexity (i.e., number of complex roots of the isocline equations with generic coefficients) regardless of the invoked mechanism, whether they are ecologically motivated or have no meaning whatsoever (AlAdwani and Saavedra, 2020). This implies that traditional approaches can be unsuitable for finding the necessary conditions for coexistence in generic systems.

Here, we propose a general formalism to obtain for any polynomial population dynamics model and any dimension the set of necessary conditions leading to species coexistence without the need to solve for the equilibrium locations. We show how to reduce these conditions into a small set of expressions. We illustrate this methodology with an example of a univariate system. Additionally, we show how to identify the feasibility conditions that are compatible with a given range of parameter values. That is, we show how to find analytic relationships between model parameters while maintaining feasibility. We illustrate this methodology with examples of multispecies systems using modified LV models with functional responses and higher-order interactions, where isocline analysis cannot be performed. Finally, we discuss advantages and limitations of our formalism, and future avenues of research derived from our study.

## Obtaining feasibility conditions

Our methodology can be applied to any dynamical system of the form:

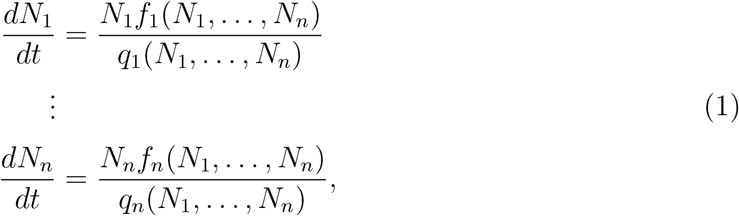

where the *f*’s and *q*’s are multivariate polynomials in species abundances. Let **Ψ** be the vector of model parameters that include, for example, species growth rates and species interaction coefficients. Feasibility conditions become consequently conditions on model parameters **Ψ** that guarantee at least one feasible equilibrium point in the system (Svirezhev and Logofet, 1983; AlAdwani and Saavedra, 2020). That is, we require that the number of roots of the system defined by polynomial equations *f*_*i*_(*N*_1_, …, *N*_*n*_) = 0 for *i* = 1, …, *n* whose components are all real and positive is at least one. To find such feasibility conditions, we develop a 3-step methodology: (1) Find symmetric sums of the roots of the polynomial. (2) Assemble a function that counts the number of of feasible roots. (3) Use the function of the number of feasible roots to deduce feasibility conditions, reduce them and eliminate redundant conditions. Below, we give details of these three steps. We also provide MATLAB code for its implementation.

### Finding symmetric sums of the roots

The first step involves in finding the symmetric sums of the roots that are needed to build the analytic formula of the number of feasible roots. Such sums can be obtained via different methodologies (Serret, 1849; Macaulay, 1902; Pedersen, 1991). One approach is outlined below:

1. Fix *i*, assume that variable *N*_*i*_ is constant, and find the total degree of each polynomial equation *f*_*j*_(*N*_1_, …, *N*_*n*_) = 0 for *j* = 1, …, *n*. The total degree of *f*_*j*_ is the maximum sum of the variables’ exponents in each term of *f*_*j*_ while treating *N*_*i*_ as constant. Denote the total degree of polynomial *f*_*j*_ by *d*_*i,j*_ for *j* = 1, …, *n*. Next, homogenize each term in each of the *f*’s with an artificial variable *W* so that the total degree of each term in *f*_*j*_ is *d*_*i,j*_. Denote to the homogenized equation by 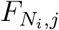. For example, if 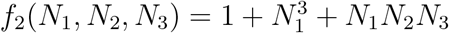 and *N*_1_ is assumed to be constant, then *d*_1,2_ = 2 and the homogenized equation is 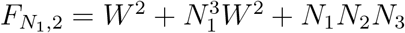.
2. Let 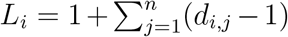 and form the set *H*_*i*_ as a union of *n* monomial sets, where 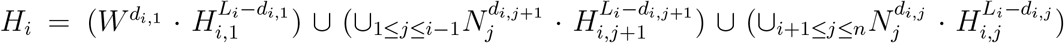. Define the outer-term of 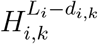 to be the one that is dotted or multiplied by it. For example 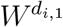 is the outer-term of 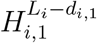. Here, 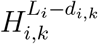 is the set of all monomials in *W, N*_1_, …, *N*_*n*_ not including *N*_*i*_ that are of total degree *L*_*i*_ −*d*_*i,k*_ and does not contain the outer-terms of any of 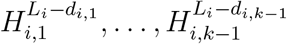. For example, if *d*_2,1_ = 2, *d*_2,2_ = 2 and *d*_2,3_ = 1, then using variables *W, N*_1_, *N*_3_ where *N*_2_ is constant, we have *L*_2_ = 3 and *H*_2_ = *W* ^2^ · {*W, N*_1_, *N*_3_} ∪ 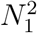 · {*W, N*_1_, *N*_3_} ∪ *N*_3_ · {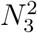, *WN*_1_, *WN*_1_, *N*_1_*N*_3_}. that the second curly bracket does not contain *W* ^2^ (i.e., outer term of the first curly bracket) and the third curly bracket does not contain *W*^2^ nor 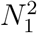 (i.e., the outer-terms of the first and second curly brackets).
3. Form the set 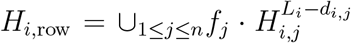 evaluated at *W* = 1. Note that ita*H*_*i,row*_ is simply *H*_*i*_ with outer-term of every 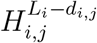 being replaced by *f*_*j*_. Next, form the monomial set *H*_*i,col*_ which is simply *H*_*i*_ evaluated at *W* = 1. After that, form the Macaulay matrix 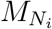, which is a square matrix whose size is 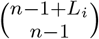 and whose (*i, j*) entry is the coefficient of *H*_*i,col*_(*j*) in the expression of *H*_*i,row*_(*i*) assuming that *N*_*i*_ is a constant. Then, find the resultant 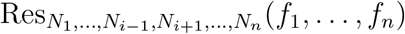 which equals to the determinant of 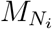. This resultant is a univariate polynomial in *N_i_* that contains no other *N*’s.
4. Next, form the matrix 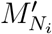, whose first column is *H*_*i,row*_ and its remaining columns *i* are the remaining columns of the matrix 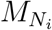. Then, compute its determinant (i.e., 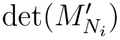), which has the form *T*_*i*1_*f*_1_ + *T*_*i*2_*f*_2_ + … + *T*_*in*_*f*_*n*_ to obtain the *i*^th^ row of the eliminant matrix. Repeat all previous steps for *i* = 1, …, *n* to obtain all entries of the eliminant matrix as well as all resultants. Then, obtain the Jacobian of the original polynomial system whose (*i, j*) entry is *∂f*_*i*_/*∂N*_*j*_. Next, find the determinant of both the eliminant matrix *T* and the determinant of the Jacobian *J*.
5. If the determinant of 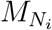 is 0, use the generalized characteristic polynomial formalism (Canny, 1988) to obtain the resultant. In this case, the resultant is the non-vanishing coefficient of the smallest power of *ϵ* in 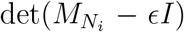, where *I* is the identity matrix of same size as matrix 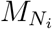. To find *T*_*ij*_ for *j* = 1, …, *n*, form the matrix 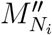, whose first column is *H*_*i,row*_ and its remaining columns are the remaining columns of the matrix 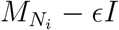. Then, compute its determinant and find the first non-zero coefficient of powers of *ϵ* in ascending order, which has the form *T*_*i*1_*f*_1_ + *T*_*i*2_*f*_2_ + … + *T*_*in*_*f*_*n*_ (see Appendix 5 for an example of this scenario).
6. Expand the generating function *G*(*f*_1_(*N*_1_, …, *N*_*n*_), …, *f*_*n*_(*N*_1_, …, *N*_*n*_)) that is shown below, around *N*_1_ = ∞, …, *N*_*n*_ = ∞ to obtain the Σ’s (symmetric sums of the roots).

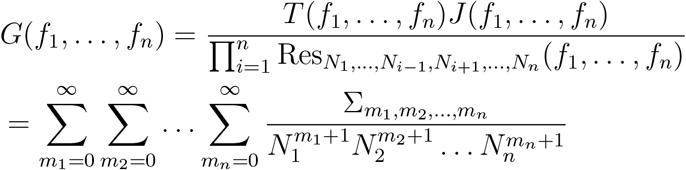

The expansion of *G* is done via performing series expansion of the reciprocal of each resultant separately then multiplying them along with *T* and *J*. For example, the reciprocal of each resultant can be expanded via MATLAB’s “taylor” command after performing change of variables *N*_*i*_ = 1/*x*_*i*_ and expanding around *x*_*i*_ = Alternatively, if the resultant is expressed as 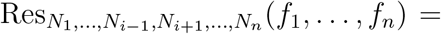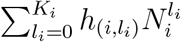, then

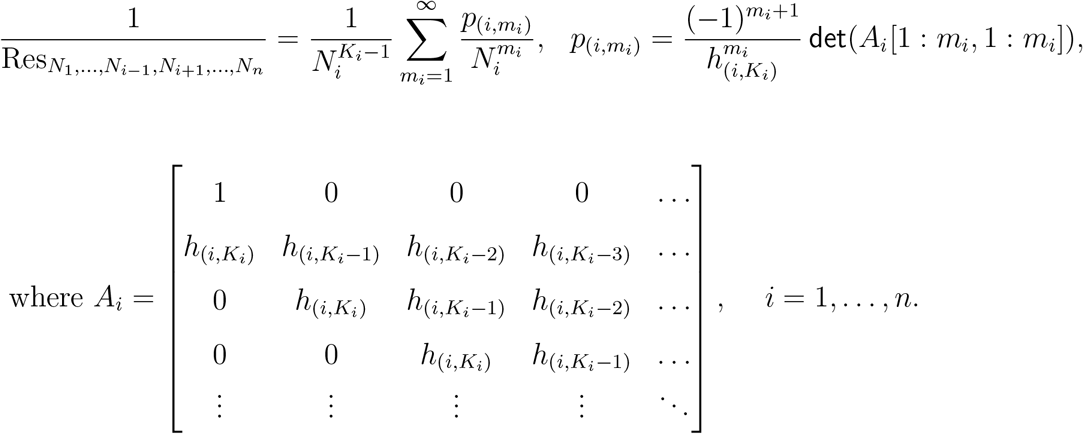

Finally, denote the roots of *f*_*i*_(*N*_1_, …, *N*_*n*_) for *i* = 1, …, *n* by ***η***_***k***_ = [*η*_***k***,1_, *η*_***k***,2_, …, *η*_***k***,*n*_]^*T*^ for *k* = 1, …, Θ. The symmetric sum 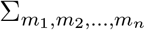 is given by 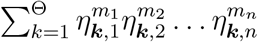 In particular, note that Θ = Σ_0,0,…,0_ is the number of complex roots of *f*_*i*_(*N*_1_, …, *N*_*n*_) for *i* = 1, …, *n* with general coefficients. It is important to record that number.

It is worth mentioning that the previous steps in univariate systems reduce significantly, where the roots of *f* (*N*) are considered. The jacobian determinant simply becomes *J* = *f*′(*N*) and the resultant is *f* (*N*) itself given that it is the only univariate polynomial in the system. In turn, the eliminant determinant is *T* = 1 as the resultant, where written in the form *T*_11_*f* (*N*) implies *T*_11_ = 1. Thus, the generating function reduces to *G* = *f* ′(*N*)/*f* (*N*) (Appendix 1 illustrates a simplified and detailed methodology for univariate systems). Similarly, in 2-dimensional systems, the two resultants simplify significantly and become determinants of Sylvester matrices involving the coefficients of two polynomial inputs. Then, to find the corresponding eliminant matrix, it is possible to modify a single column in each of the two Sylvester matrices without changing their determinant to write the resultants in the form *T*_*i*1_*f*_1_ + *T*_*i*2_*f*_2_ (Appendix 2 illustrates a simplified and detailed methodology for 2-species systems). For higher dimensional systems, we need to find the symmetric sums as described above or any other suitable implementation.

### Assembling the function that counts the number of feasible roots

Once we find the symmetric sums of the roots, we construct an analytical formula of the number of positive roots of the polynomial system of equations—we call that function *F* (**Ψ**). To derive *F* (**Ψ**), we apply previous work (Pedersen et al., 1993), which deals with counting real roots in arbitrary domains, to count the number of real roots in an orthotope that lies in the first quadrant (i.e., feasible region), which rests on all the positive axes. Then, we expand the orthotope allowing all non-zero components of all its vertices to go to infinity to cover the entire feasible domain. This can be achieved as follows:

1. Choose a map *m*(*N*_1_, *N*_2_, …, *N*_*n*_) of length Θ and with independent monomial entries. Typically, the first entry of *m* is the constant 1. Note that such monomials are chosen so that the coefficients of the characteristic equation shown in the following step do not vanish. Next, let *Q*(*N*_1_, *N*_2_, …, *N*_*n*_) = *N*_1_*N*_2_ …, *N*_*n*_ and compute the symmetric matrix *S*(*s*_1_, *s*_2_, …, *s*_*n*_) = *W* Δ*W* ^*t*^ where *W*_*ij*_ = *m*_*i*_(*η*_***j***,1_, *η*_***j***,2_, …, *η*_***j***,*n*_) and Δ_*ii*_ = *Q*(*η*_***i***,1_ − *s*_1_, *η*_***i***,2_ − *s*_2_, …, *η*_***i***,*n*_ − *s*_*n*_) is a diagonal matrix.
2. The next task is to evaluate the determinant of *S*(*s*_1_, *s*_2_, …, *s*_*n*_) and write it in the form det(*S*(*s*_1_, *s*_2_, …, *s*_*n*_) − *λI*) = (−1)^Θ^*λ*^Θ^ + *v*_Θ−1_(*s*_1_, *s*_2_, …, *s*_*n*_)*λ*^Θ−1^ + … + *v*_0_(*s*_1_, *s*_2_, …, *s*_*n*_). Then, consider the sequence **v** = [*v*_Θ_(*s*_1_, *s*_2_, …, *s*_*n*_) = (−1)^Θ^, *v*_Θ−1_(*s*_1_, *s*_2_, …, *s*_*n*_), …, *v*_0_(*s*_1_, *s*_2_, …, *s*_*n*_)] and let *V* (*s*_1_, *s*_2_, …, *s*_*n*_) be the number of consecutive sign changes in **v**. The formula of *V* (*s*_1_, *s*_2_, …, *s*_*n*_) is

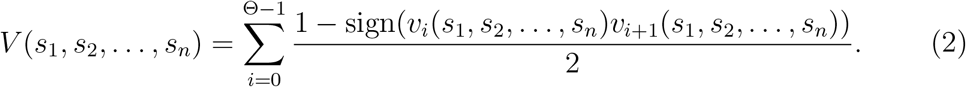
3. Consider the feasibility domain and think about it as a box whose 2^*n*^ vertices compose of zeros and infinities. Note that *v*_*i*_(*m*_1_, *m*_2_, …, *m*_*n*_), where *m*_1_, *m*_2_, …, *m*_*n*_ ∈ {0, ∞} is the coefficient of the highest power of 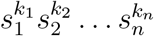 in *v*_*i*_(*s*_1_, *s*_2_, …, *s*_*n*_) where *k*_*i*_ = 0 if *m*_*i*_ = 0 and *k*_*i*_ = 1 if *m*_*i*_ = ∞. Finally, let #(*s*_1_, *s*_2_, …, *s*_*n*_) be the number of infinities that appear in the string *s*_1_, *s*_2_, …, *s*_*n*_. The expression of *F* (**Ψ**) is given by

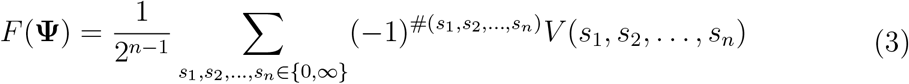

### Deducing feasibility conditions and reducing them

The third and last step of our methodology involves deducing feasibility conditions and reducing them. This has the purpose of unveiling the key inequalities that need to be satisfied in order to reach feasibility. This can be achieved as follows:

1. Call *v*_*i*_(*m*_1_, *m*_2_, …, *m*_*n*_), where *m*_1_, *m*_2_, …, *m*_*n*_ ∈ {0, ∞} and *i* = 0, 1, …, Θ − 1 forms the feasibility basis involving Θ2^*n*^ quantities (feasibility conditions are only dependent on those quantities). Because there are Θ2^*n*^ quantities and each can take a positive or a negative sign (we neglect the zero case as the values of ecological parameters are never exact), then there are 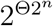 sign combinations. Many of those combinations are impossible to occur (empty) for any choice of real **Ψ**. To detect the non-empty sign combinations, compute the signs of all the *v*’s (the feasibility basis) as well as *F* (**Ψ**) for a range of parameters **Ψ**, where each component of **Ψ** varies independently in a large domain (say uniformly between −100 and 100 or in any suitable domain) when parameters are unrestricted. If one or more parameters are restricted, they need to be varied randomly in the domains they are defined at. This operation can be easily computed as it is only necessary to evaluate a few functions without solving systems of equations. Next, extract unique sign combinations of the *v*’s, which yield *F* (**Ψ**) ≥ 1. Then, put these sign combinations in a feasibility table (i.e., matrix), whose rows are the signs of the *v*’s and columns are the individual feasibility conditions.
2. After forming the feasibility table, perform a minimization to it. Here, we illustrate a simple minimization technique: If two columns differ by a single sign (in one row), the two columns are combined into one and an X (or 0) is placed in the row where there is a single sign difference. We repeat the same process until no two columns differ by a single sign. Next, we go through a single column at a time and iterate through each quantity in the basis. Then, we compute the conditional probabilities where the quantity takes its correspondent sign given that all remaining quantities have their correspondent signs. If one or more conditional probabilities are 1, the sign of one of those quantities may be replaced by **X** in the table. We then repeat computing the same conditional probabilities, which were 1 but without the **X**’ed quantity being part of the calculation. We repeat the process until no conditional probability is one. We then go through all columns and repeat the same process until it terminates. It is worth noting that these are not the only minimization approaches. For instance, comparing signs of *v*’s with *F* (**Ψ**) may reveal to us redundant quantities in the system (see the examples in Appendices 3–6).

### Illustrative Example

We illustrate the methodology above using the following univariate system:

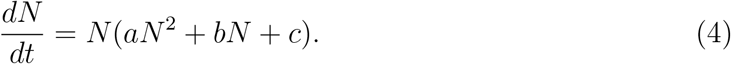

First, we find the symmetric sums of roots. For this purpose, let us focus on the quadratic polynomial *f* (*N*) = *aN* ^2^+*bN* +*c* of the equation above with model parameters **Ψ** = (*a, b, c*). This example has the same mathematical form of a a population model with an Allee effect (Case, 2000; Sun, 2016). Denote the two roots of *f* (*N*) by *η*_1_ and *η*_2_. Let *m*(*N*) = [1, *N*] be a monomial map of length *n* = 2 and *Q*(*N*) = *N*. Now, we can compute the matrix *S*(*s*_1_) = *W* Δ*W* ^*t*^, where *W*_*ij*_ = *m*_*i*_(*η_j_*) and Δ_*ii*_ = *Q*(*η*_*i*_ − *s*_1_) is the diagonal matrix

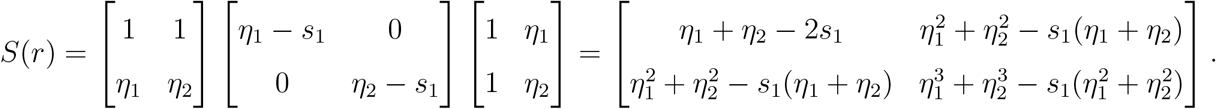

Note that we only have symmetric sums of *η*’s up to the power of 2*n* − 1 = 3 (i.e., 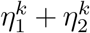 where *k* = 1, 2, 3). Second, we need to assemble the function that counts the number of feasible roots. Thus, to evaluate these symmetric sums, we need to evaluate the Laurent series of the generating function *G*(*N*) = *f* ′(*N*)/*f* (*N*) at *N* = ∞ up to the order *O*(*N* ^−5^) as shown below

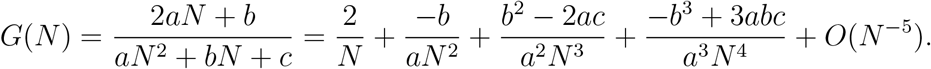

Hence, *η*_1_ + *η*_2_ = −*b/a*, 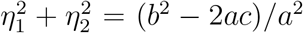, and 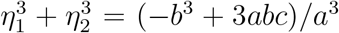. Let us denote these sums by Σ_1_, Σ_2_ and Σ_3_ respectively. Now, the characteristic equation of *S*(*s*_1_) is

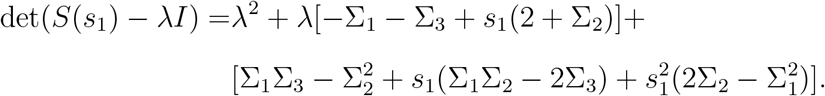

Next, we can construct the characteristic equation whose coefficients are [*v*_2_(*s*_1_) = 1, *v*_1_(*s*_1_), *v*_0_(*s*_1_)] and evaluate the signs of *v*’s at both *s*_1_ = 0 and *s*_1_ = ∞. That is,

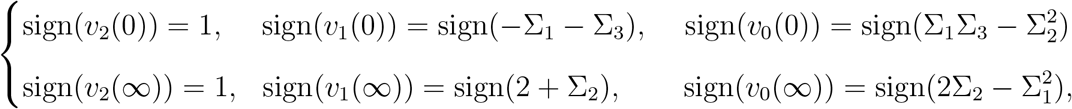

where *v*_*i*_(0) and *v*_*i*_(∞) are the coefficient of the trailing (constant) and leading term of *v*_*i*_(*s*_1_) respectively. Now, we compute *V* (0) and *V* (∞) to have

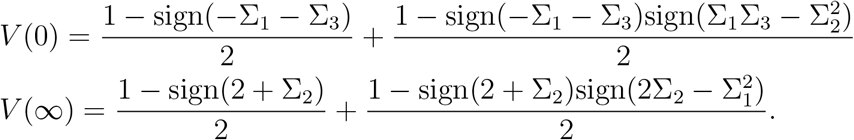

Using the formula *F* (*a, b, c*) = *V* (0) − *V* (∞) together with two basic properties of sign functions (namely sign(*xy*) = sign(*x*)sign(*y*) and sign(*y*) = 1/sign(*y*) for any non-zero real numbers *x* and *y*), we obtain the expression of *F* (*a, b, c*):

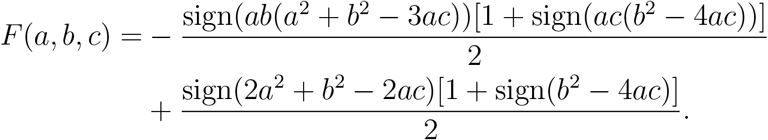

The feasibility basis in this case is given by *v*_0_(0), *v*_1_(0), *v*_0_(∞), *v*_1_(∞). We use the factors shown in the expression of *F* (*a, b, c*) as our basis in the feasibility table. The five quantities that constitute the basis are *Q*_1_ = *ab*, *Q*_2_ = *a*^2^ + *b*^2^ − 3*ac*, *Q*_3_ = *ac*, *Q*_4_ = *b*^2^ − 4*ac*, *Q*_5_ = 2*a*^2^ + *b*^2^ − 2*ac*. Next, we randomize *a, b* and *c* uniformly between −100 to 100 and evaluate the signs of the *Q_i_*’s as well as *F* (*a, b, c*). We find that there are only 3 sign combinations that yield *F* (*a, b, c*) ≥ 1 and are given by the feasibility conditions *C*_1_, *C*_2_ and *C*_3_ shown below

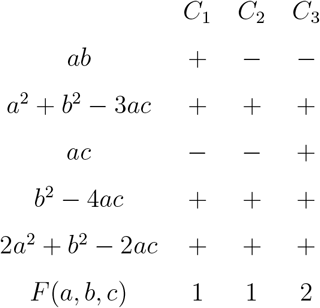

Once the table is obtained, we start the minimization process of the number of feasibility conditions. It is clear from columns 1 and 2 above that the sign of *Q*_1_ does not matter and can be replaced by an X symbol. This concludes the first minimization step as no two columns differ by a single sign and we end up with the feasibility conditions *C*_1+2_ = {*Q*_2_ > 0, *Q*_3_ < 0, *Q*_4_ > 0, *Q*_5_ > 0} and *C*_3_ = {*Q*_1_ < 0, *Q*_2_ > 0, *Q*_3_ > 0, *Q*_4_ > 0, *Q*_5_ > 0}. For the second minimization step, we focus on column *C*_1+2_. We find that the conditional probabilities *P* (*Q*_2_ > 0|*Q*_3_ < 0, *Q*_4_ > 0, *Q*_5_ > 0) = 1, *P* (*Q*_3_ < 0|*Q*_2_ > 0, *Q*_4_ > 0, *Q*_5_ > 0) ≠ 1, *P* (*Q*_4_ > 0|*Q*_2_ > 0, *Q*_3_ < 0, *Q*_5_ > 0) = 1 and *P* (*Q*_5_ > 0|*Q*_2_ > 0, *Q*_3_ < 0, *Q*_4_ > 0) = 1, which implies that the sign of *Q*_2_, *Q*_4_, or *Q*_5_ can be replaced by **X** in that column. Then, let us replace the sign of *Q*_2_ by **X**. Next, we continue computing the conditional properties that were one but without the condition *Q*_2_ > 0. We find that *P* (*Q*_4_ > 0|*Q*_3_ < 0, *Q*_5_ > 0) = 1 and *P* (*Q*_5_ > 0|*Q*_3_ < 0, *Q*_4_ > 0) = 1. This implies that we can replace the sign of *Q*_4_ or *Q*_5_ by **X**. Now, let us replace the sign of *Q*_4_ by **X** and eliminate it from the latter conditional probability to find that *P* (*Q*_5_ > 0|*Q*_3_ < 0) = 1. This time, the sign of *Q*_4_ can be replaced by **X** in column *C*_1+2_. We repeat the same process with the column *C*_3_ and obtain the feasibility table including the 2-step minimization shown below

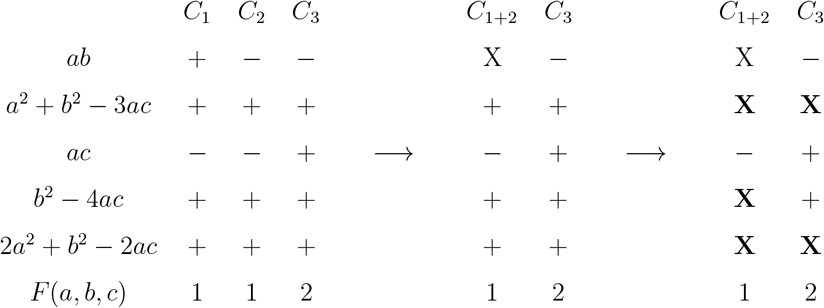

From the last step, we conclude that the condition *ac* < 0 guarantees exactly one feasible equilibrium point (i.e., *F* (*a, b, c*) = 1), while the condition *ab* < 0, *ac* > 0, *b*^2^ − 4*ac* > 0 guarantees exactly 2 feasible equilibrium points (i.e., *F* (*a, b, c*) = 2). Note that a special case of this is the Allee effect model that has the following form (Sun, 2016):

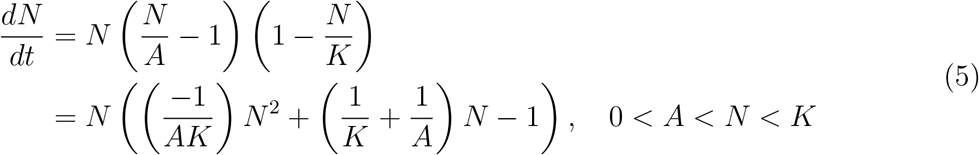

where *a* = −1/*AK*, *b* = 1/*A* + 1/*K* and *c* = −1. It is clear that the second feasibility condition is satisfied as *ab* < 0, *ac* > 0 and *b*^2^ − 4*ac* = (*A* − *K*)^2^/(*A*^2^*K*^2^) > 0 (see Appendix 3 for a minimized feasibility table of a 2-species system with higher-order terms).

## Unfolding links between model parameters

To illustrate additional applications of our methodology, we study the mathematical relationships between model parameters while satisfying feasibility conditions in models that are impossible to solve via isocline approaches. First, let us consider the simplest 2-species LV model with type III functional responses (Turchin, 2003) that is impossible to solve for the location of the equilibrium points analytically.

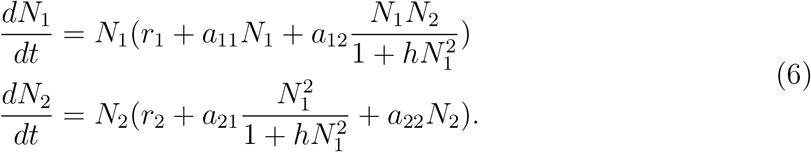

Here, the set of model parameters is given by **Ψ** = (*r*_1_, *r*_2_, *a*_11_, *a*_12_, *a*_21_, *a*_22_, *h*). The common numerators of the RHS of the system above, after deleting *N*_1_ and *N*_2_ outside the brackets, are given by

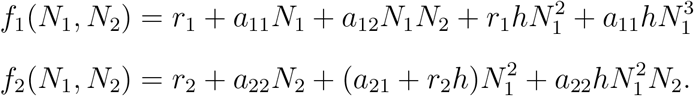

Upon eliminating *N*_1_ from both *f*_1_(*N*_1_, *N*_2_) and *f*_2_(*N*_1_, *N*_2_), we obtain 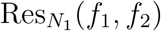 which is a polynomial of degree 5 in *N*_2_ and cannot be solved analytically in closed-form (Abel, 1826). Similarly, upon eliminating *N*_2_ from both *f*_1_(*N*_1_, *N*_2_) and *f*_2_(*N*_1_, *N*_2_), we obtain 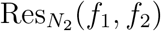 that is a polynomial of degree 5 in *N*_1_ as shown below:

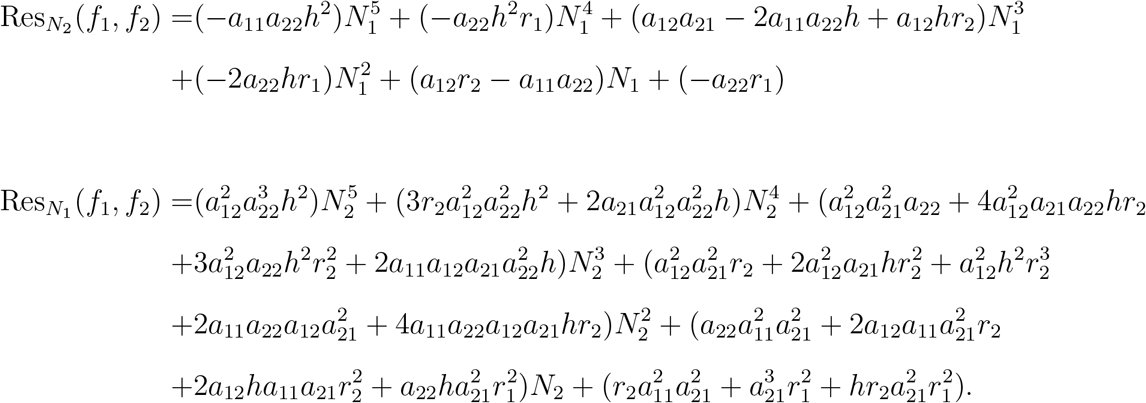

In other words, the number of roots of *f*_1_(*N*_1_, *N*_2_) and *f*_2_(*N*_1_, *N*_2_) is 5. Note that the roots of the univariate polynomials 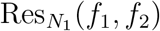 and 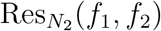, upon appropriate pairing of roots of the first polynomial with the second, are the roots of the system *f*_1_(*N*_1_, *N*_2_) = 0 and *f*_2_(*N*_1_, *N*_2_) = Since the roots of either 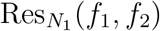 or 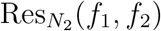 are unattainable analytically, then the system *f*_1_(*N*_1_, *N*_2_) = 0 and *f*_2_(*N*_1_, *N*_2_) = 0 is unsolvable analytically.

Next, to find relationships between model parameters, for illustration purposes, let us consider the parameters **Ψ** = (*r*_1_, *r*_2_, *a*_11_, *a*_12_, *a*_22_) = (0.5, −1.5, 1, −1.5, 1), and the parameters *a*_21_ ∈ [−6, −1] and *h* ∈ [0.5, 4]. In this special case, we find that feasibility (i.e., *F* (**Ψ**) ≥ 1) can only be satisfied under the single condition *v*_0_(0, 0) < 0. See Appendix 4 for the expression of *v*_0_(0, 0) written as symmetric sums (i.e., sigmas), along with closed forms of the sigmas and derivations. We find that the feasibility domain generated via solving numerically (using the software tool PHCLab) the isocline equations (i.e., *f*_1_(*N*_1_, *N*_2_) = 0, *f*_2_(*N*_1_, *N*_2_) = 0) and checking for the feasibility of roots matches the domain generated by the inequality *v*_0_(0, 0) < 0 (see Fig. 1A-B). Note that *a*_21_ and *h* are independent in the model, yet they are bounded by feasibility.

**Figure 1:**
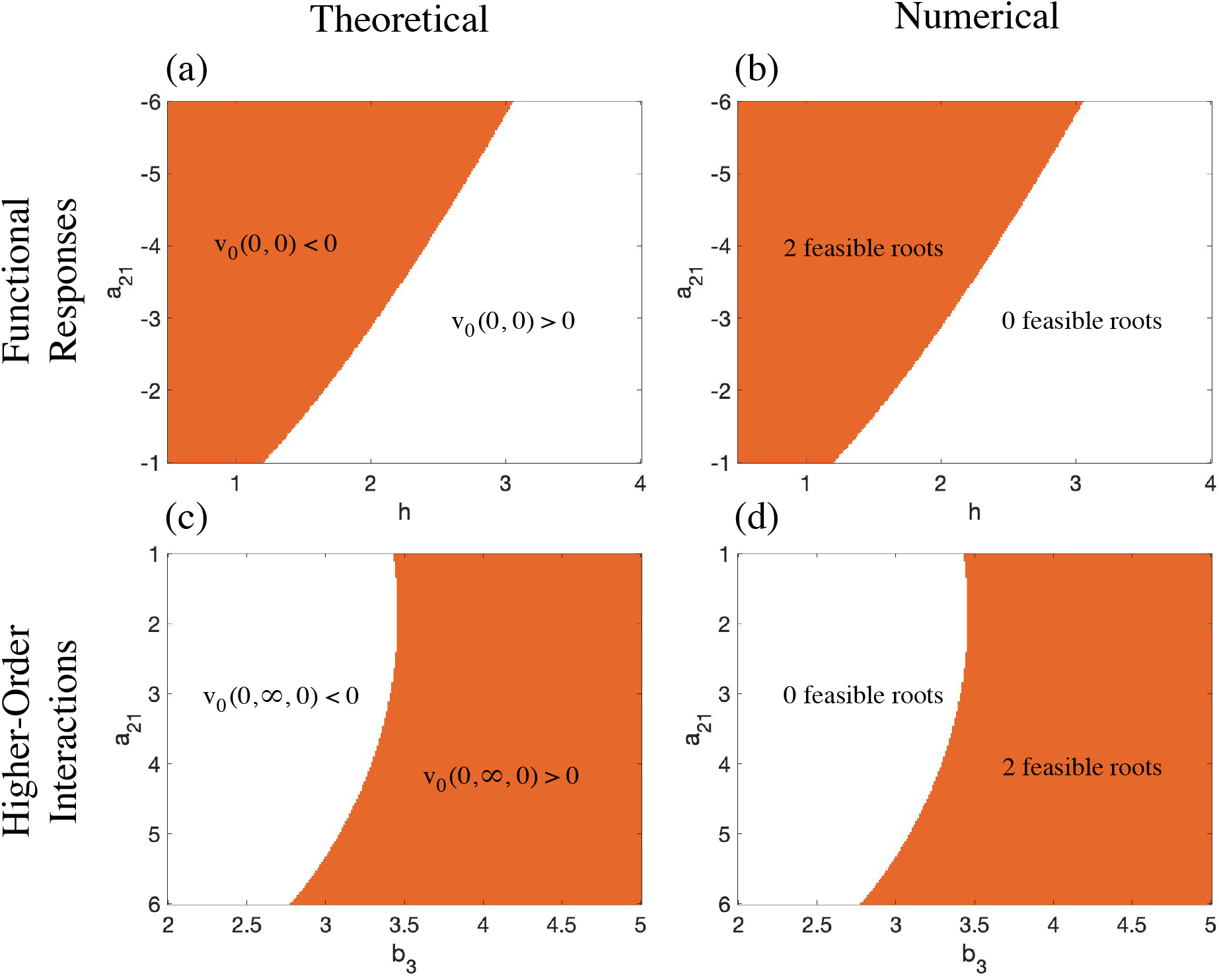
Unfolding mathematical links between model parameters. Panels A-B show the mathematical link between a pairwise interaction *a*_21_ and the constant *h* while maintaining feasibility in a modified Lotka-Volterra model with type III functional responses (see main text), where (*r*_1_, *r*_2_, *a*_11_, *a*_12_, *a*_22_) = (0.5, −1.5, 1, −1.5, 1), *a*_21_ ∈ [−6, −1] and *h* ∈ [0.5, 4]. The panels show the sign of *v*_0_(0, 0) and the number of feasible roots. Note that *v*_0_(0, 0) is the constant or trailing term of the characteristic equation (i.e., coefficient of *λ*^0^) evaluated at *s*_1_ = 0 and *s*_2_ = 0 (the *s*’s are the variables in the symmetric matrix *S = W* Δ*W* ^*t*^, see Methodology). The number of feasible roots is obtained by solving the isocline equations numerically using the software package PHCLab and checking for the feasibility of roots. Both panels confirm that the number of feasible roots is greater than zero when *v*_0_(0, 0) < 0. Hence, the theoretical relationship is given by *F* (**Ψ**) = −2∗sign(*v*_0_(0, 0)). Panels C-D show the mathematical link between a pairwise interaction *a*_21_ and a higher-order interaction *b*_3_ while maintaining feasibility in a modified Lotka-Volterra model with higher-order interactions (see main text), where (*r*_1_, *r*_2_, *r*_3_, *a*_11_, *a*_12_, *a*_13_, *a*_22_, *a*_23_, *a*_31_, *a*_32_, *a*_33_, *b*_1_, *b*_2_) = (1.5, −1.5, −1.5, 2, −1.5, −1.5, 2, −1.5, −1.5, −1, 1, 1, −1), *a*_21_ ∈ [1, 6] and *b*_3_ ∈ [2, 5]. The panels show the sign of *v*_0_(0, ∞, 0) and the number of feasible roots. Note that *v*_0_(0, ∞, 0) is the coefficient of the highest power in *s*_2_ in the trailing term of the characteristic equation (see Methodology). Again, the number of feasible roots is obtained by solving the isocline equations numerically using the software package PHCLab and checking for the feasibility of roots. Both panels confirm that the number of feasible roots is positive when *v*_0_(0, ∞, 0) > 0. Hence, the theoretical relationship is given by *F* (**Ψ**) = 2∗sign(*v*_0_(0, ∞, 0)).

As a second example, let us consider the LV model with higher-order interactions that is shown below. This example is the simplest ecological 3-species model whose isocline equations are impossible to be solved analytically as it has five roots (see Appendix 6).

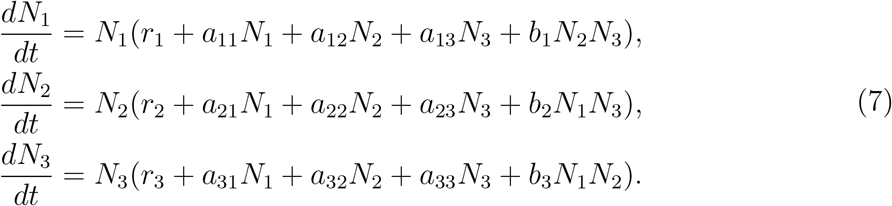

To study the feasibility conditions of this model, we need to consider the three polynomials inside the brackets. The resultants are shown in Appendix 6. Let us consider

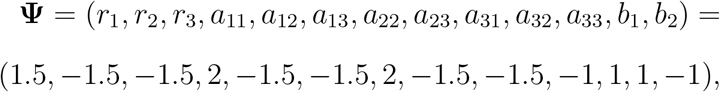

where the parameters *a*_21_ ∈ [1, 6] (pairwise effect of species 1 on 2) and *b*_3_ ∈ [2, 5] (higher-order effect on species 3) are restricted. We find that feasibility (i.e., *F* (**Ψ**) ≥ 1) is satisfied when *v*_0_(0, ∞, 0) > 0 (see Appendix 6 for more details). Again, for confirmation purposes, the feasibility domain generated by solving numerically the isocline equations (i.e., *f*_*i*_(*N*_1_, *N*_2_, *N*_3_) = 0 for *i* = 1, 2, 3) using the software tool PHCLab and checking for the feasibility of roots matches the domain generated by the inequality *v*_0_(0, ∞, 0) > 0 (see Fig. 1C-D). This illustrates that pairwise and higher-order interactions can be non-trivially linked and their incorporation into ecological models must be done with caution.

## Discussion

Feasibility conditions can be obtained analytically by solving the isocline equations for species abundances 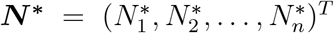 before imposing the positivity condition ***N***\* > **0**. This approach works well for LV model, whose isocline equations is the linear system ***r*** + ***AN***\* = **0** and whose feasibility conditions are given by ***N***\* = ***−A***^−1^***r*** > **0** (Goh, 1976; Volterra and Brelot, 1931; Saavedra et al., 2020). However, when the isocline equations have five or more complex roots, the system of polynomial equations cannot be solved analytically. This is a consequence of Grobner elimination theorem combined with Abel’s impossibility theorem (Adams et al., 1994; Abel, 1824, 1826). Specifically, from the elimination theorem, in any system of polynomial equations which has Θ complex roots and *n* variables, any *n* − 1 variables can be eliminated from the system to obtain a univariate polynomial with the remaining variable of degree at least Θ. The roots of this univariate polynomial are all the correspondent coordinates of the roots of the isocline equations (Adams et al., 1994). This is a generalization of Gaussian elimination, which can eliminate any *n* − 1 variables from the system leaving a single linear univariate polynomial in the remaining variable to be solved (Lazard, 1983). However, from Abel’s impossibility theorem, it is impossible to solve a univariate polynomial in terms of radicals (i.e., analytically) (Abel, 1824, 1826) if this polynomial has five or more roots. For instance, this number of roots is quickly reached by adding Type III functional responses to a 2-species LV model or adding higher-order interactions to a 3-species LV model (AlAdwani and Saavedra, 2019).

In this work, we have proposed a general formalism to analytically obtain the feasibility conditions for any multivariate, polynomial, population, dynamics model of any dimensions without the need to solve for the equilibrium locations. We found that feasibility conditions are entirely functions of symmetric sums of the roots of the isocline equations. Unlike the location of the roots, which cannot be obtained analytically, symmetric sums of the roots can be obtained for any polynomial system regardless of order and dimension. We have also created an analytical formula of the number of feasible roots in the system, which are functions of signs of Θ2^*n*^ quantities (i.e., the *v*’s evaluated at the feasibility box whose coordinates compose of zeros and infinities). We have shown how to create a feasibility table (i.e., matrix) whose columns are the individual feasibility conditions of the model. We have then provided a minimization process that can combine feasibility conditions into fewer ones and remove redundant quantities. Of course, the expressions involved in the inequality are complicated, nevertheless, they can be significantly simplified by sophisticated factorization.

Additionally, we have shown how to provide feasibility conditions under parameter restrictions. We have shown that by restricting parameters, the feasibility domain can be described by a single inequality only. In recent years, the topic of feasibility has been focused on relationships between parameters while maintaining feasibility (Saavedra et al., 2017). Using simulations (i.e., solving for the location of the isocline equations numerically then checking for the feasibility of roots) one can plot the feasibility domain for one, two, or three parameters at most while fixing the remaining ones. However, it is impossible to generate a four-dimensional plot that the human eye can capture. Also, it is impossible to find an analytical expression of the feasibility domain using numerical simulations. Of course, someone can find an approximate formula of the feasibility domain, nevertheless, there is no unique formula and different approximations may lead to different interpretations of how parameters are linked while maintaining feasibility. Following our proposed methodology, we can determine mathematically how any number of parameters are linked by describing polynomial inequalities that are functions of those free-parameters while maintaining feasibility: a task that is impossible to perform with simulations. This is an important property to consider in ecological modeling given that mathematical expressions are frequently formed assuming that parameters are independent of each other. However, once one imposes mechanisms or constraints, such as feasibility, these parameters can be linked and break the conclusions based on independent parameters (Song et al., 2019).

Our methodology provides a fast method for plotting feasibility domains, computing the number of feasible roots, and displaying feasibility conditions. For example, for our 3-species example with higher-order interactions, plotting the feasibility domain by solving the isocline equations numerically using the software package PHCLab (Guan and Verschelde, 2008) took more than 1.5 hours to compute the number of feasible points with 2^16^ trials. Instead, using our methodology (and code which involves a naive implementation of our methodology without parallelization), it took less than 11.5 minutes to run the analysis, and a few seconds to plot the feasibility domain for different ranges of the free parameters using the same number of trials. Moreover, when we change the ranges of our free parameters *a*_21_ and *b*_3_, we only need a few seconds to run our code, whereas we need to repeat the entire 1.5 hours with the traditional numerical technique. With a clever implementation of the methodology and parallelizing the code (since the entire methodology can be parallelized), a faster computation of the feasibility domain/conditions and links between parameters can be achieved.

One significant drawback of the methodology is that it requires the handling of large symbolic expressions. Thus, careful implementation is required to run a successful code using the presented methodology. For example, when we created the generating function *G*, we did not multiply the determinant of the eliminant *T* and the determinant of the Jacobian of the isocline equations *J*, divided them by the product of all resultants, and took the series expansion of the final polynomial quotient. Instead, we took the series expansion of each resultant reciprocal separately, wrote *TJ* as multivariate polynomial in species abundances, found the coefficients of each term, and multiplied it by a single appropriate term in the series expansion of each resultant reciprocal to find the Σ’s. However, it is always possible to handle such large expressions as the entire methodology can be parallelized. The second drawback of the methodology is its susceptibility to numerical errors. In our 3-species application example, our code gives as output non-integer values of the number of feasible roots in the system. Nevertheless, in our example we rectified it quickly by assigning non-integer values to their closest integers (see Appendix 6). Remember that the methodology requires only checking signs of large symbolic expressions, and we do not need them to be computed accurately. Nevertheless, such quantities can be computed more accurately by following several techniques such as increasing precision of numeric calculations. Similarly, cancellation errors can be reduced by combining positive numbers and negative ones together, and then performing a single subtraction. Round-off and truncation errors can also be avoided when ratios are computed. For example, instead of computing (10^90^ − 10^91^)/10^90^ by computing (10^90^ − 10^91^) then dividing the result by 10^90^, it is better to add 10^90^/10^90^ = 1 with −10^91^/10^90^ = −10 as the latter reduces round-off errors in large computations (Trefethen and Bau III, 1997). Of course, there are other techniques to reduce such errors, nevertheless, it is important to think about numerical errors in the implementation process.

In sum, the contribution of this theoretical work is that it provides a foundation for important ecological concepts such as species coexistence, stability, and permanence. Indeed, it has been shown that the existence of a feasible solution is a necessary condition for persistence and permanence in dynamical models of the form *dN*_*i*_/*dt* = *N*_*i*_*f*_*i*_(***N***)/*q*_*i*_(***N***) (Hofbauer and Sigmund, 1998; Stadler and Happel, 1993). Similarly, it has been proved that this type of models cannot have bounded orbits in the feasibility domain without a feasible free-equilibrium point (Hofbauer and Sigmund, 1998). In fact, we cannot talk about asymptotic or local stability without the existence of a feasible equilibrium point (AlAdwani and Saavedra, 2020). Hence, coexistence, stability, or permanence domains are subsets of the feasibility domain and their conditions are effectively the feasibility conditions obtained in this work plus some added conditions. Thus, this work unlocks the opportunity to increase our systematic understanding of multispecies coexistence.

## Acknowledgments

MA would like to thank Kuwait Foundation for the Advancement of Sciences (KFAS) and Kuwait Chamber of Commerce and Industry (KCCI) for their support. Also, MA would like to thank MathWorks Inc. for their fellowship. SS acknowledges funding by NSF grant No. DEB-2024349.

## Data sharing

The code supporting the results can be found at https://github.com/MITEcology/Feasibility_AlAdwani_2021.

## Appendix

## S1 Methodology: One-Dimensional Systems (1 species)

In this section, we focus on univariate polynomial systems. The aim here is to find a closed form expression for the number of feasible roots in the system *F* (**Ψ**), which is a function of model parameters **Ψ**. From the expression of *F* and exploiting its property, we can deduce feasibility conditions which are sets of polynomial inequalities. For this section, let us consider the following polynomial dynamical system of a single variable *N* as shown below

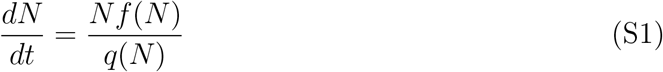

where *f* (*N*) is a polynomial of degree *n* whose coefficients are in **Ψ**. We already know that the number of roots of *f* (*N*) is *n*, a consequence of the Fundamental Theorem of Algebra. In this section, we derive the formula of *F* (**Ψ**) and derive feasibility conditions from it. The procedure involves the following steps:

1. Consider the monomial map *m*(*N*) = [1, *N, N*^2^, …, *N*^*n*−1^]^*T*^ which is of length *n* and let *Q*(*N*) = *N*. Next, denote the roots of *f* (*N*) by *η*_1_, *η*_2_, …, *η*_*n*_ then denote the symmetric sums of the roots 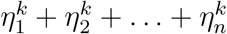 by Σ_*k*_ for *k* = 0, 1, 2, ….
2. Construct the symmetric matrix *S*(*s*_1_) = *W* Δ*W*^*t*^ where *W*_*ij*_ = *m*_*i*_(*η*_*j*_) and Δ_*ii*_ = *Q*(*η*_*i*_ − *s*_1_) is a diagonal matrix. Note that all entries of *S*(*s*_1_) contains only symmetric sums of the *η*’s (i.e, *S*(*s*_1_) only contains *s*_1_’s and Σ’s).
3. Construct the generating function *G*(*N*) = *f* ′(*N*)/*f* (*N*) and evaluate the Laurent series of *G*(*N*) at *N* = ∞. The purpose of the series is to evaluate the Σ’s from looking at the coefficients of the Laurent series of *G*(*N*), which are functions of model parameters **Ψ** (or coefficients of *f* (*N*)). Assuming that 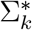 is the highest symmetric sum that is needed to be evaluated, the following identity is valid:

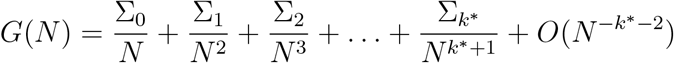
4. After evaluating the Laurent series up to order 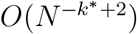, *S*(*s*_1_) is a function of *s*_1_ and model parameters only, evaluate the characteristic polynomial of *S*(*s*_1_) and write it in the form det(*S*(*s*_1_) − *λI*) = (−1)^*n*^λ^*n*^ + *v*_*n*−1_(*s*_1_)*λ*^*n*−1^ + … + *v*_0_(*s*_1_). After that consider the sequence **v** = [*v*_*n*_(*s*_1_) = (−1)^*n*^, *v*_*n*−1_(*s*_1_), …, *v*_0_(*s*_1_)] and let *V* (*s*_1_) be the number of consecutive sign changes in **v**.
5. Define the function sign(*x*) to be 1 when *x* > 0, 0 when *x* = 0 and −1 when *x* < 0. Before writing down the expression of *V* (*s*_1_), note that in order to determine whether there is a sign change between two real numbers *x* and *y*, we simply evaluate [1 − sign(*xy*)]/2, which is 0 when *x* and *y* have the same sign and 1 otherwise. With this expression, the formula of *V* (*s*_1_) is

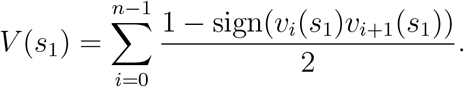
6. For any interval (*a, b*], the number of real roots of *f* (*N*) in (*a, b*] is exactly *V* (*a*)−*V* (*b*). Hence, to obtain the analytical expression for *F* (**Ψ**), we consider the interval (0, ∞) to obtain *F* (**Ψ**) = *V*(0) − *V*(∞) or simply

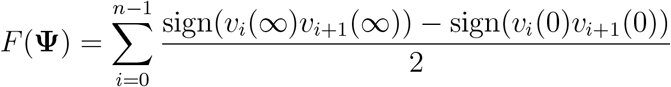
7. Call *v*_0_(0), …, *v*_*n*−1_(0), *v*_0_(∞), …, *v*_*n*−1_(∞) the feasibility basis. Since each of the *v*_*i*_’s can take a positive or a negative sign, then there are 2^2*n*^ sign combinations the feasibility basis can take. Many of those combinations are impossible to occur (empty) for any choice of real **Ψ**. To detect the non-empty sign combinations, we compute the signs of all *v*_*i*_’s as well as *F* (**Ψ**) for a range of parameters **Ψ**, where each component of **Ψ** varies independently in a large domain (say uniformly between −100 and 100). This operation is cheaply computed as it is evaluation a few functions and not solving systems of equations. After that, we extract unique sign combinations of the *v*_*i*_’s which yield *F* (**Ψ**) ≥ 1 and put them in a feasibility table whose rows are the signs of the *v*_*i*_’s and columns are the individual feasibility conditions. For a cleaner representation of feasibility conditions, we can investigate all sign combinations of all the factors of each of the *v*_*i*_’s (feasibility basis) and deleting all perfect square factors from them (if possible).
8. After we obtain the table, we perform minimization to it by combining the feasibility conditions (the columns). If two columns with the same value of *F* (**Ψ**) differ by a single sign (in one row), combine the two columns into one and place X in the row where there is a single sign difference to indicate that that no condition is needed to be imposed for the quantity associated with that row. We can combine columns with different values of *F* (**Ψ**) if the user does not care about separating the conditions based on the value of *F*. Then we iterate through the process until it terminates (no two columns differ by a single sign). For further minimization, we eliminate redundant signs where the sign of one or more quantities that constitute the basis implies the sign of another quantity in the same basis. For example, if the quantities *ac* and *a*^2^ + *b*^2^ − 3*ac* are in the basis, then *ac* < 0 implies *a*^2^ + *b*^2^ − 3*ac* > 0 making the later inequality redundant. Sometimes, the quantity in the basis is always positive or negative regardless of the sign of the others (e.g *a*^2^ + *b*^2^ > 0 is always true). To find these cases, we go through a single column at a time and iterate through each quantity in the basis then compute the conditional probability that the quantity in the basis takes its correspondent sign given that all other remaining quantities in the same basis have their correspondent signs. If one or more conditional probabilities are 1, any of those quantities may be replaced by **X** in the table. We then repeat computing the same conditional probabilities which were 1 but without the **X**’ed quantity being part of the calculation. Then, we check whether the conditional probabilities are still 1 or not. If any is 1, we delete a redundant sign and keep repeating the process until no conditional probability is 1. We then go through all columns and repeat the same process until it terminates

## S2 Methodology: Two-Dimensional Systems (2 species)

Let us consider the following dynamical system with two species as shown below

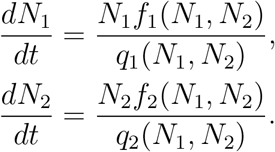

where *f*_1_(*N*_1_, *N*_2_) and *f*_2_(*N*_1_, *N*_2_) are multivariate polynomial in *N*_1_ and *N*_2_ and whose coefficients are in the vector **Ψ**. To describe the feasibility domain analytically, the following steps are followed:

1. Let *d*_1_ and *d*_2_ equal to the largest exponent of *N*_1_ in *f*_1_ and *f*_2_ respectively. Write 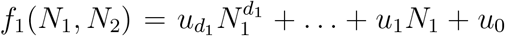 and 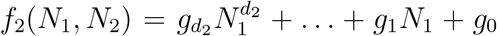 where the *u*’s and *g*’s are functions of *N*_2_ and are not functions of *N*_1_. Next, find *T*_21_ and *T*_22_ such that the resultant 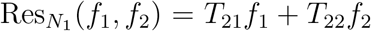 where 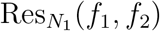 is a determinant of a square matrix of dimension *d*_1_ + *d*_2_ as shown below.

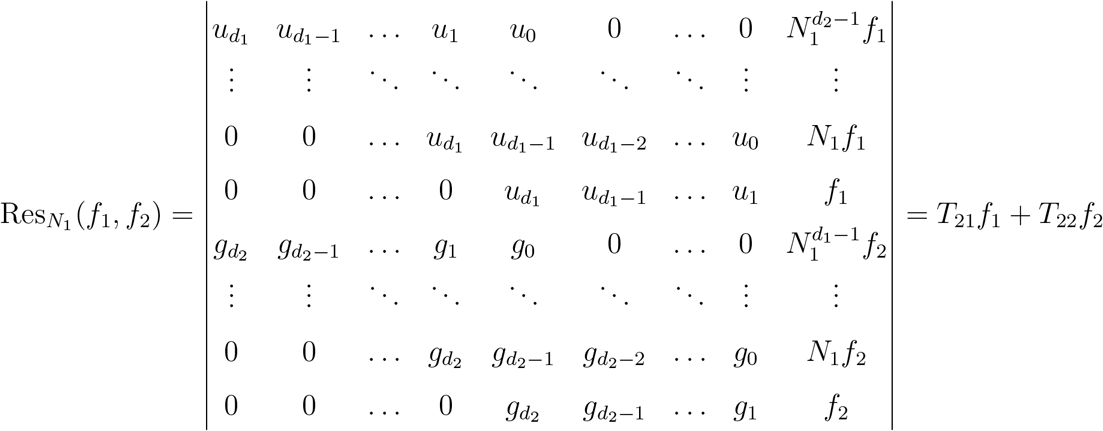

Note that in the rows where the last entry is *f*_1_ or *f*_2_, there are no *u*_0_ nor *g*_0_ there. To form 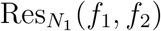, it is better to start with the two rows whose the last entries are *f*_1_ and *f*_2_ then construct the matrix up. Now, if 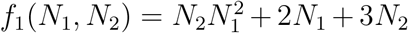 and *f*_2_(*N*_1_, *N*_2_) = 4*N*_2_*N*_1_ + 5, then *d*_1_ = 2, *d*_2_ = 1 and 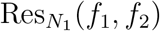 is a determinant of a 3 by 3 matrix as shown below:

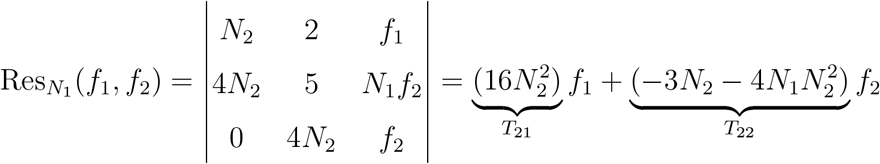
2. Let *ϵ*_1_ and *ϵ*_2_ equal to the largest exponent of *N*_2_ in *f*_1_ and *f*_2_ respectively. Write 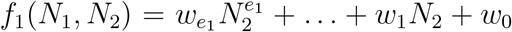 and 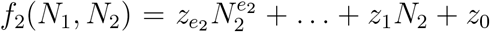 where the *w*’s and *z*’s are functions of *N*_1_ and are not functions of *N*_2_. Next, find *T*_11_ and *T*_12_ such that the resultant 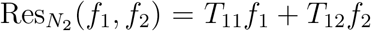 where 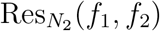 is a determinant of a square matrix of dimension *e*_1_ + *e*_2_ as shown below.

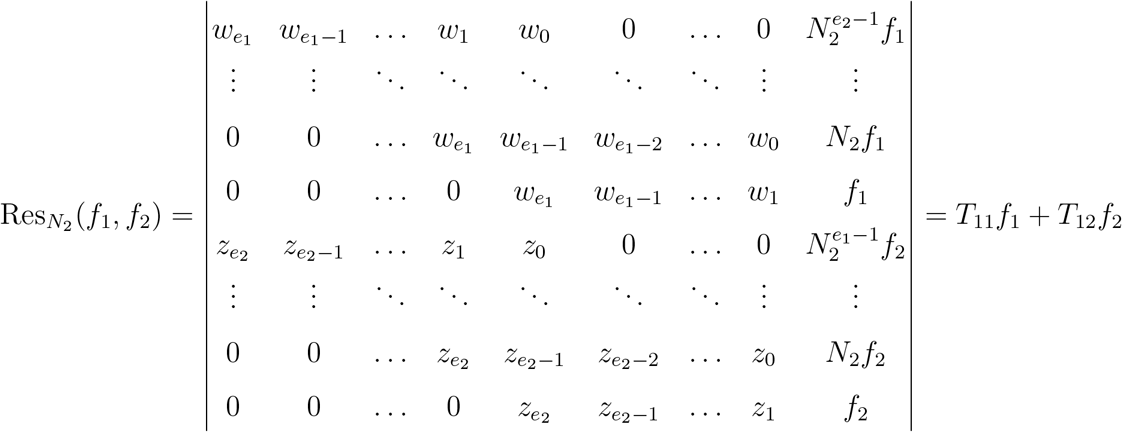

Note that in the rows where the last entry is *f*_1_ or *f*_2_, there are no *w*_0_ or *z*_0_ there. Again, if 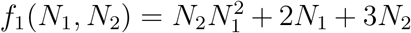 and *f*_2_(*N*_1_, *N*_2_) = 4*N*_2_*N*_1_ + 5, then *e*_1_ = 1, *e*_2_ = 1 and 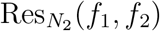 is a determinant of a 2 by 2 matrix as shown below:

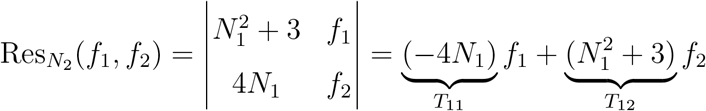
3. Evaluate the determinant of the eliminating matrix *T* (*f*_1_, *f*_2_), whose elements *T*_11_, *T*_12_, *T*_21_, *T*_22_ have been obtained in the earlier two steps, as well as the determinant of the Jacobian of *f*_1_ and *f*_2_. Note that the first row of *T* (*f*_1_, *f*_2_) corresponds to the coefficients of *f*_1_ and *f*_2_ in 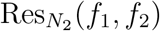 while the second row corresponds to the coefficients of *f*_1_ and *f*_2_ in 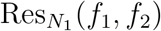.

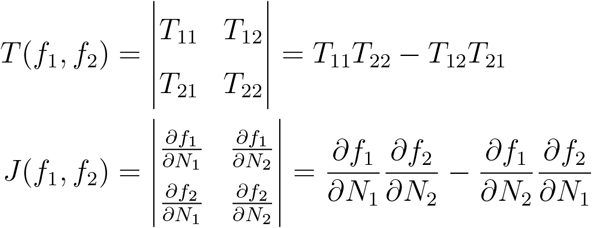
4. Expand the function *G*(*f*_1_(*N*_1_, *N*_2_), *f*_2_(*N*_1_, *N*_2_)) that is shown below, around *N*_1_ = ∞ and *N*_2_ = ∞ (or perform series expansion of *G*(*f*_1_(1/*x*, 1/*y*), *f*_2_(1/*x*, 1/*y*)) around *x* = 0 and *y* = 0 which gives identical coefficients) to obtain the Σ’s (symmetric sums of the roots).

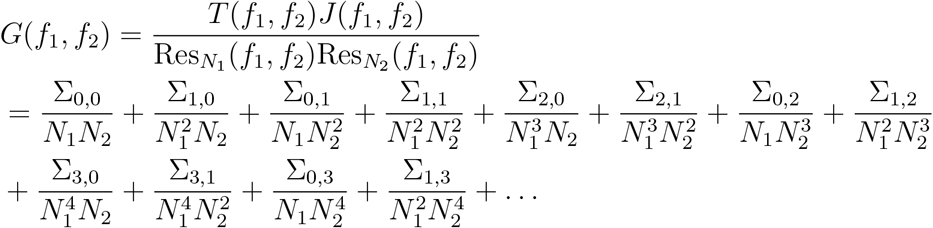

Note that *f*_1_ and *f*_2_ are substituted in both 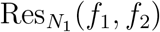 and 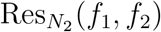 to fully express the resultants in terms of *N*_1_, *N*_2_ and model parameters **Ψ** before evaluating *G*(*f*_1_, *f*_2_) and expanding it. For the symmetric sums, denote the roots of *f*_1_(*N*_1_, *N*_2_) and *f*_2_(*N*_1_, *N*_2_) by ***η***_***k***_ = [*η*_***k***,1_, *η*_***k***,2_]^*T*^ for *k* = 1, …, Θ. The symmetric sum ∑_*m,n*_ for any *m* and *n* is given by 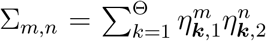. In particular, note that Θ = ∑_0,0_ is the number of complex roots of *f*_1_ and *f*_2_ with a general coefficients. It is important to record that number.
5. Choose a map *m*(*N*_1_, *N*_2_) = [1, *m*_1_, *m*_2_, …, *m*_Θ−1_]^*T*^ of length Θ with independent entries that are functions of *N*_1_ and *N*_2_. If Θ = 4, we can let *m*(*N*_1_, *N*_2_) = [1, *N*_1_, *N*_2_, *N*_1_*N*_2_]^*T*^. It does not matter what the *m*’s are as long as no entry is a linear combination of the others and step 7 of the procedure does not fail (in step 7 we give more details). Then let *Q*(*N*_1_, *N*_2_) = *N*_1_*N*_2_ and compute the symmetric matrix *S*(*s*_1_, *s*_2_) = *W* Δ*W* ^*t*^ where *W*_*ij*_ = *m*_*i*_(*η*_***j***,1_, *η*_***j***,2_) and Δ_*ii*_ = *Q*(*η*_***i***,1_ − *s*_1_, *η*_***i***,2_ − *s*_2_) is a diagonal matrix.
6. The next task is to evaluate the determinant of *S*(*s*_1_, *s*_2_) and write it in the form det(*S*(*s*_1_, *s*_2_)−*λI*) = (−1)^Θ^*λ*^Θ^+*v*_Θ−1_(*s*_1_, *s*_2_)*λ*^Θ−1^ +*…*+*v*_0_(*s*_1_, *s*_2_). After that consider the sequence **v** = [*v*_Θ_(*s*_1_, *s*_2_) = (−1)^Θ^, *v*_Θ−1_(*s*_1_, *s*_2_), …, *v*_0_(*s*_1_, *s*_2_)] and let *V* (*s*_1_, *s*_2_) be the number of consecutive sign changes in **v**. The formula of *V* (*s*_1_, *s*_2_) is

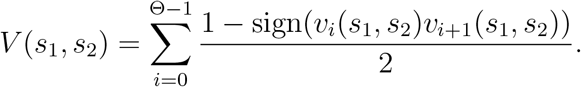
7. For any interval (*a, b*] × (*c, d*], the number of real roots of *f*_1_(*N*_1_, *N*_2_) and *f*_2_(*N*_1_, *N*_2_) in (*a, b*] × (*c, d*] is exactly [*V* (*a, c*) − *V* (*a, b*) + *V* (*b, d*) − *V* (*b, c*)]/2. For the feasibility domain, note that the points {(0, 0), (0, ∞), (∞, 0), (∞, ∞)} are the vertices of the “box” that bound it. Hence, the expression of *F* (**Ψ**) is simply *F* (**Ψ**) = [*V* (0, 0) − *V* (0, ∞) − *V* (∞, 0) + *V* (∞, ∞)]/2. Here, ∞ is a limit; therefore, 0 is evaluated first before the limit is taken at infinity. For *V* (∞, ∞) where the limit of two quantities approach infinity, the limit is unique. Therefore, it does not matter along which direction the limit is taken. To evaluate those *V*’s we need to evaluate the *v*’s at those limits. Note that for any function *p*, sign(*p*(*S*(0, 0)) = sign of the constant term in *p*(*S*(*s*_1_, *s*_2_)). For the other cases, note that

- sign(*p*(*S*(0, ∞)) = sign of the coefficient of the term associated with the highest power of *s*_2_ in *p*(*S*(*s*_1_, *s*_2_)) = sign of the constant term of the common numerator of *p*(*S*(0, 1/*y*)) = sign of the numerator of *p*(*S*(0, 1/*y*)) evaluated at *y* = 0
- sign(*p*(*S*(∞, 0)) = sign of the coefficient of the term associated with the highest power of *s*_1_ in *p*(*S*(*s*_1_, *s*_2_)) = sign of the constant term of the common numerator of *p*(*S*(1/*x*, 0)) = sign of the numerator of *p*(*S*(1/*x*, 0)) evaluated at *x* = 0
- sign(*p*(*S*(∞, ∞)) = sign of the coefficient of the term associated with the highest power of *s*_1_*s*_2_ in *p*(*S*(*s*_1_, *s*_2_)) = sign of the constant term of the common numerator of *p*(*S*(1/*x*, 1/*y*)) = sign of the numerator of *p*(*S*(1/*x*, 1/*y*)) evaluated at *x, y* = 0

After evaluating the *v*’s, we assemble *F* (**Ψ**). If *F* (**Ψ**) or any of the *V*’s is not a non-negative integer, even for a single case where **Ψ** is randomly chosen, or the vector **v** contains zeros, then the map *m*(*N*_1_, *N*_2_) must be changed. One remedy to rectify this is increasing the order of one of the components of *m*. For example, if *m*(*N*_1_, *N*_2_) = [1, *N*_1_, *N*_2_, *N*_1_*N*_2_] fails, then one can try *m*(*N*_1_, *N*_2_) = [1, 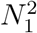, *N*_2_, *N*_1_*N*_2_].
8. As in the univariate case, call *v*_0_(0, 0), *…*, *v*_Θ−1_(0, 0), *v*_0_(0, ∞), *…*, *v*_Θ−1_(0, ∞), *v*_0_(∞, ∞), *…*, *v*_Θ−1_(∞, ∞), *v*_0_(∞, 0), *…*, *v*_Θ−1_(∞, 0) the feasibility basis which involves 4Θ quantities as feasibility conditions are only dependent on those quantities. Since there are 4Θ quantities and each can take a positive or a negative sign, then there are 2^4Θ^ sign combinations. Many of those combinations are impossible to occur (empty) for any choice of real **Ψ**. To detect the non-empty sign combinations, we compute the signs of all the *c*’s (the feasibility basis) as well as *F* (**Ψ**) for a range of parameters **Ψ**, where each component of **Ψ** varies independently in a large domain (say uniformly between −100 and 100 or in any suitable domain). This operation is cheaply computed as it is evaluation a few functions and not solving systems of equations. After that, we extract unique sign combinations of the *v*’s which yield *F* (**Ψ**) ≥ 1 and put them in a feasibility table whose rows are the signs of the *c*’s and columns are the individual feasibility conditions.
9. After we obtain the feasibility table, we perform minimization to it. If two columns differ by a single sign (in one row), the two columns are combined into one and an X is placed in the row where there is a single sign difference. We repeat the same process until no two columns differ by a single sign. After that we go through a single column at a time and iterate through each quantity in the basis then compute the conditional probabilities that the quantity takes its correspondent sign given that all remaining quantities have their correspondent signs. If one or more conditional probabilities are 1, the sign of one of those quantities may be replaced by **X** in the table. We then repeat computing the same conditional probabilities which were 1 but without the **X**’ed quantity being part of the calculation. If any conditional probability is 1 we repeat the process until it terminates. Plotting the signs of the feasibility basis against *F* (**Ψ**) may reveal extra minimization information (see application section).

## S3 Ex 1: 2-Species with Simple Higher-Order Terms

Consider the following LV system with a simple higher-order term *N*_1_*N*_2_

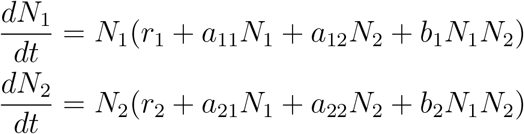

Let *f*_1_(*N*_1_, *N*_2_) = *r*_1_ +*a*_11_*N*_1_ +*a*_12_*N*_2_ +*b*_1_*N*_1_*N*_2_ and *f*_2_(*N*_1_, *N*_2_) = *r*_2_ +*a*_21_*N*_1_ +*a*_22_*N*_2_ +*b*_2_*N*_1_*N*_2_ with **Ψ** = (*r*_1_, *r*_2_, *a*_11_, *a*_12_, *a*_21_, *a*_22_, *b*_1_, *b*_2_) being the vector of model parameters. The first task is evaluating the resultants as follows:

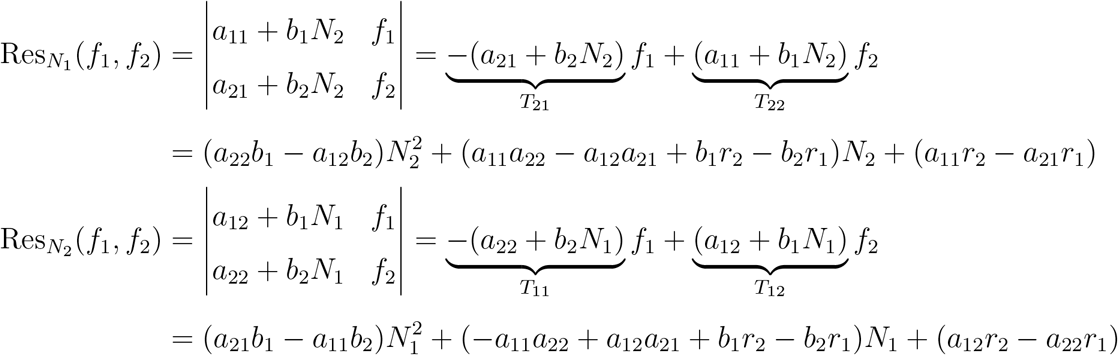

From above, the entries of the eliminating matrix *T* (*f*_1_, *f*_2_) are *T*_11_ = −(*a*_22_ + *b*_2_*N*_1_), *T*_12_ = (*a*_12_ + *b*_1_*N*_1_), *T*_21_ = −(*a*_21_ + *b*_2_*N*_2_) and *T*_22_ = (*a*_11_ + *b*_1_*N*_2_). After that we evaluate the determinant of both eliminating matrix *T* (*f*_1_, *f*_2_) and the Jacobian of *f*_1_ and *f*_2_:

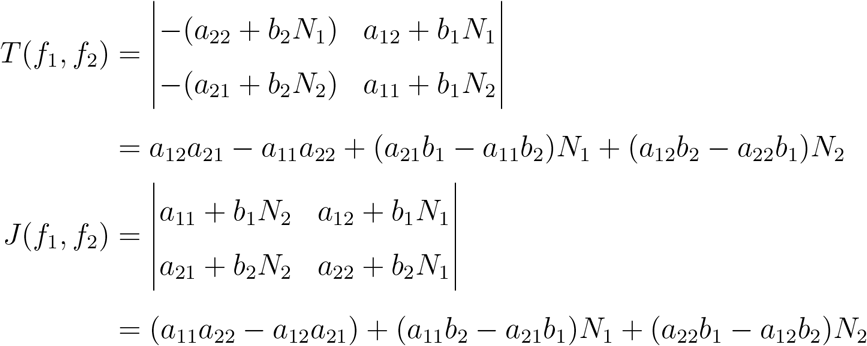

Expand the generating function *G*(*f*_1_(*N*_1_, *N*_2_), *f*_2_(*N*_1_, *N*_2_)) around *N*_1_, *N*_2_ = ∞ to obtain

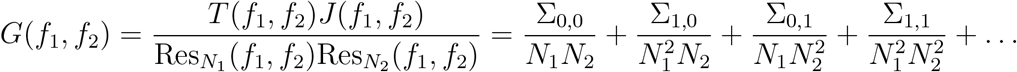

The expression for each of the ∑’s (symmetric power sums of the roots) are shown below where 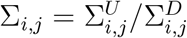 is written as a fraction of two polynomials.

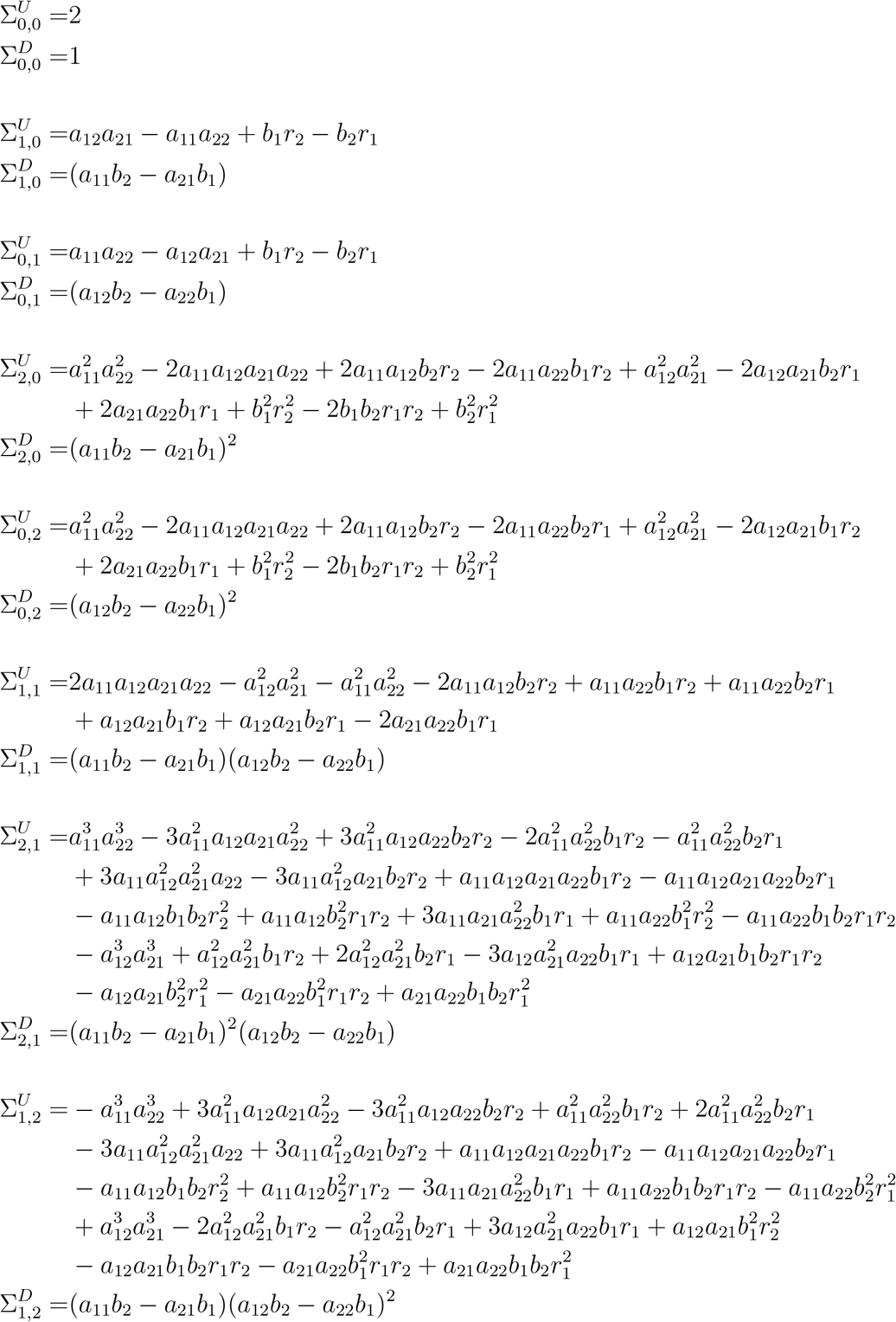

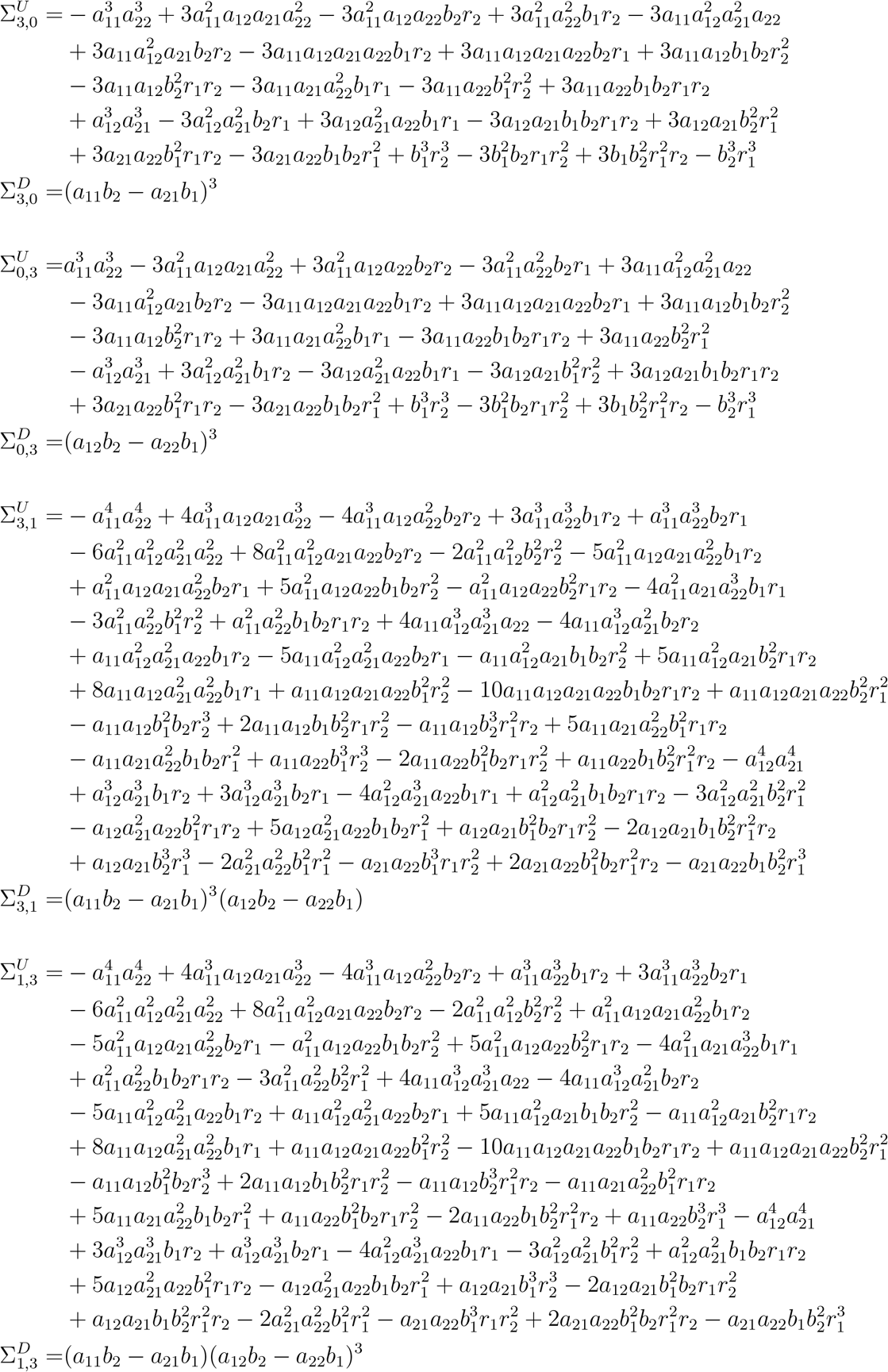

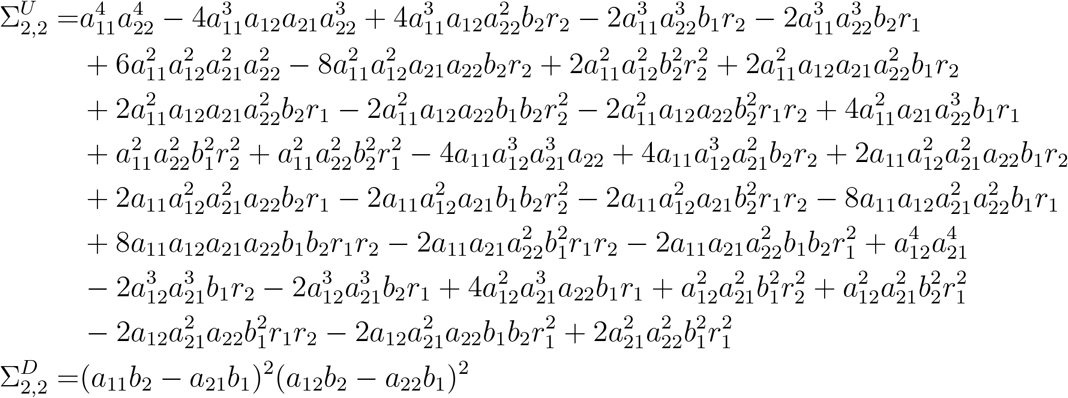

Denote the roots of *f*_1_(*N*_1_, *N*_2_) and *f*_2_(*N*_1_, *N*_2_) by ***η***_**1**_ = [*η*_**1**,1_, *η*_**1**,2_]^*T*^ and ***η***_**2**_ = [*η*_**2**,1_, *η*_**2**,2_]^*T*^. Choose a monomial map *m*(*N*_1_, *N*_2_) = [1, *c*_1_*N*_1_ + *c*_2_*N*_2_]^*T*^ for some constants *c*_1_ and *c*_2_. Then, let *Q*(*N*_1_, *N*_2_) = *N*_1_*N*_2_ and compute *S*(*s*_1_, *s*_2_) = *W* Δ*W* ^*t*^ where *W*_*ij*_ = *m*_*i*_(*η*_**1**,*j*_, *η*_**2**,*j*_) and Δ_*ii*_ = *Q*(*η*_**1**,*i*_ − *s*_1_, *η*_**2**,*i*_ − *s*_2_) is a diagonal matrix as follows.

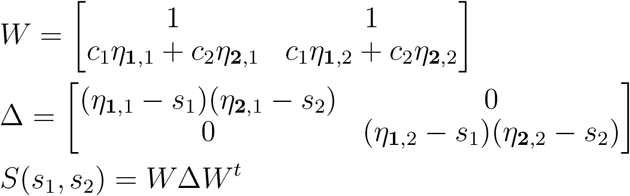

Since the symmetric sums of the roots 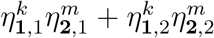 equal ∑_*k,m*_ for *k, m* = 0, 1, 2, *…*, then the components of the symmetric 2×2 matrix *S* are shown below:

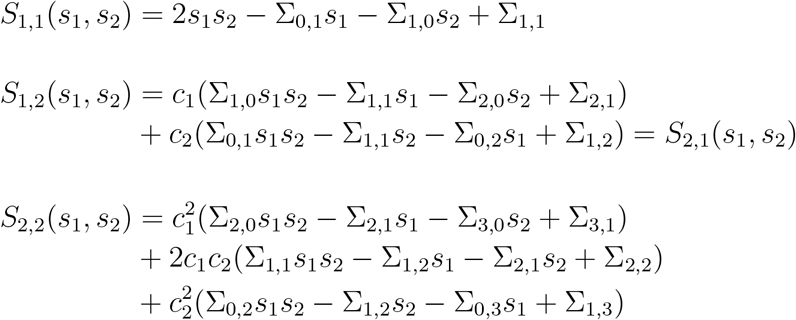

The characteristic equation of the matrix *S* is simply *λ*^2^ − Tr(*S*(*s*_1_, *s*_2_))*λ* + det(*S*(*s*_1_, *s*_2_)) whose coefficients are given by ***v*** = [1, −Tr(*S*(*s*_1_, *s*_2_)), det(*S*(*s*_1_, *s*_2_))]. Hence, the quantities of interest are −Tr(*S*(*s*_1_, *s*_2_)) and det(*S*(*s*_1_, *s*_2_)) which are shown next.

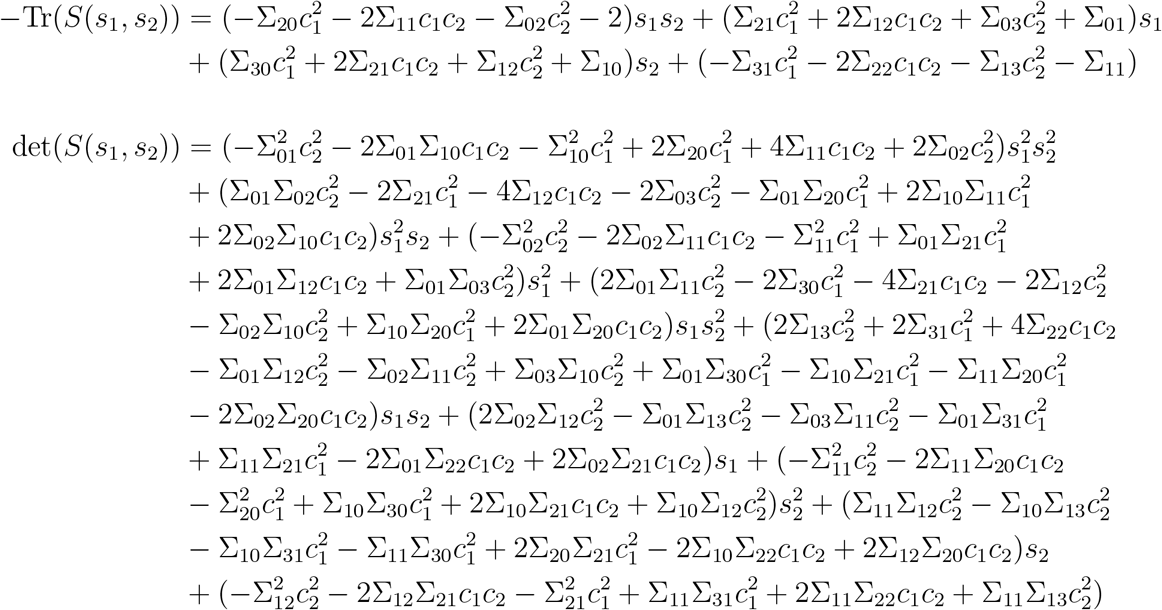

Let *V* (*s*_1_, *s*_2_) be the number of consecutive sign changes in ***v***. Since we are interested in the feasibility domain, note that the points {(0, 0), (0, ∞), (∞, 0), (∞, ∞)} are the vertices of the “box” that bound the feasibility domain. Hence, the expression of *F* (**Ψ**) is simply *F* (**Ψ**) = [*V* (0, 0) − *V* (0, ∞) − *V* (∞, 0) + *V* (∞, ∞)]/2. Here, ∞ is a limit; therefore, 0 is evaluated first before the limit is taken at infinity. For *V* (∞, ∞) where the limit of two quantities approach infinity, the limit is unique. Therefore, it does not matter along which direction the limit is taken. Now, we need to evaluate −Tr(*S*) and det(*S*) at those four vertices which are the basis to construct the inequalities that describe the feasible domain.

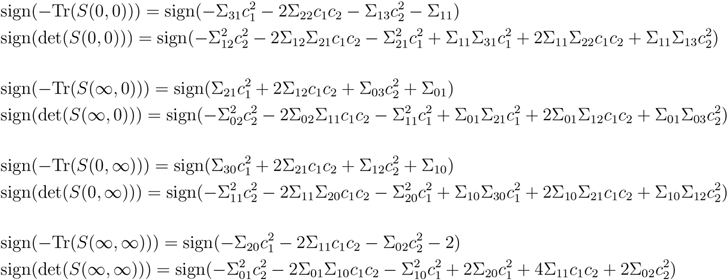

Note that the formula of *F* (**Ψ**) is completely independent of *c*_1_ and *c*_2_ and the property can be checked with our provided code. Let us set *c*_1_ = 1 and *c*_2_ = 0 for convenience. Now, the feasible domain is the set of all inequalities so that *F* (**Ψ**) ≥ 1 and found 13 non-empty ones. This was done via computing *F* (**Ψ**) for a range of parameters (**Ψ**, with each component is varied independently and uniformly between −1 and 1. There was no more increase in the number of non-empty sets (the 13 ones) when the range of each parameter is varied independently and uniformly between −100 to 100. The 13 sets are shown in the columns below and satisfying any of those guarantees feasibility. The signs − and + mean that the quantity on the left-hand most in the table is less than 0 and greater than 0 respectively. In this table, we care about conditions that satisfy *F* (**Ψ**) and we will not separate them based on whether *F* takes a value of 1 or 2.

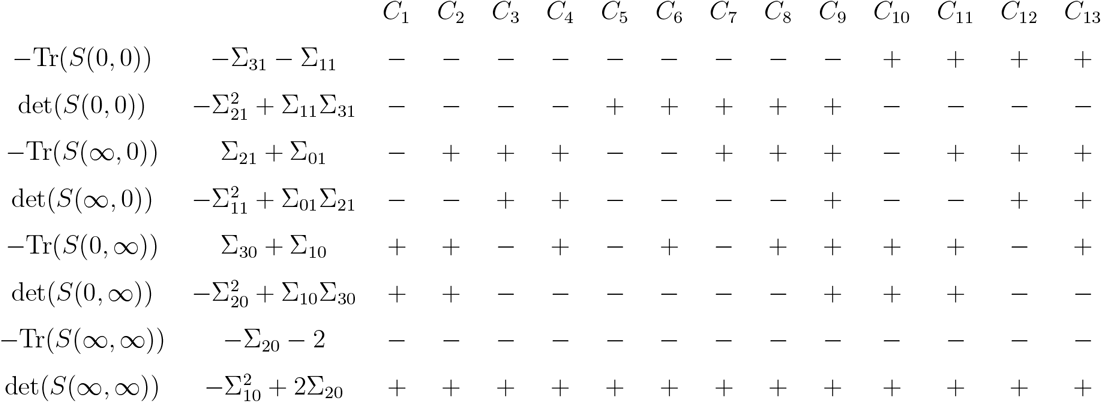

From the table, −Tr(*S*(∞, ∞)) is negative and det(*S*(∞, ∞)) is positive for all 13 conditions. This is because the relations −Σ_20_ −2 < 0 always holds (minus sum of squares minus a positive number must be negative). Also, the relation 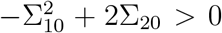 always holds, which follows from the AM-GM inequality. Hence, the last two conditions are redundant and can be eliminated as they are automatically satisfied. In addition, the 13 sets can be compressed nicely into 4 as follows. Note that columns 1 and 2 (i.e., *C*_1_ and *C*_2_) differ in sign in the third row (−Tr(*S*(∞, 0))). Hence, the two conditions can be combined into one without caring about the sign of −Tr(*S*(∞, 0)). The same applies to columns 3 and 4 (i.e., *C*_3_ and *C*_4_), 5 and 6 (i.e., *C*_5_ and *C*_6_), 7 and 8 (i.e., *C*_7_ and *C*_8_), 10 and 11 (i.e., *C*_10_ and *C*_11_) as well as 12 and 13 (i.e., *C*_12_ and *C*_13_) since these pairs of columns differ by a single sign only. The reduced table from combining columns (conditions) is shown below where X denotes to no condition:

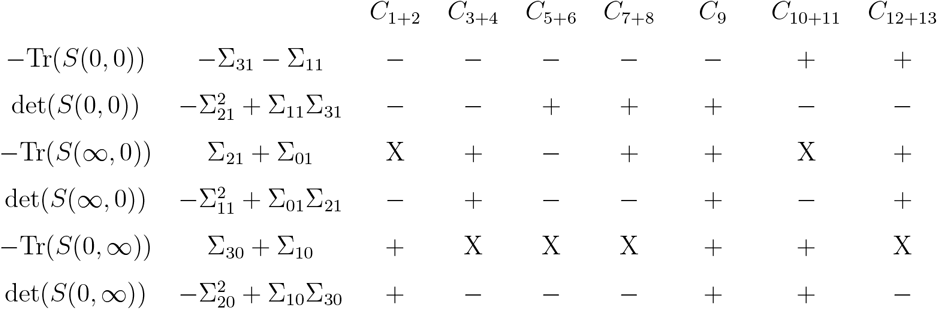

Furthermore, we can combine *C*_1+2_ with *C*_10+11_, *C*_3+4_ with *C*_12+13_ and *C*_5+6_ with *C*_7+8_ to produce the following table:

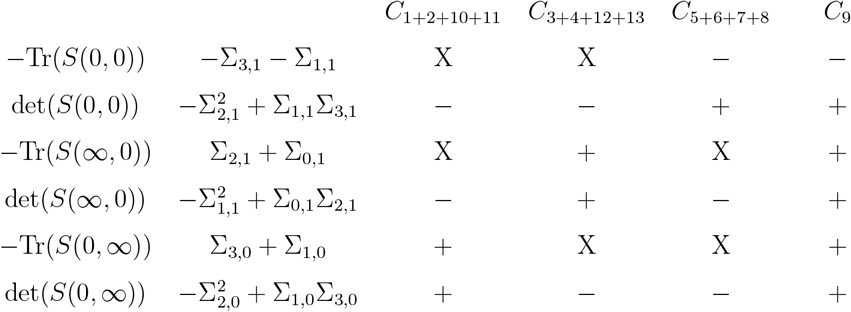

So far, we have combined columns but have not investigated whether there are redundant signs in each column. Upon computing the conditional probability that a sign occurs given that all other signs occur in the same column and keep deleting signs until there is no conditional probability of 1, we find the following minimized table that is shown below:

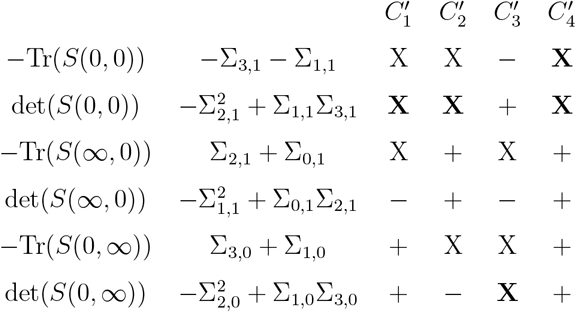

Let *Q*_*i*_ be the basis quantity in row *i*. The minimized table above is not unique. For column *C*_1_′, the user may eliminate either *Q*_2_ < 0 or *Q*_4_ < 0 in that column but not both. This is because *P* (*Q*_2_ < 0|*Q*_4_ < 0, *Q*_5_ > 0, *Q*_6_ > 0) = 1 and *P* (*Q*_4_ < 0|*Q*_2_ < 0, *Q*_5_ > 0, *Q*_6_ > 0) = 1 but upon deleting either *Q*_2_ < 0 or *Q*_4_ < 0 from these conditional probabilities, we find *P* (*Q*_2_ < 0|*Q*_5_ > 0, *Q*_6_ > 0) ≠ 1 and *P* (*Q*_4_ < 0|*Q*_5_ > 0, *Q*_6_ > 0) ≠ 1 meaning that the inequalities *Q*_2_ < 0 and *Q*_4_ < 0 imply one another given that *Q*_5_ > 0 and *Q*_6_ > 0.

## S4 Ex 2: 2-species with Type III Functional Responses

Consider the simplest 2-species LV model with type III functional responses model that is impossible to solve for the location of the equilibrium points analytically.

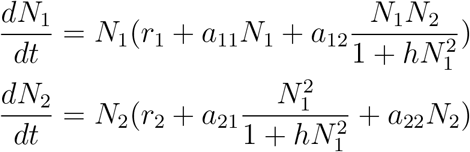

Let **Ψ** = (*r*_1_, *r*_2_, *a*_11_, *a*_12_, *a*_21_, *a*_22_, *h*) be the vector of model parameters. The common numerators of the RHS of the system above, after deleting *N*_1_ and *N*_2_ outside the brackets, are given by 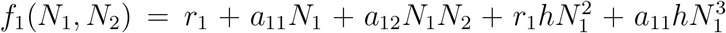 and 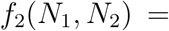 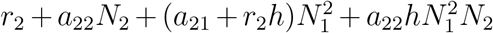 for lines 1 and 2 respectively. Upon eliminating *N*_1_ from both *f*_1_(*N*_1_, *N*_2_) and *f*_2_(*N*_1_, *N*_2_) we obtain 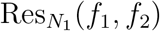 which is a polynomial of degree 5 in *N*_2_ which cannot be solved analytically in closed-form. Similarly, upon eliminating *N*_2_ from both *f*_1_(*N*_1_, *N*_2_) and *f*_2_(*N*_1_, *N*_2_) we obtain 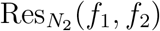 which is a polynomial of degree 5 in *N*_1_. The two resultants, each written in two forms (i.e., polynomial combination of *f*_1_ and *f*_2_ or in terms of *N*’s) are shown below:

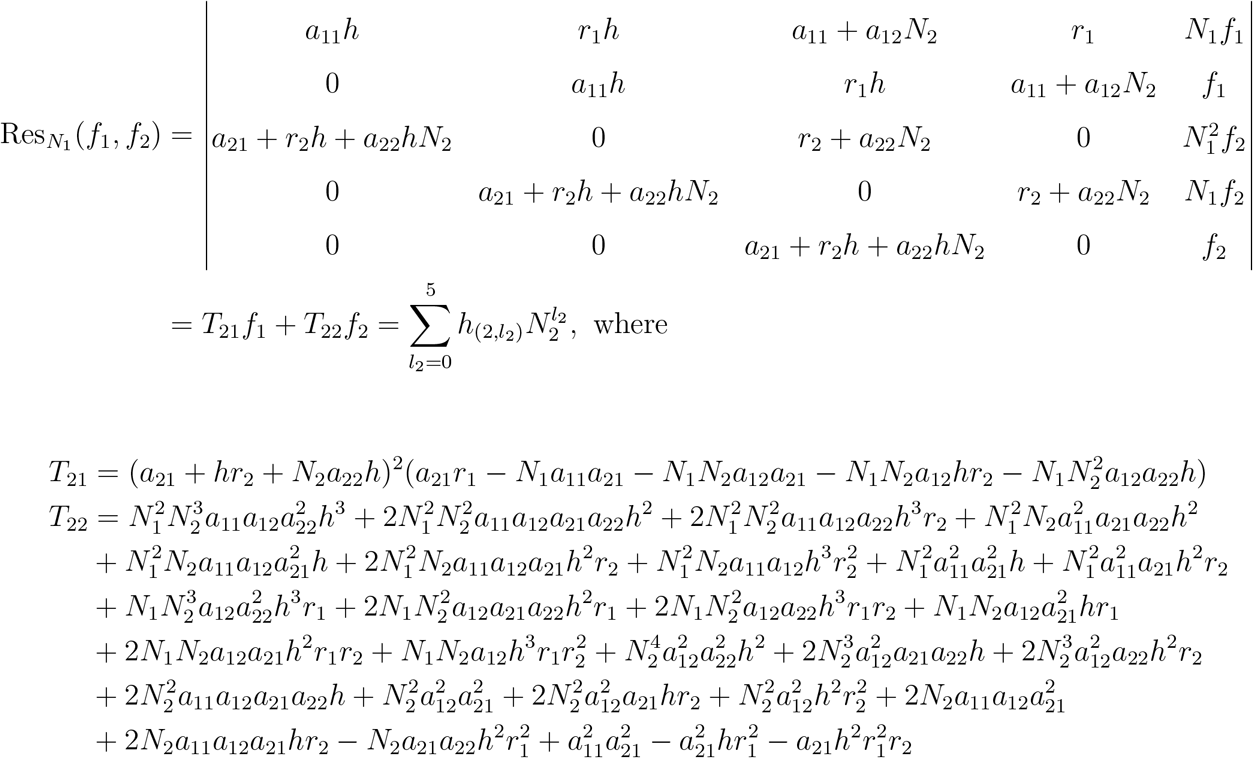

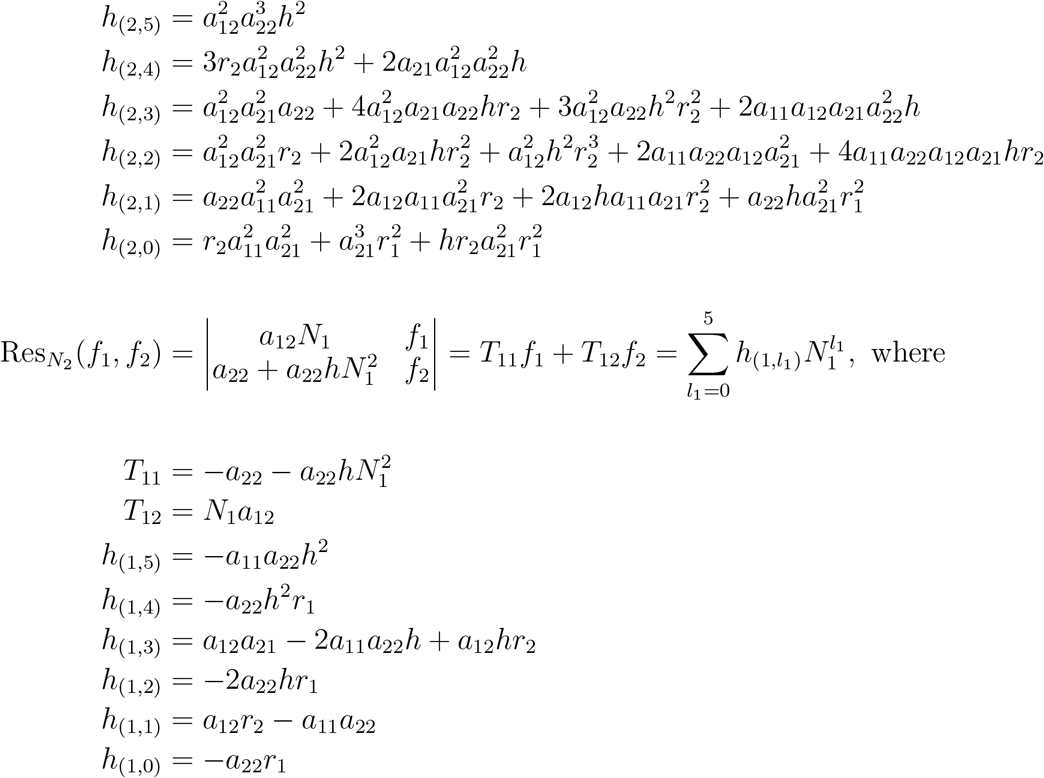

Observe that 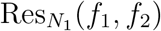 contains no *N*_1_ and is a polynomial of degree 5 in *N*_2_ only. Similarly, 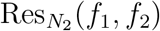 contains no *N*_2_ and is a polynomial of degree 5 in *N*_1_ only. This is an indication that the number of roots of *f*_1_(*N*_1_, *N*_2_) and *f*_2_(*N*_1_, *N*_2_) is 5. Note that the roots of the univariate polynomials 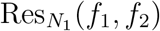 and 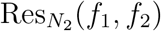, upon appropriate pairing of roots of the first polynomial with the second, are the roots of the system *f*_1_(*N*_1_, *N*_2_) = 0 and *f*_2_(*N*_1_, *N*_2_) = 0. From Abel’s impossibility theorem, since it is impossible to solve for the roots of a quintic or higher degree polynomials in terms of radicals, then the roots of either 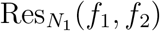 or 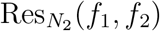 are unattainable analytically which implies that the system *f*_1_(*N*_1_, *N*_2_) = 0 and *f*_2_(*N*_1_, *N*_2_) = 0 cannot be solved. After finding the resultants in both forms, we evaluate the determinant of the eliminating matrix, which is *T* (*f*_1_, *f*_2_) = *T*_11_*T*_22_ − *T*_12_*T*_21_ and the determinant of the Jacobian of *f*_1_ and *f*_2_ as following

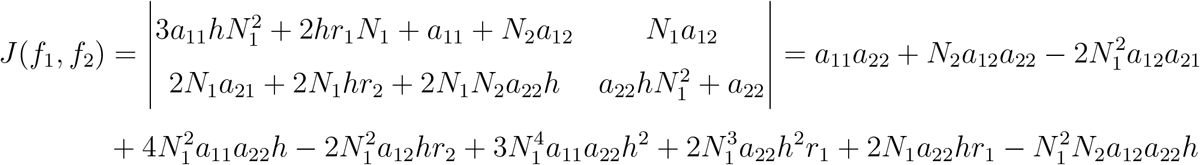

Now, we need to expand the generating function *G*(*f*_1_(*N*_1_, *N*_2_), *f*_2_(*N*_1_, *N*_2_)) around *N*_1_ = ∞ and *N*_2_ = ∞ (no need to perform a two-variable series expansion). Since 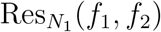 and 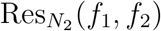 are univariate polynomials, we expand their reciprocal individually to get the series 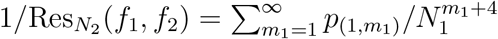 and 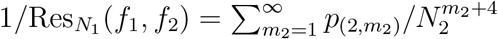 The coefficients can be obtained via MATLAB’s ‘taylor’ function where *N*_1_ and *N*_2_ are substituted by 1/*x* and 1/*y* respectively. Alternatively, these *p*’s can be obtained analytically as follows. We have already written 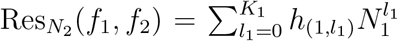 and 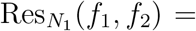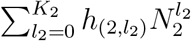 where we have *K*_1_ = *K*_2_ = 5. After that, construct the following 2 matrices *A*_1_ and *A*_2_

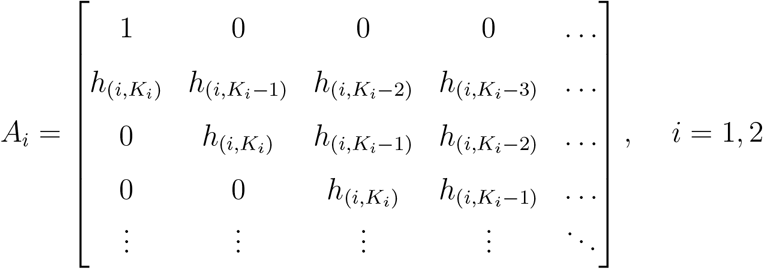

Next, let 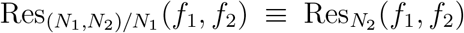 and 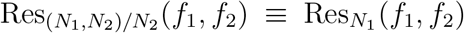. The reciprocal of each resultant is given by

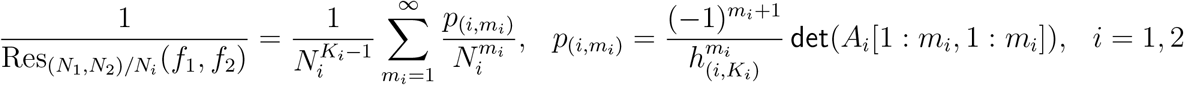

Here, *A*_*i*_[1 : *m*_*i*_, 1 : *m*_*i*_] is the sub-matrix of *A*_*i*_ that contains its first *m*_*i*_ rows and columns. After obtaining both series expansion of the resultant reciprocal, multiply the result by *T* (*f*_1_, *f*_2_)*J* (*f*_1_, *f*_2_) to obtain

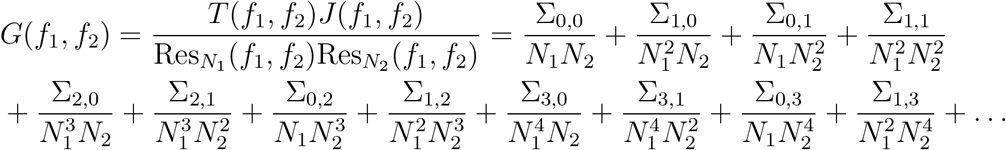

The expression for each of the Σ’s (symmetric power sums of the roots) are shown below where 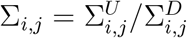 is written as a fraction of two polynomials.

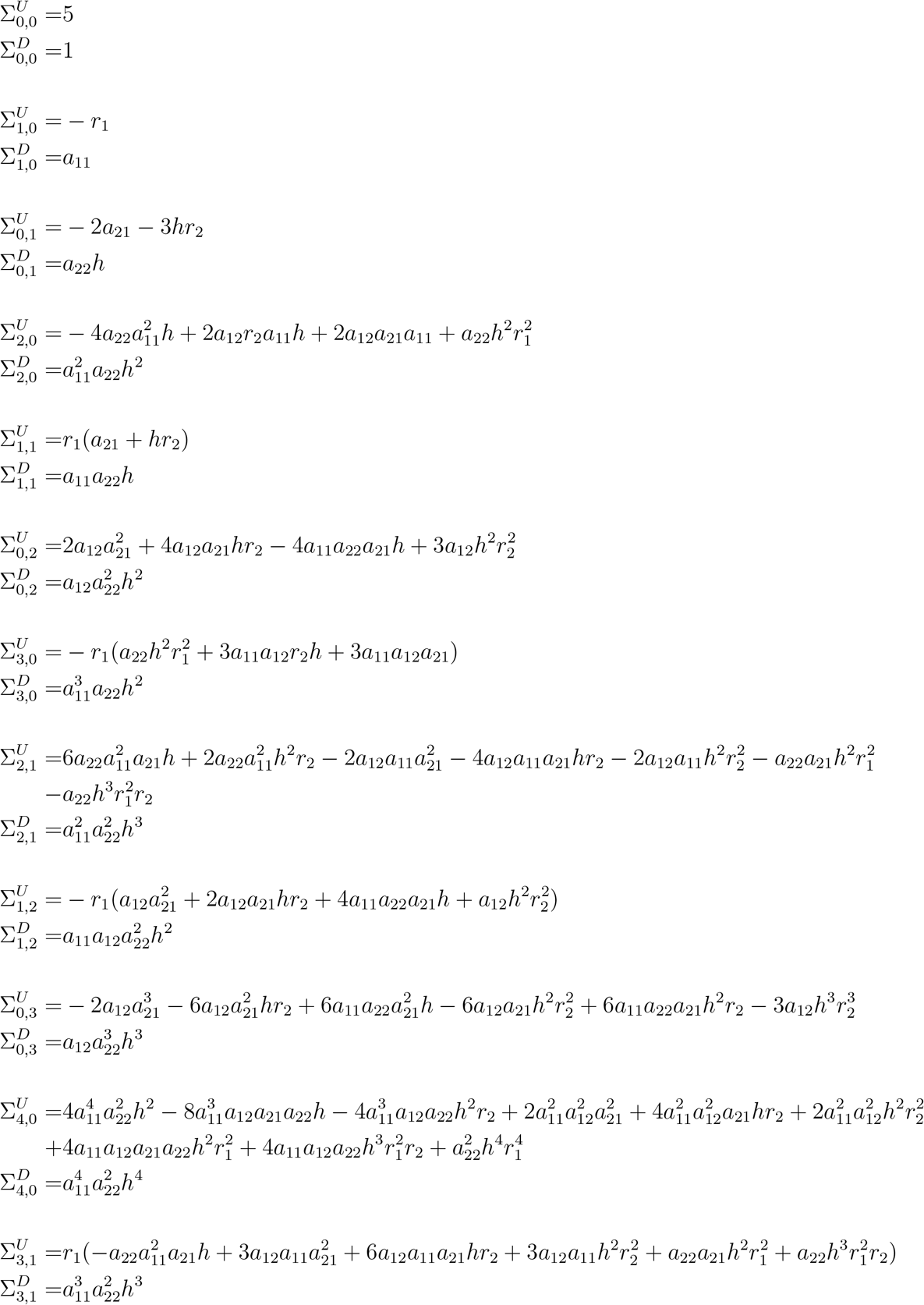

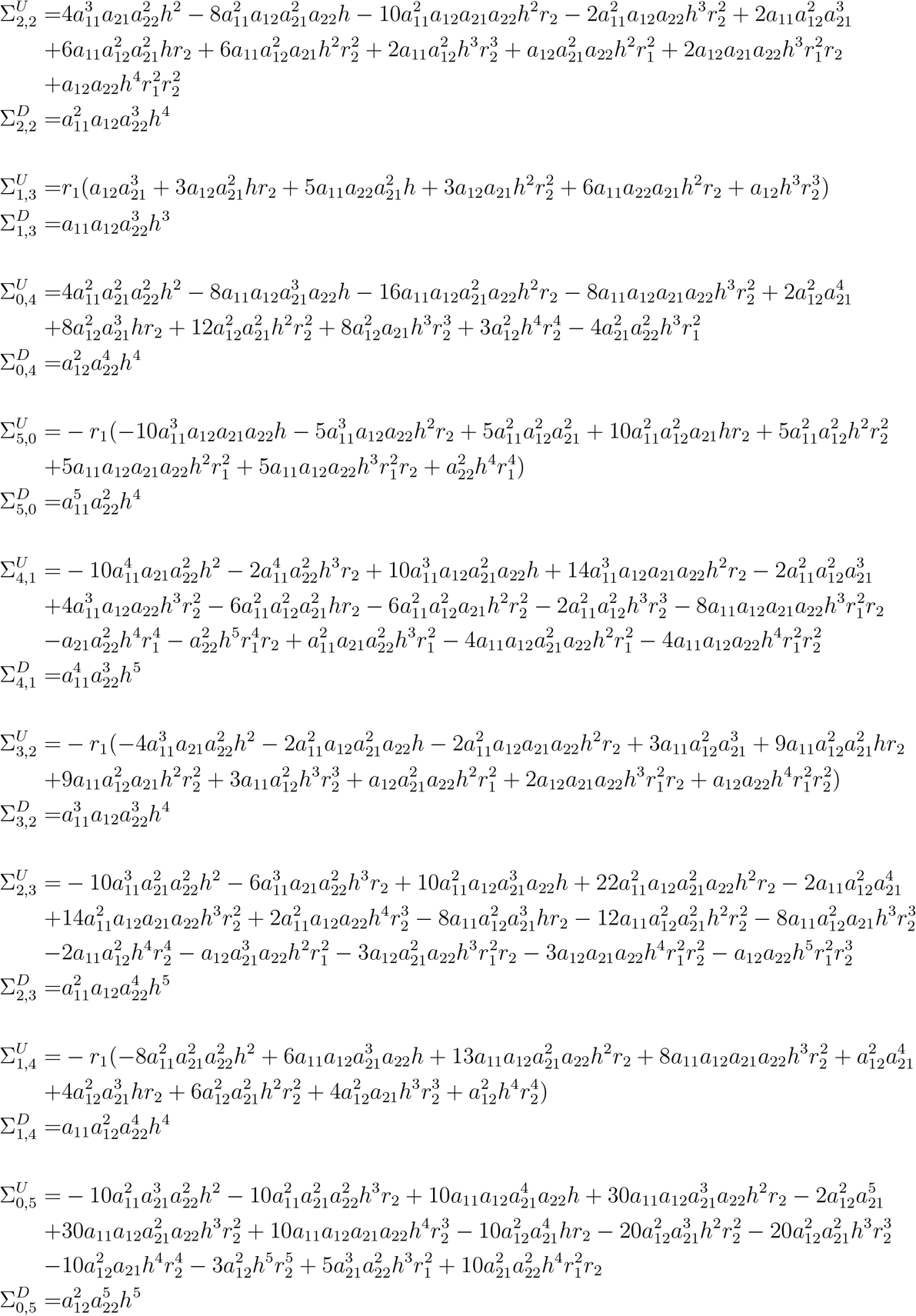

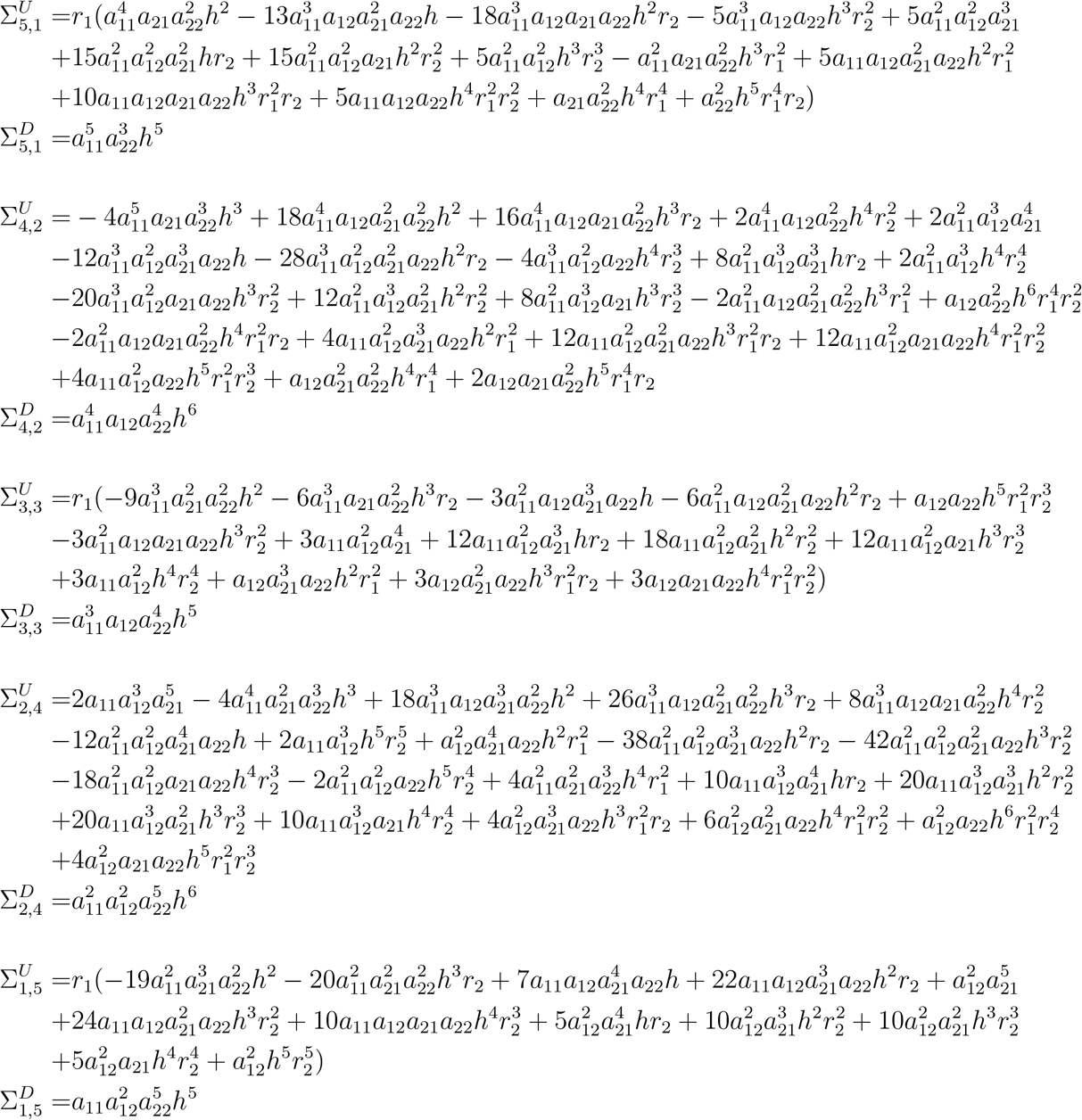

Note that if any of the parameters *a*_11_, *a*_12_, *a*_22_, *h* is zero, the Σ’s will blow up. If one needs to consider cases where any of the latter parameters is zero, that zero should be first substituted in *f*_1_(*N*_1_, *N*_2_) and *f*_2_(*N*_1_, *N*_2_) before carrying on with what we have shown already. Next, denote the roots of *f*_1_(*N*_1_, *N*_2_) and *f*_2_(*N*_1_, *N*_2_) by ***η***_**1**_ = [*η*_**1**,1_, *η*_**1**,2_, …, *η*_**1**,5_]^*T*^ and ***η***_**2**_ = [*η*_**2**,1_, *η*_**2**,2_, …, *η*_**2**,5_]^*T*^. Choose a monomial map 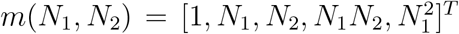 then, let *Q*(*N*_1_, *N*_2_) = *N*_1_*N*_2_ and compute *S*(*s*_1_, *s*_2_) = *W* Δ*W*^*t*^ where *W*_*ij*_ = *m*_*i*_(*η*_**1**,*j*_, *η*_**2**,*j*_) and Δ_*ii*_ = *Q*(*η*_**1**,*i*_ − *s*_1_, *η*_**2**,*i*_ − *s*_2_) is a diagonal matrix as follows.

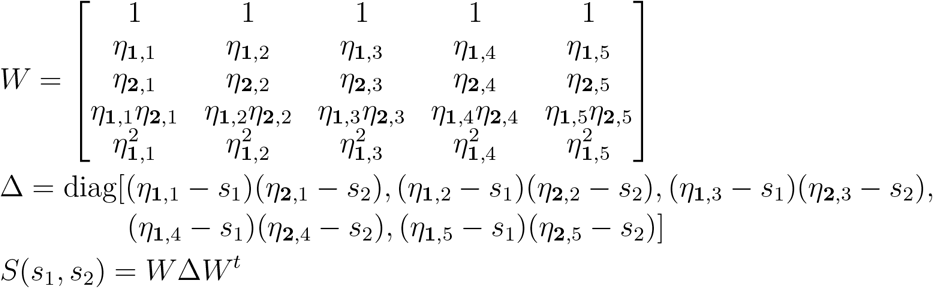

Note that 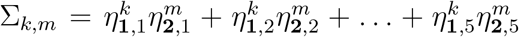 for *k, m* = 0, 1, 2, *…*. Therefore, the components of the symmetric 5×5 matrix *S* are shown below:

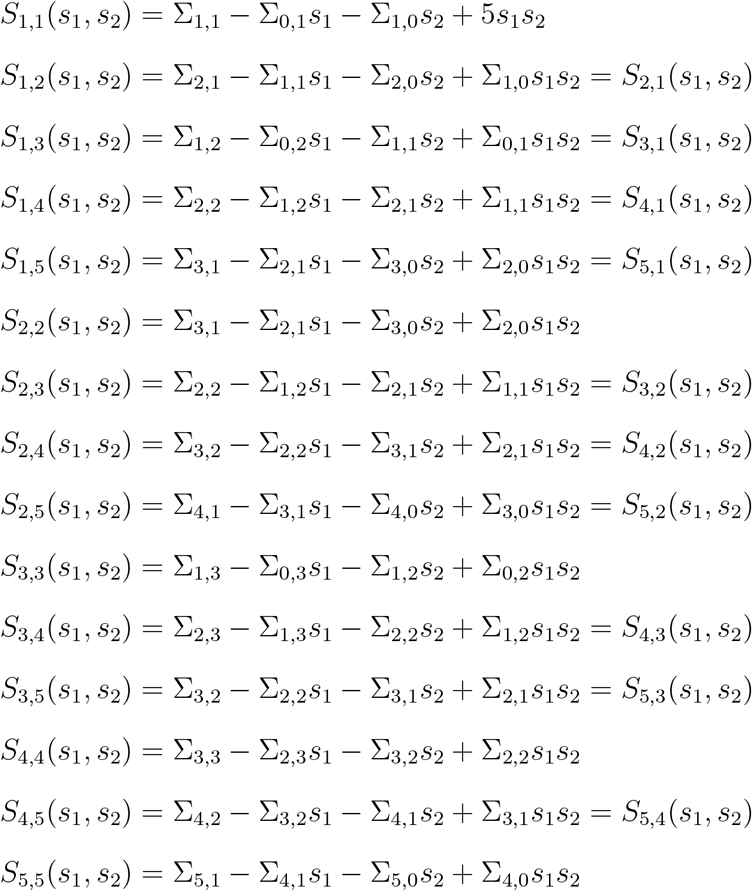

The characteristic equation of the matrix *S* is simply det(*S*(*s*_1_, *s*_2_)) = *λ*^5^ + *v*_4_(*s*_1_, *s*_2_)*λ*^4^ + *v*_3_(*s*_1_, *s*_2_)*λ*^3^ + *v*_2_(*s*_1_, *s*_2_)*λ*^2^ + *v*_1_(*s*_1_, *s*_2_)*λ + v*_0_(*s*_1_, *s*_2_). The coefficients of the characteristic equation evaluated at (*s*_1_, *s*_2_) = {(0, 0), (∞, 0), (0, ∞), (∞, ∞)} are shown in the following pages. Note that *v*_*i*_(0, 0),*v*_*i*_(∞, 0),*v*_*i*_(0, ∞) and *v*_*i*_(∞, ∞) are the coefficients of (*s*_1_*s*_2_)^0^, 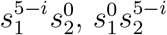 and (*s*_1_*s*_2_)^5−*i*^ of *v*_*i*_(*s*_1_, *s*_2_) respectively for *i* = 0, 1, *…*, 5.

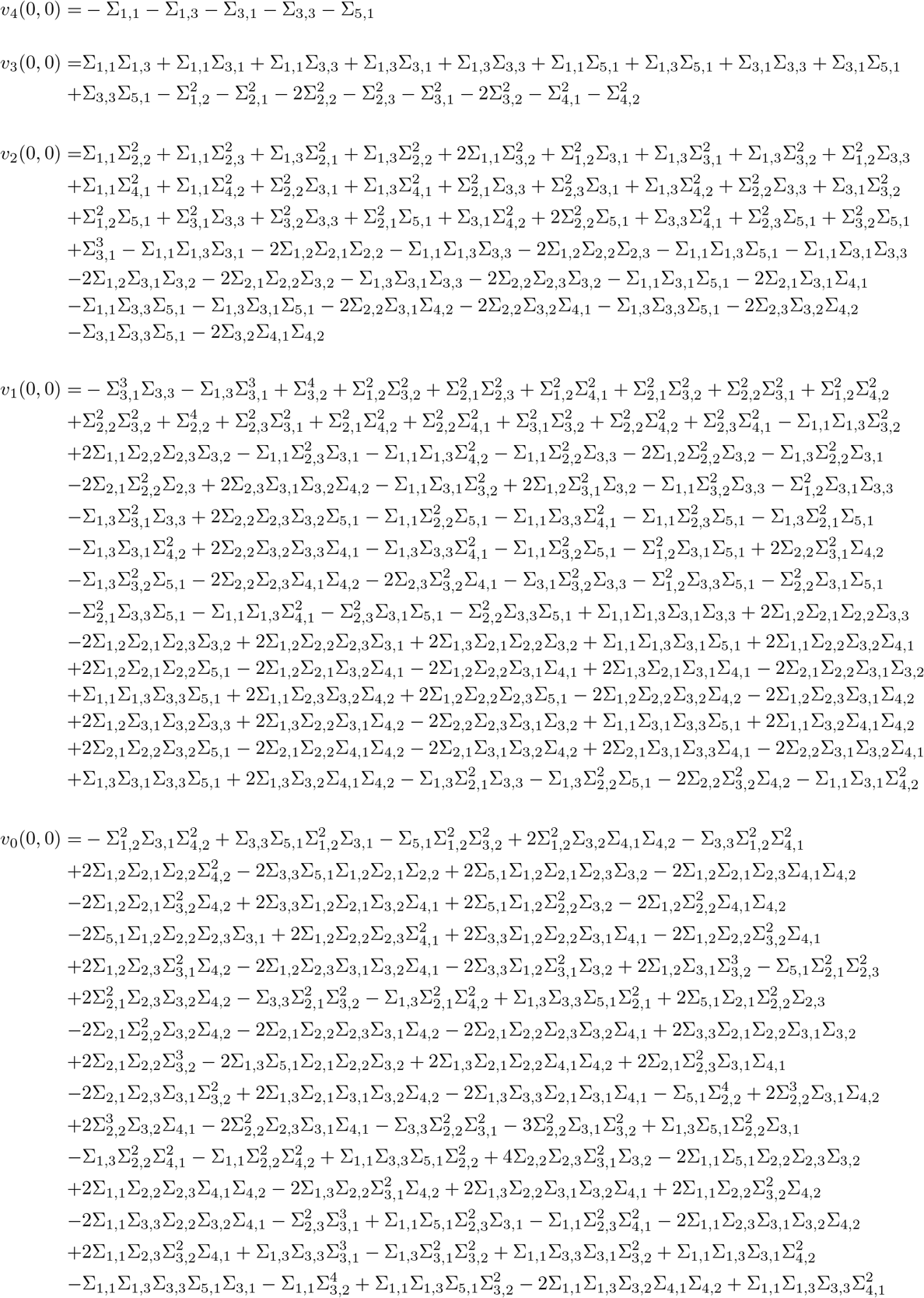

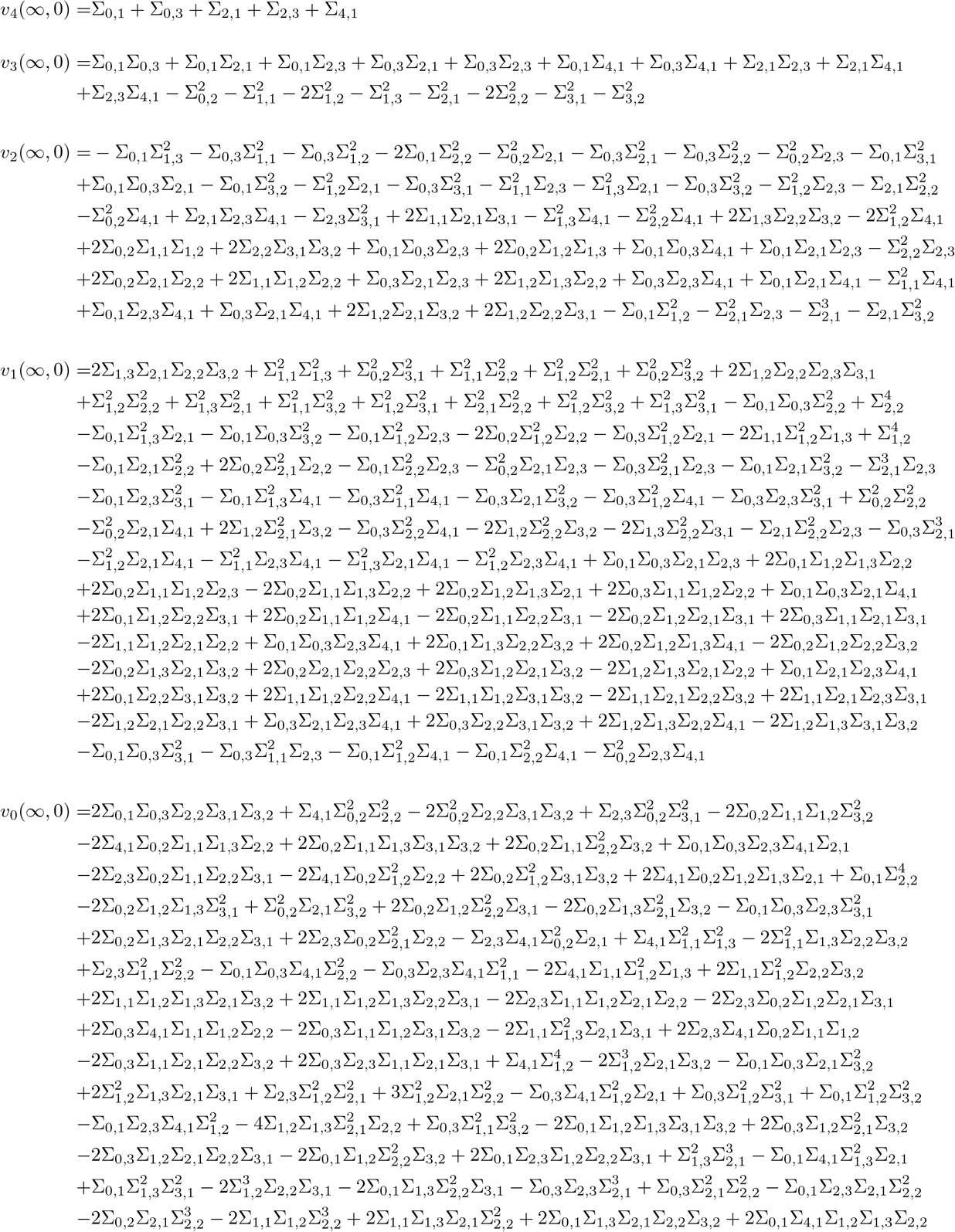

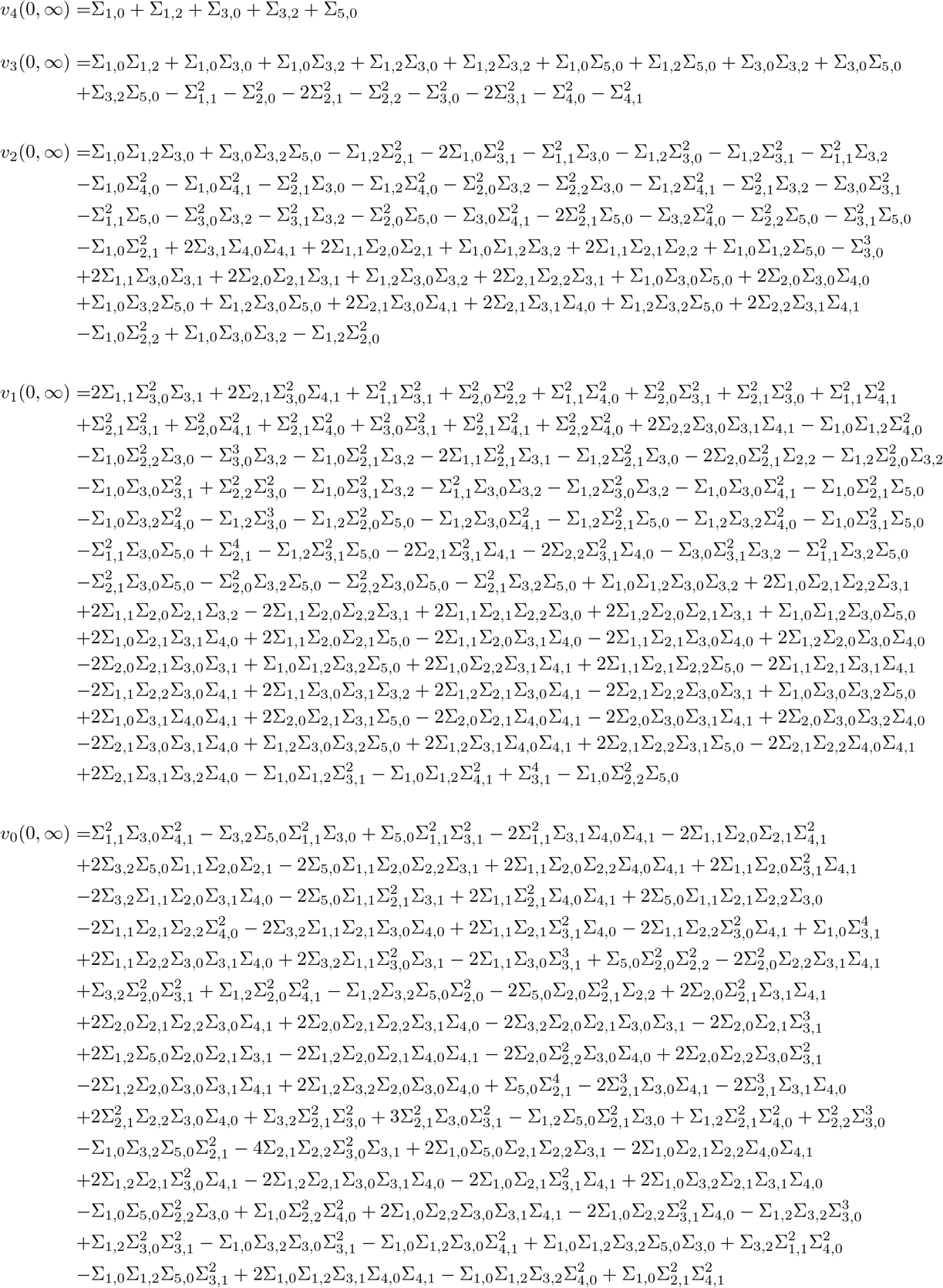

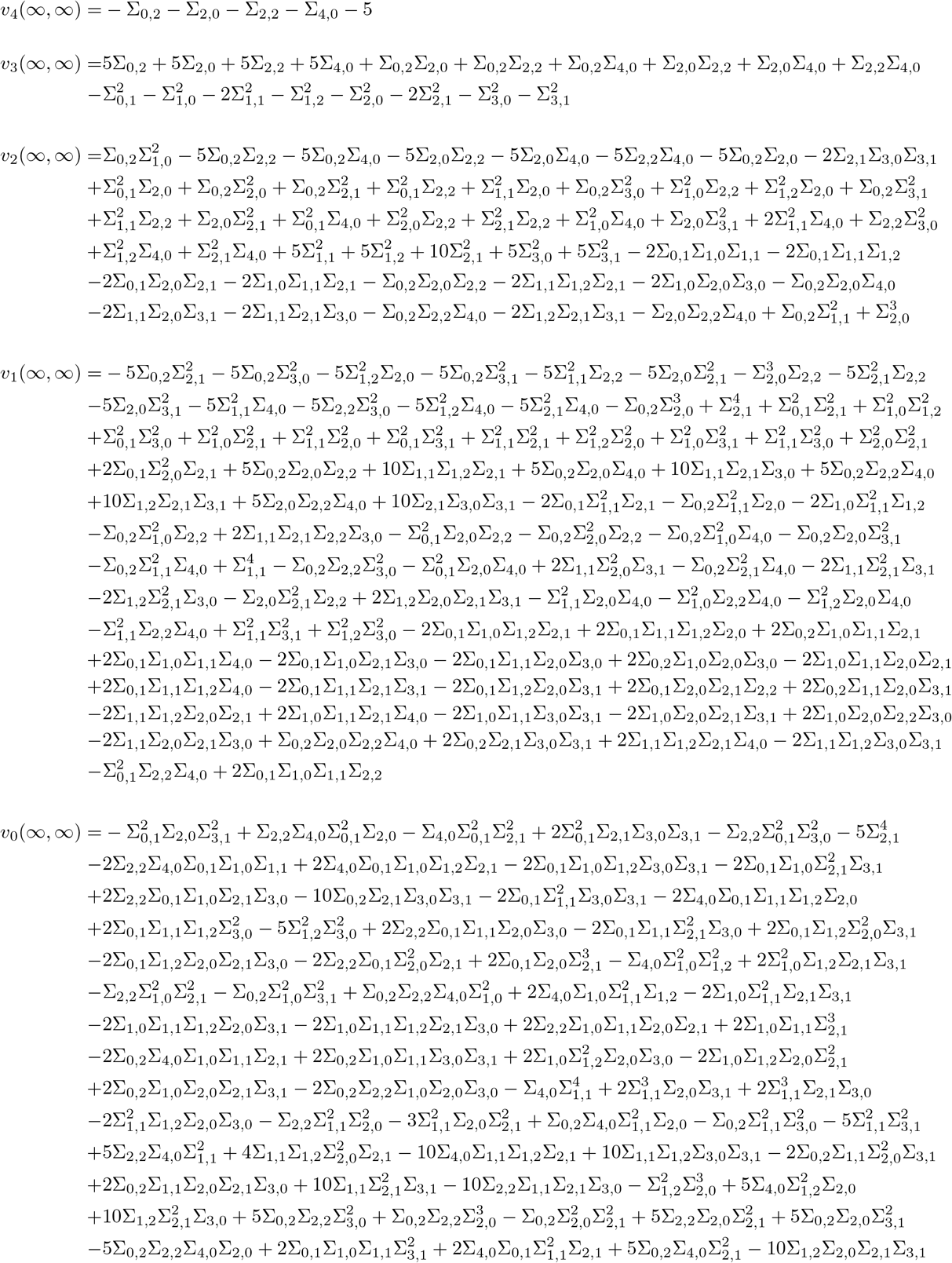

Let *V* (*a, b*) be the number of consecutive sign changes in [1, *v*_4_(*a, b*), *v*_3_(*a, b*), *v*_2_(*a, b*), *v*_1_(*a, b*), *v*_0_(*a, b*)] where *a* and *b* are either 0 or ∞. The formula of *V* (*a, b*) is shown below

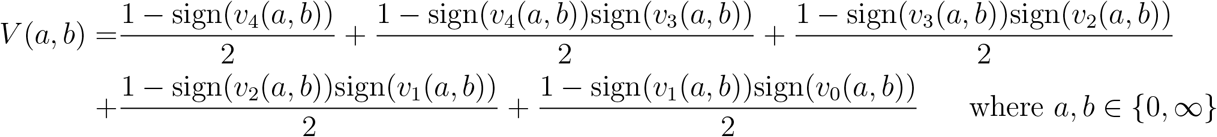

From the *V*’s, we can find the formula of the number of feasible roots of *f*_1_(*N*_1_, *N*_2_) and *f*_2_(*N*_1_, *N*_2_) which is given by *F* (**Ψ**) = (*V* (0, 0) − *V* (∞, 0) − *V* (0, ∞) + *V* (∞, ∞))/2. The feasibility table for this example is huge and instead of finding the link between all parameters while maintaining feasibility, we will perform a demo on how to find the link between the parameters *h* and *a*_21_ while they are in a restricted domain. Let us consider the parameter vector **Ψ** = (*r*_1_, *r*_2_, *a*_11_, *a*_12_, *a*_21_, *a*_22_, *h*) = (0.5, −1.5, 1, −1.5, *a*_21_, 1, *h*) where the parameters *a*_21_ ∈ [−6, −1] and *h* ∈ [0.5, 4] are restricted, we find that feasibility (i.e., *F* (**Ψ**) ≥ 1) can only be satisfied under the single condition that is shown below:

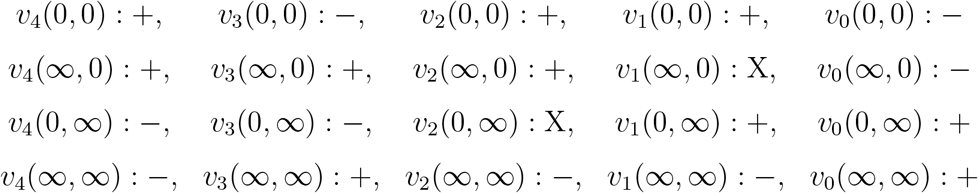

Without the need to compute conditional probabilities, upon plotting the signs of the *v*’s in the single condition above, we find that feasibility is maintained if and only if *v*_0_(0, 0) < 0. To check that our finding is correct, we plot both *F* (**Ψ**) and sign(*v*_0_(0, 0)) in the grid *a*_21_ ∈[−6, −1] and *h* ∈ [0.5, 4]. From the two plots, we can see that *F* (**Ψ**) > 0 if and only if *v*_0_(0, 0) < 0 and that *a*_21_ and *h* are related. Nevertheless, in other domains, *a*_21_ and *h* may not be linked. For example, with the same parameter values that were chosen earlier, if we change *r*_1_ from 0.5 to −0.5, the number of feasible roots in the domain *a*_21_ ∈ [−6, −1] and *h* ∈ [0.5, 4] will be exactly 1 no matter what values *a*_21_ and *h* take. Of course, the demo that we have illustrated here can be applied to any parameter ranges and combination. It is true that the expression of *v*_0_(0, 0) is huge and messy, however it can compacted by factoring it after plugging the fixed values into it, which is not needed for this example.

**Supplementary Figure S1:**
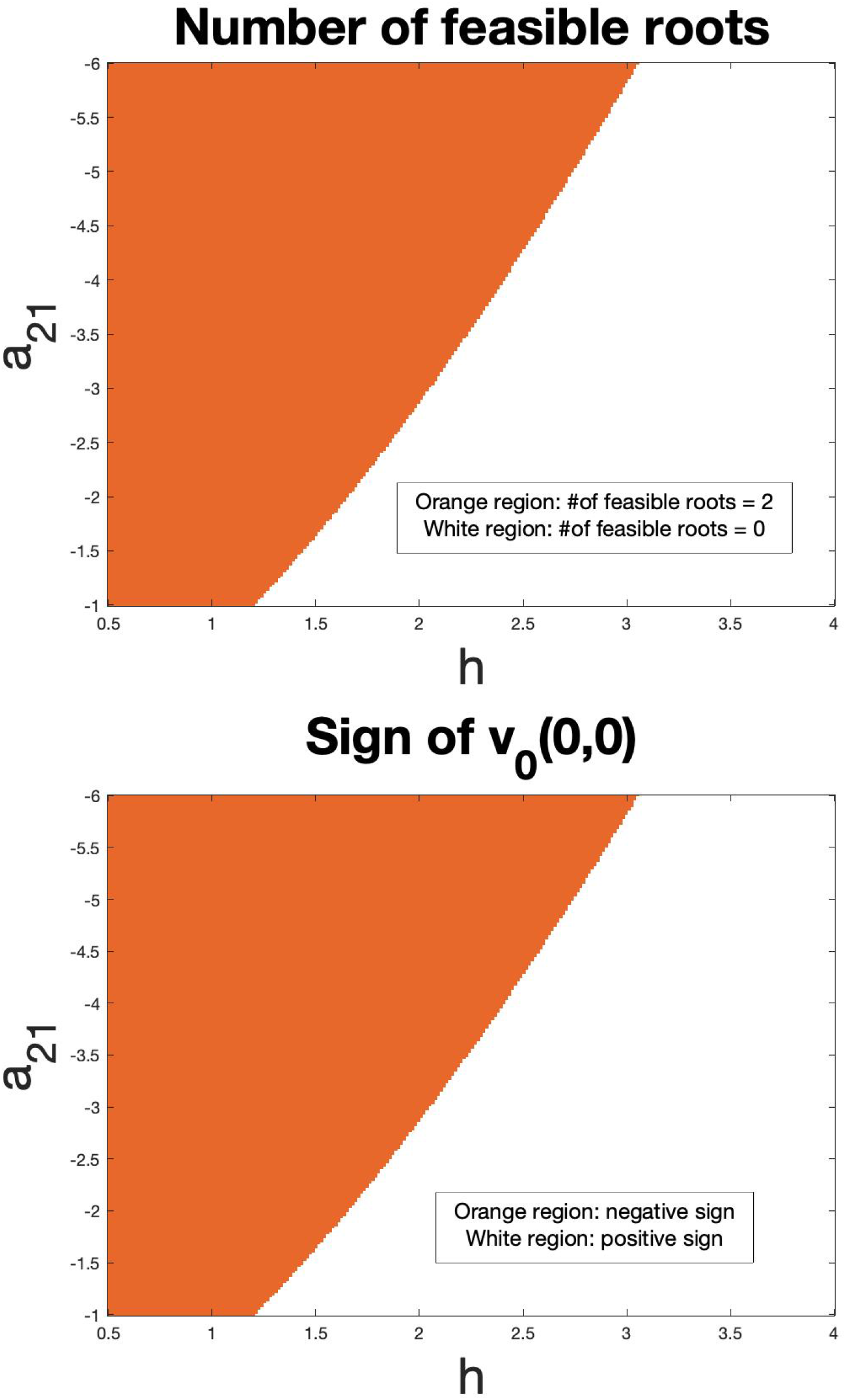
The top figure shows the number of feasible roots *F* in Lotka-Volterra model with type III functional responses where (*r*_1_, *r*_2_, *a*_11_, *a*_12_, *a*_22_) = (0.5, −1.5, 1, −1.5, 1), *a*_21_ ∈ [−6, −1] and *h* ∈ [0.5, 4]. The bottom figure shows the sign of *v*_0_(0, 0) with the same model and parameter values and ranges. Both figures confirm that *F* > 0 if and only if *v*_0_(0, 0) < 0. Simulations done via solving the isocline equations numerically and checking for the feasibility of roots match the two figures displayed here.

## S5 Ex 3: 3-Species with Simple Higher-Order Terms

Consider the dynamical system that is shown below

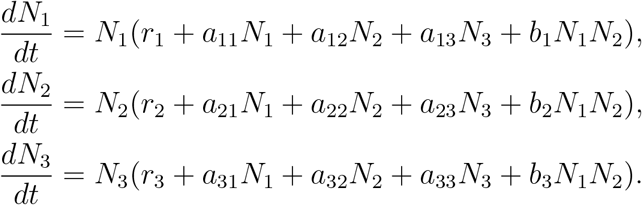

To study feasibility, the polynomials that are needed to be considered are *f*_1_(*N*_1_, *N*_2_, *N*_3_) = *r*_1_ + *a*_11_*N*_1_ + *a*_12_*N*_2_ + *a*_13_*N*_3_ + *b*_1_*N*_1_*N*_2_, *f*_2_(*N*_1_, *N*_2_, *N*_3_) = *r*_2_ + *a*_21_*N*_1_ + *a*_22_*N*_2_ + *a*_23_*N*_3_ + *b*_2_*N*_1_*N*_2_ and *f*_3_(*N*_1_, *N*_2_, *N*_3_) = *r*_3_+*a*_31_*N*_1_+*a*_32_*N*_2_+*a*_33_*N*_3_+*b*_3_*N*_1_*N*_2_. Next, assume that *N*_1_ is constant and homogenize *f*_1_, *f*_2_ and *f*_3_ with a forth variable *W* as follows:

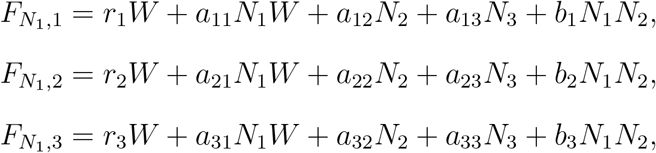

Note that the total degree of each of 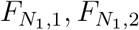 and 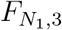 (or the total degree of *f*_1_, *f*_2_ and *f*_3_ assuming *N*_1_ is a constant) is *d*_1,1_ = 1, *d*_1,2_ = 1 and *d*_1,3_ = 1 respectively. From the *d*’s, we compute 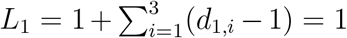. Now, we form the monomial set *H*_1_, which is a union of three disjoint monomials 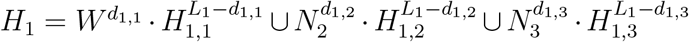 where none of these *H*’s involve *N*_1_ and each is indicated below in curly brackets:

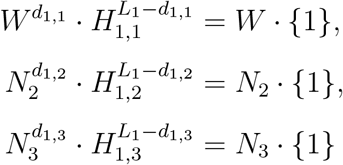

Next, form the monomial set 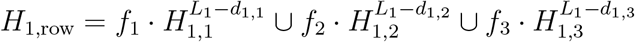 evaluated at *W* = 1 that is shown below. In addition, form the monomial set *H*_1,col_ which is simply *H*_1_ evaluated at *W* = 1 to get

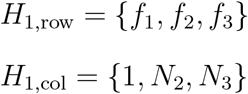

After that, form the Macaulay matrix 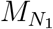 which is a square matrix whose size is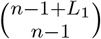 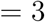. The (*i, j*) entry of the Macaulay matrix is the coefficient of *H*_1,col_(*j*) in the expression of *H*_1,row_(*i*) assuming that *N*_1_ is a constant. For example, the (1, 2) entry in the matrix is the coefficient of *N*_2_ in *f*_1_ which is *a*_12_ + *b*_1_*N*_1_. The matrix 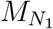 is shown below:

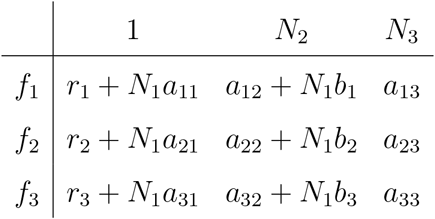

Next, form the matrix 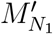 whose first column is *H*_1,row_ and its remaining columns are the remaining columns (i.e., columns 2 to 3) of the matrix 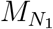 (i.e., replace the first column of 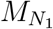 whose top header is 1 with the leftmost column which contains the *f*’s). From the formula of 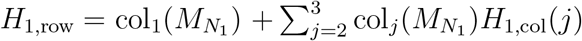, we can see that *H*_1,row_ is the first column of 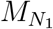 added to it a multiple of every other column of 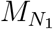, implying that 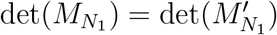. This determinant (i.e., 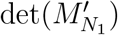) can be written as *T*_11_*f*_1_ + *T*_12_*f*_2_ + *T*_13_*f*_3_ where the formulas of *T*_11_, *T*_12_ and *T*_13_ are shown below.

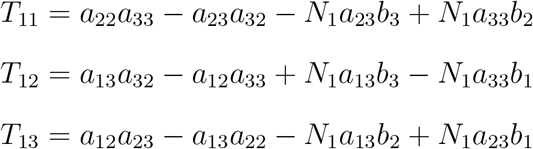

Upon substituting *f*_1_, *f*_2_ and *f*_3_ into *T*_11_*f*_1_ + *T*_12_*f*_2_ + *T*_13_*f*_3_ and simplifying the expression (or evaluating the determinant of the matrix 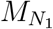 directly), we have the formula of the resultant 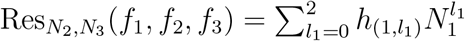 which is a polynomial of degree 2 in *N*_1_ and contains no *N*_2_’s nor *N*_3_’s. The three coefficients of the resultant *h*_(1,2)_, *h*_(1,1)_ and *h*_(1,0)_ are shown below. Notice that none of the coefficients contain any of the *N*’s.

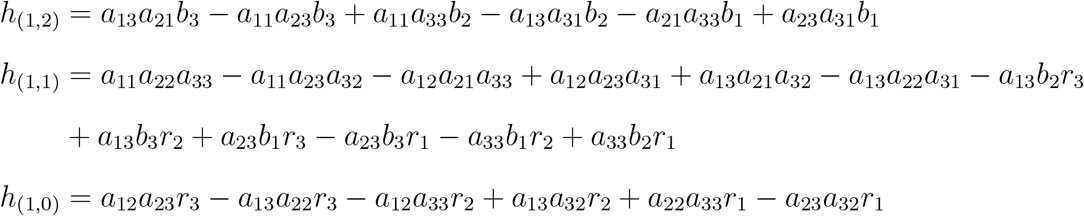

Next, assume that *N*_2_ is constant and homogenize *f*_1_, *f*_2_ and *f*_3_ with a forth variable *W* as follows:

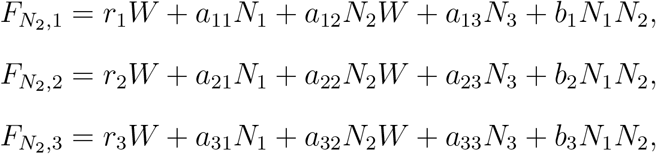

Note that the total degree of each of 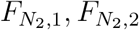 and 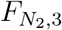 (or the total degree of *f*_1_, *f*_2_ and *f*_3_ assuming *N*_2_ is a constant) is *d*_2,1_ = 1, *d*_2,2_ = 1 and *d*_2,3_ = 1 respectively. From the *d*’s, we compute 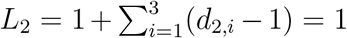. Now, we form the monomial set *H*_2_, which is a union of three disjoint monomials 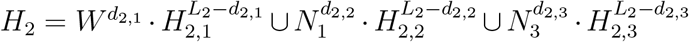 where none of these *H*’s involve *N*_2_ and each is indicated below:

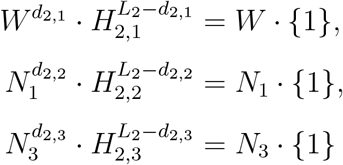

Next, form the monomial set 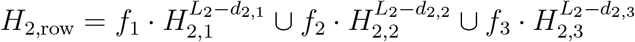 evaluated at *W* = 1 that is shown below. In addition, form the monomial set *H*_2,col_ which is simply *H*_2_ evaluated at *W* = 1 to get

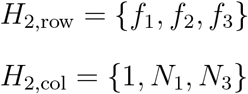

After that, form the Macaulay matrix 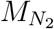 which is a square matrix whose size is 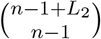 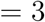. The (*i, j*) entry of the Macaulay matrix is the coefficient of *H*_2,col_(*j*) in the expression of *H*_2,row_(*i*) assuming that *N*_2_ is a constant. For example, the (1, 2) entry in the matrix is the coefficient of *N*_1_ in *f*_1_ which is *a*_11_ + *N*_2_*b*_1_. The matrix 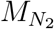 is shown below:

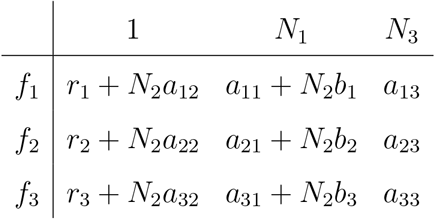

Next, form the matrix 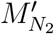 whose first column is *H*_2,row_ and its remaining columns are the remaining columns (i.e., columns 2 to 3) of the matrix 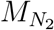 (i.e., replace the first column of 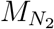 whose top header is 1 with the leftmost column which contains the *f*’s). Again, from the formula of 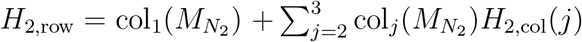, we can see that *H*_2,row_ is the first column of 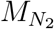 added to it a multiple of every other column of 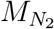, implying that 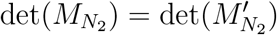. This determinant (i.e., 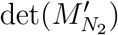) can be written as *T*_21_*f*_1_ + *T*_22_*f*_2_ + *T*_23_*f*_3_ where the formulas of *T*_21_, *T*_22_ and *T*_23_ are shown below.

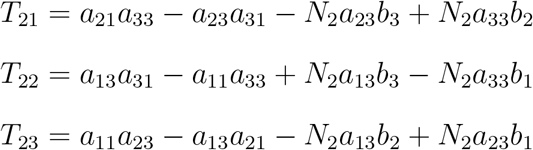

Upon substituting *f*_1_, *f*_2_ and *f*_3_ into *T*_21_*f*_1_ + *T*_22_*f*_2_ + *T*_23_*f*_3_ and simplifying the expression (or evaluating the determinant of the matrix 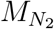 directly), we have the formula of the resultant 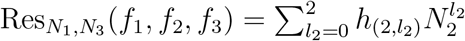 which is a polynomial of degree 2 in *N*_2_ and contains no *N*_1_’s nor *N*_3_’s. The three coefficients of the resultant *h*_(2,2)_, *h*_(2,1)_ and *h*_(2,0)_ are shown below. Notice that none of the coefficients contain any of the *N*’s.

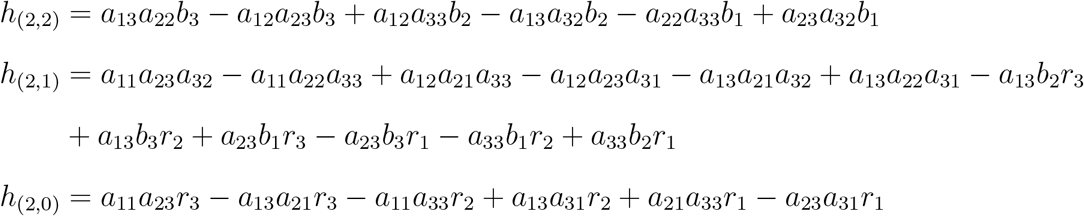

Next, assume that *N*_3_ is constant and homogenize *f*_1_, *f*_2_ and *f*_3_ with a forth variable *W* as follows:

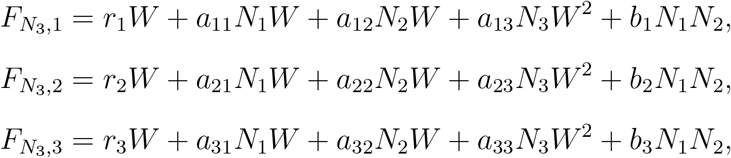

Note that the total degree of each of 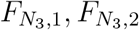 and 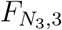 (or the total degree of *f*_1_, *f*_2_ and *f*_3_ assuming *N*_3_ is a constant) is *d*_3,1_ = 2, *d*_3,2_ = 2 and *d*_3,3_ = 2 respectively. From the *d*’s, we compute 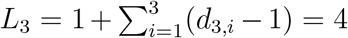. Now, we form the monomial set *H*_3_, which is a union of three disjoint monomials 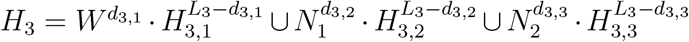 where none of these *H*’s involve *N*_3_ and each is indicated below in curly brackets:

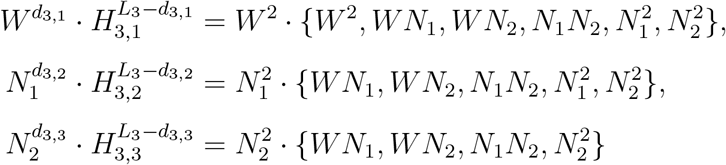

Next, form the monomial set 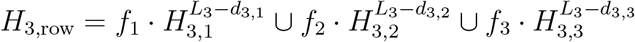 evaluated at *W* = 1 that is shown below. In addition, form the monomial set *H*_3,col_ which is simply *H*_3_ evaluated at *W* = 1 to get

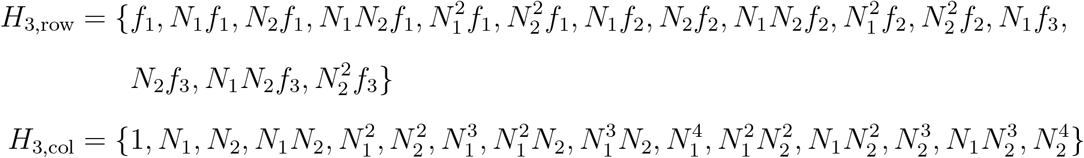

After that, form the Macaulay matrix 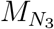 which is a square matrix whose size is 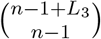 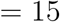. The (*i, j*) entry of the Macaulay matrix is the coefficient of *H*_3,col_(*j*) in the expression of *H*_3,row_(*i*) assuming that *N*_3_ is a constant. For example, the (1, 2) entry in the matrix is the coefficient of *N*_1_ in *f*_1_ which is *a*_11_. The matrix 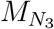 is shown below:

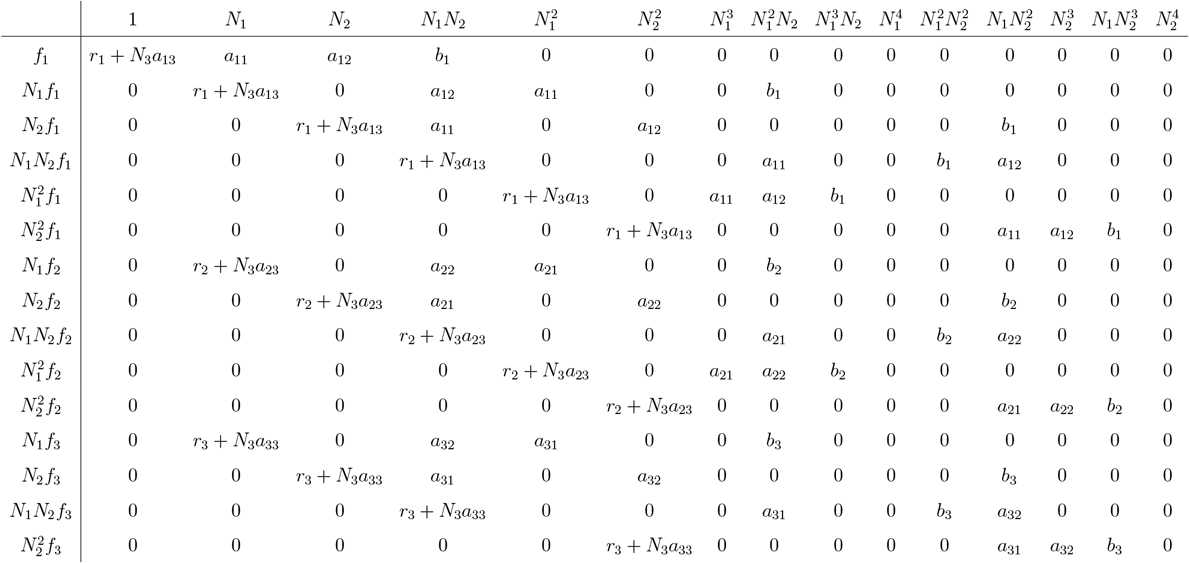

From columns 10 and 15 which are all zeros, we can see that the determinant of the matrix 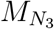 is zero. Therefore, the resultant substitution is the non-vanishing coefficient of the smallest power of *ϵ* in 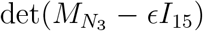 where *I*_15_ is the identity matrix of size 15. Next, form the matrix 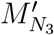 whose first column is given by 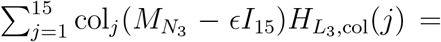 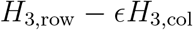 and its remaining columns are the remaining columns (i.e., columns 2 to 15) of the matrix 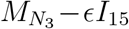. From properties of determinants, 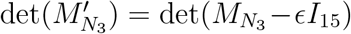 as the first column of 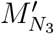 is the first column of 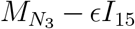 added to it a multiple of every other column of 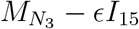 which does not alter the value of the determinant.

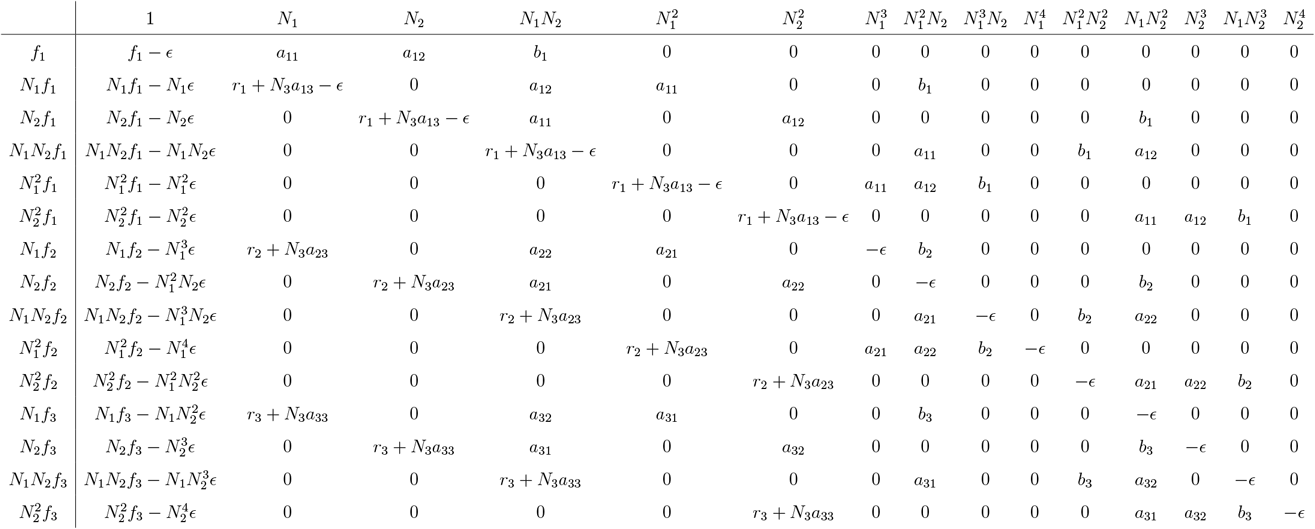

The determinant of the matrix above can be computed and the coefficient of the lowest power of *ϵ* can be extracted. Alternatively and for easier computation, let 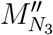 be the matrix 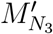 but whose first column is *H*_3,row_ instead of *H*_3,row_ − *ϵH*_3,col_. Note that 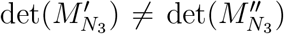, however the first non-zero coefficient of powers of *ϵ* in ascending order *ϵ* in 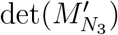 is exactly the first non-zero coefficient of powers of *ϵ* in ascending order in 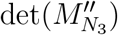. This can be proven by expanding 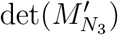 along the first column. After evaluating 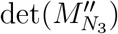, we find that the first non-zero coefficient of powers of *ϵ* in ascending order is the coefficient of *ϵ*^3^ (i.e., coefficients of *ϵ*^2^, *ϵ*^1^ and *ϵ*^0^ are all zero). This coefficient, which acts as a substitution to the resultant, can be written as *T*_31_*f*_1_ + *T*_32_*f*_2_ + *T*_33_*f*_3_ where *T*_31_, *T*_32_ and *T*_33_ have the following form.

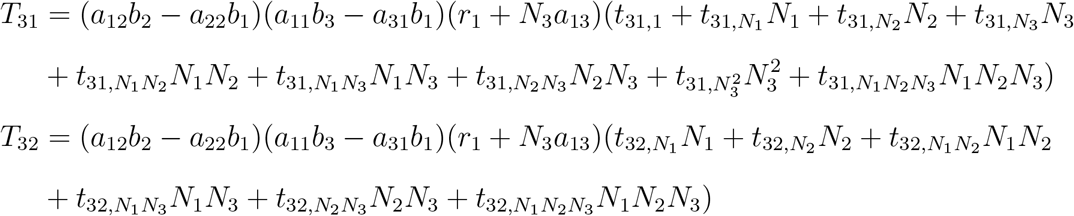

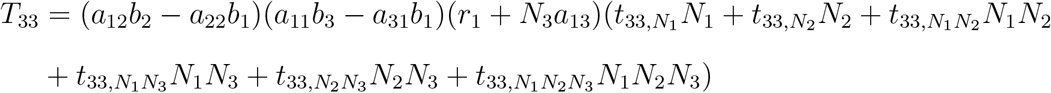

The *t*’s are polynomials in model parameters (i.e., the *r*’s, *a*’s and *b*’s) and their expressions are too large to display here. However, for illustration purposes, closed form expressions for 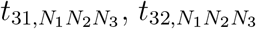 and 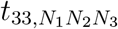 are shown below

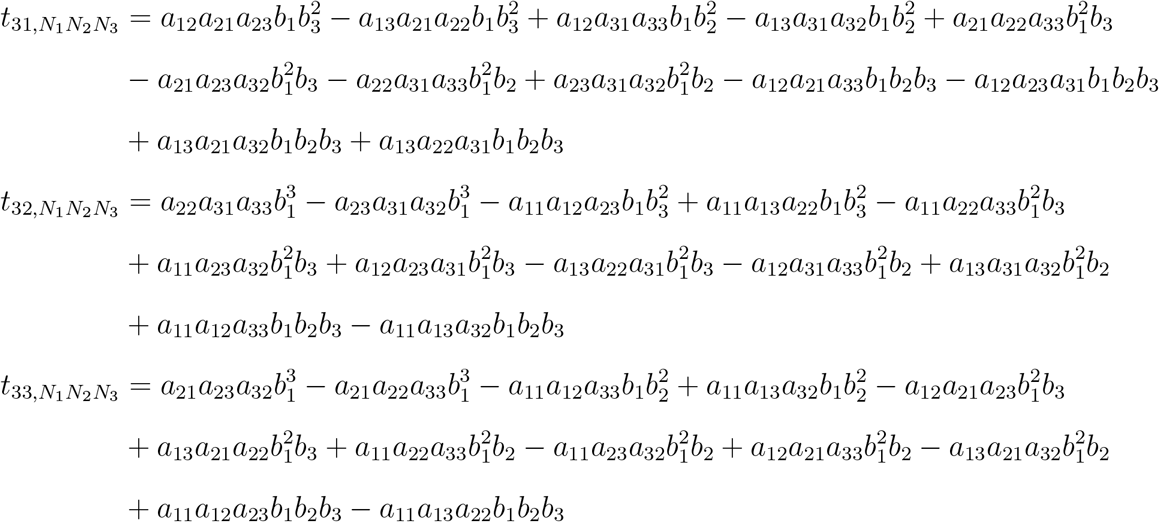

Upon substituting *f*_1_, *f*_2_ and *f*_3_ into *T*_21_*f*_1_ + *T*_22_*f*_2_ + *T*_23_*f*_3_ and simplifying the expression (or finding the coefficient of *ϵ*^3^ in the determinant of the matrix 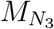 directly), we have the formula of the resultant 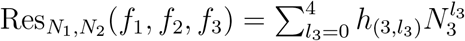 which is a polynomial of degree 4 in *N*_3_ and contains no *N*_1_’s nor *N*_2_’s. The coefficients of the resultant *h*_(3,4)_, *h*_(3,3)_, *h*_(3,2)_, *h*_(3,1)_ and *h*_(3,0)_ are too large to display here and can found via any symbolic toolbox. After finding the resultants, we evaluate *T* (*f*_1_, *f*_2_, *f*_3_) (i.e., the determinant of the eliminating matrix) as well as *J* (*f*_1_, *f*_2_, *f*_3_) (i.e., the determinant of the Jacobian of *f*_1_, *f*_2_ and *f*_3_) which are shown below:

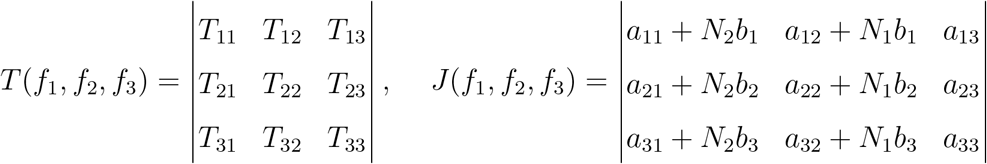

Let 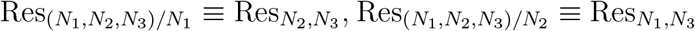 and 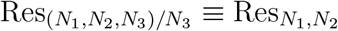. Next, expand the generating function *G*(*f*_1_, *f*_2_, *f*_3_) around *N*_1_ = ∞, *N*_2_ = ∞ and *N*_3_ = ∞. Since the three resultants are univariate polynomials in a single variable, we can expand their reciprocal individually using MATLAB’s taylor command upon substituting *N*_1_ = 1/*x, N*_2_ = 1/*y, N*_3_ = 1/*z* or via the following expression

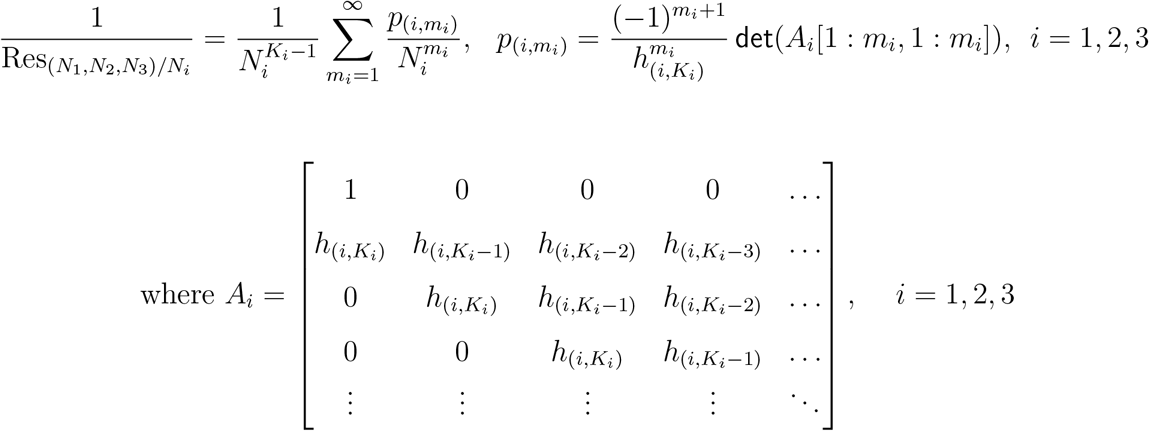

Here, *A*_*i*_[1 : *m*_*i*_, 1 : *m*_*i*_] is the sub-matrix of *A*_*i*_ that contains its first *m*_*i*_ rows and columns. After obtaining both series expansion of the resultant reciprocal, multiply the result by *T* (*f*_1_, *f*_2_, *f*_3_)*J* (*f*_1_, *f*_2_, *f*_3_) to obtain

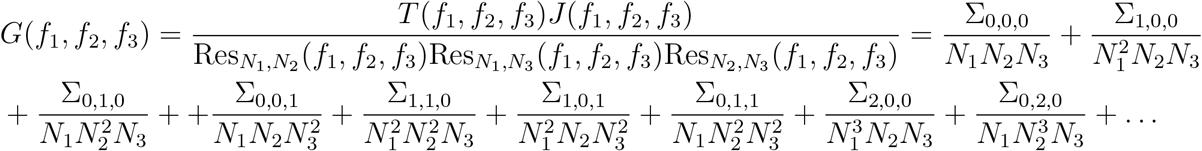

Without factorization, expressions of some of the Σ’s can extend to multiple pages. The expression for some of the lower Σ’s are shown below where 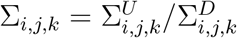 is written as a fraction of two polynomials.

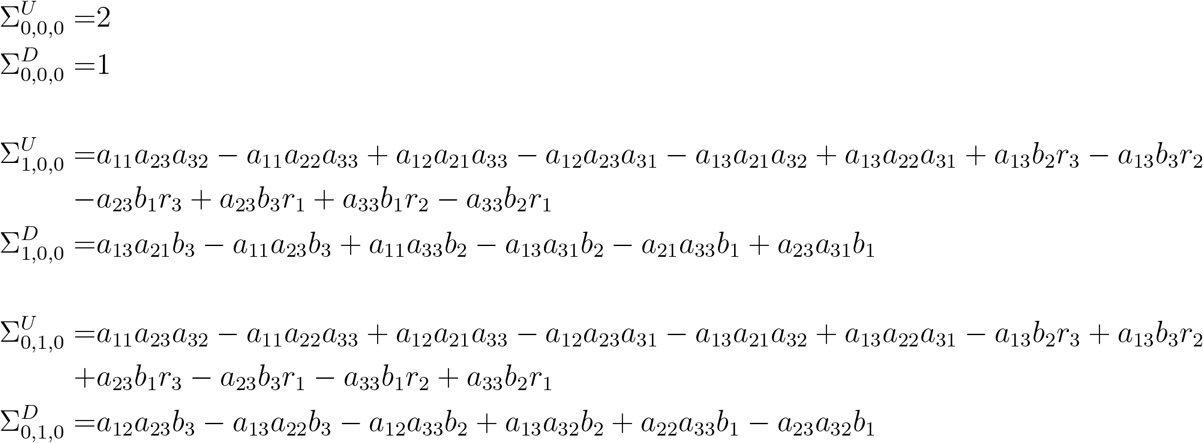

Since Σ_0,0,0_ = 2, then the system *f*_*i*_(*N*_1_, *N*_2_, *N*_3_) = 0 for *i* = 1, 2, 3 has exactly 2 complex roots. Denote to these roots by ***η***_**1**_ = [*η*_**1**,1_, *η*_**1**,2_]^*T*^, ***η***_**2**_ = [*η*_**2**,1_, *η*_**2**,2_]^*T*^ and ***η***_**3**_ = [*η*_**3**,1_, *η*_**3**,2_]^*T*^. Choose a map *m*(*N*_1_, *N*_2_, *N*_3_) = [1, *N*_1_]^*T*^ then, let *q*(*N*_1_, *N*_2_, *N*_3_) = *N*_1_*N*_2_*N*_3_ and compute *S*(*s*_1_, *s*_2_, *s*_3_) = *W* Δ*W* ^*t*^ where *W*_*ij*_ = *m*_*i*_(*η*_**1**,*j*_, *η*_2*,j*_, *η*_**3***,j*_) and Δ_*ii*_ = *q*(*η*_**1***,i*_−*s*_1_, *η*_**2***,i*_−*s*_2_, *η*_**3***,i*_−*s*_3_) is a diagonal matrix as follows.

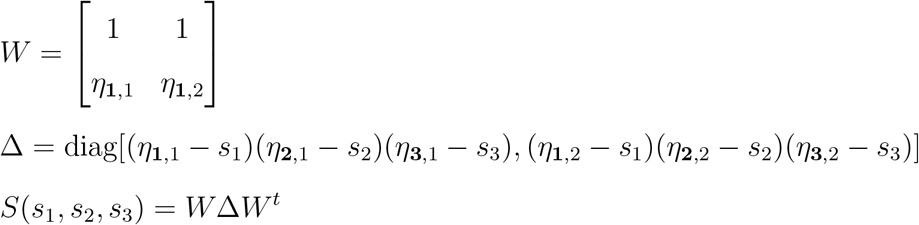

Note that 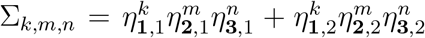 for *k, m, n* = 0, 1, 2, *…*. The components of the symmetric 2×2 matrix *S* are shown below:

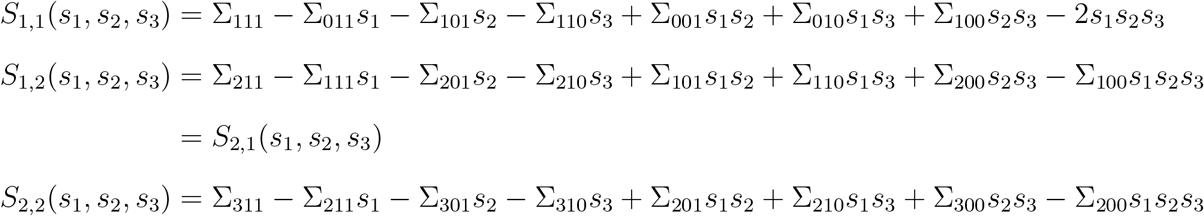

The characteristic equation of the matrix *S* is det(*S*(*s*_1_, *s*_2_, *s*_3_)) = *λ*^2^ + *v*_1_(*s*_1_, *s*_2_, *s*_3_)*λ + v*_0_(*s*_1_, *s*_2_, *s*_3_). The coefficients of the characteristic equation evaluated at (*s*_1_, *s*_2_, *s*_3_) = {(0, 0, 0), (∞, 0, 0), (0, ∞, 0), (∞, ∞, 0), (0, 0, ∞), (∞, 0, ∞), (0, ∞, ∞), (∞, ∞, ∞)} are displayed below. Note that *v*_*i*_(*m*_1_, *m*_2_, *m*_3_) where *m*_1_, *m*_2_, *m*_3_ ∈ {0, ∞} is the coefficient of 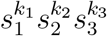 in *v*_*i*_(*s*_1_, *s*_2_, *s*_3_) where *k*_*j*_ = 0 if *m*_*j*_ = 0 and *k*_*j*_ = 2 − *i* if *m*_*j*_ = ∞ for *j* = 1, 2, 3.

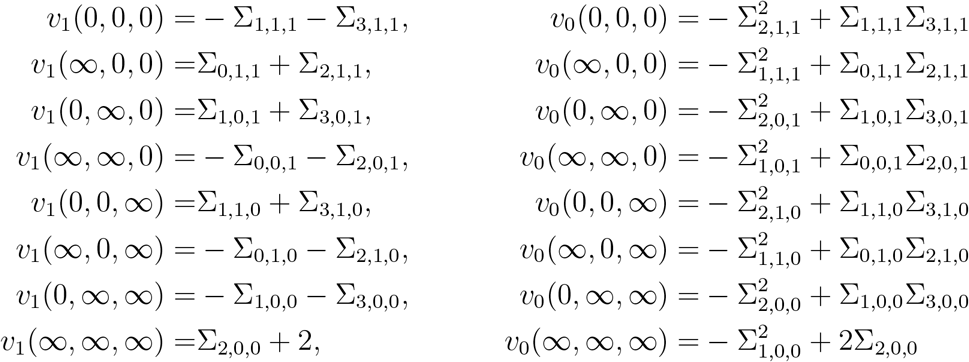

Let *V* (*a, b, c*) be the number of consecutive sign changes in [1, *v*_1_(*a, b, c*), *v*_0_(*a, b, c*)] where *a, b* and *c* are either 0 or ∞. The formula of *V* (*a, b, c*) is shown below

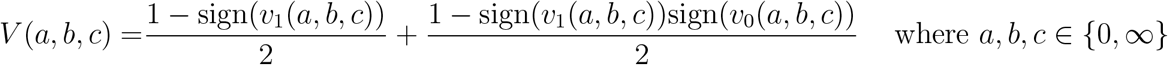

From the *V*’s, we can find the formula of the number of feasible roots of *f*_1_(*N*_1_, *N*_2_, *N*_3_), *f*_2_(*N*_1_, *N*_2_, *N*_3_) and *f*_3_(*N*_1_, *N*_2_, *N*_3_) which is given by *F* (**Ψ**) = (*V* (0, 0, 0) − *V* (∞, 0, 0) − *V* (0, ∞, 0) − *V* (0, 0, ∞) + *V* (∞, ∞, 0) + *V* (∞, 0, ∞) + *V* (0, ∞, ∞) − *V* (∞, ∞, ∞))/4. Let us consider the parameter **Ψ** = (*r*_1_, *r*_2_, *r*_3_, *a*_11_, *a*_12_, *a*_13_, *a*_21_, *a*_22_, *a*_23_, *a*_31_, *a*_32_, *a*_33_, *b*_1_, *b*_2_, *b*_3_) = (0.5, −1.5, −0.5, 0.5, −1.5, −0.5, *a*_21_, 2.6, −5, −0.5, −10, 1, 0.2, −0.1, *b*_3_) where the parameters *a*_21_ ∈ [−7, −1] and *b*_3_ ∈ [1.5, 5] are restricted, we find that feasibility (i.e., *F* (**Ψ**) ≥ 1) can only be satisfied under the two condition that are shown below:

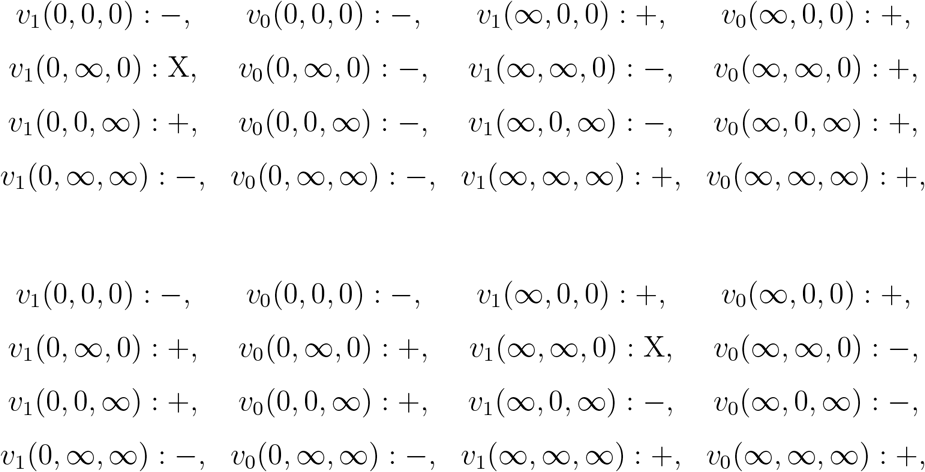

When we plot the sign of each of the quantities (i.e., the *v*_*i*_’s) in the two conditions above, we find that feasibility is satisfied if any of the following four conditions hold: *v*_1_(0, 0, 0) < 0,*v*_0_(0, 0, 0) < 0, *v*_1_(∞, 0, 0) > 0 or *v*_0_(∞, 0, 0) > 0 which are equivalent to each other in the domain prescribed by **Ψ**. Note that these four inequalities are shared among the two conditions described above. In addition, note that the simplest inequality among those (i.e., with lowest symmetric sums) is *v*_1_(∞, 0, 0) > 0. In the next plots, we plot the sign of *v*_1_(∞, 0, 0) and verify that it matches the feasibility region given by *F* (**Ψ**) which verifies the correctness of our methodology.

## S6 Ex 4: 3-Species with Higher-Order Interactions

Consider Lotka-Volterra model with higher-order interactions that is shown below:

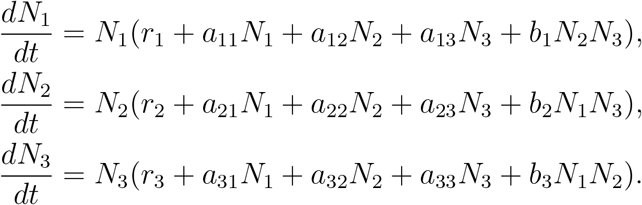

To study feasibility, the polynomials that are needed to be considered are *f*_1_(*N*_1_, *N*_2_, *N*_3_) = *r*_1_ + *a*_11_*N*_1_ + *a*_12_*N*_2_ + *a*_13_*N*_3_ + *b*_1_*N*_2_*N*_3_, *f*_2_(*N*_1_, *N*_2_, *N*_3_) = *r*_2_ + *a*_21_*N*_1_ + *a*_22_*N*_2_ + *a*_23_*N*_3_ + *b*_2_*N*_1_*N*_3_ and *f*_3_(*N*_1_, *N*_2_, *N*_3_) = *r*_3_+*a*_31_*N*_1_+*a*_32_*N*_2_+*a*_33_*N*_3_+*b*_3_*N*_1_*N*_2_. Next, assume that *N*_1_ is constant and homogenize *f*_1_, *f*_2_ and *f*_3_ with a forth variable *W* as follows:

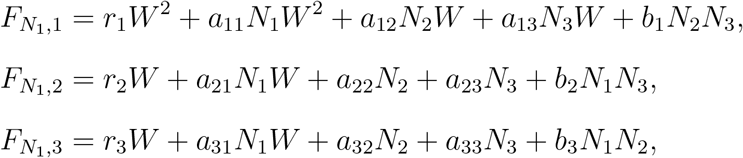

Note that the total degree of each of 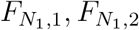 and 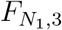 (or the total degree of *f*_1_, *f*_2_ and *f*_3_ assuming *N*_1_ is a constant) is *d*_1,1_ = 2, *d*_1,2_ = 1 and *d*_1,3_ = 1 respectively. From the *d*’s, we compute 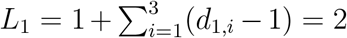. Now, we form the monomial set *H*_1_, which is a union of three disjoint monomials 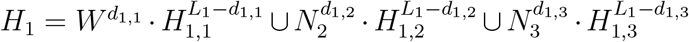 where none of these *H*’s involve *N*_1_ and each is indicated below in curly brackets:

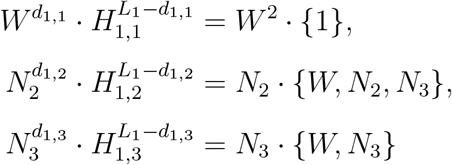

Form the monomial set 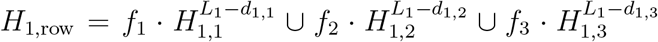 evaluated at *W* = 1 that is shown below. In addition, form the monomial set *H*_1,col_ which is simply *H*_1_ evaluated at *W* = 1 to get

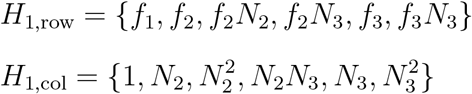

After that, form the Macaulay matrix 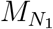 which is a square matrix whose size is 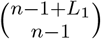 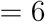. The (*i, j*) entry of the Macaulay matrix is the coefficient of *H*_1,col_(*j*) in the expression of *H*_1,row_(*i*) assuming that *N*_1_ is a constant. For example, the (3, 2) entry in the matrix is the coefficient of *N*_2_ in *N*_2_*f*_2_ which is *r*_2_ + *N*_1_*a*_21_. The matrix 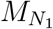 is shown below:

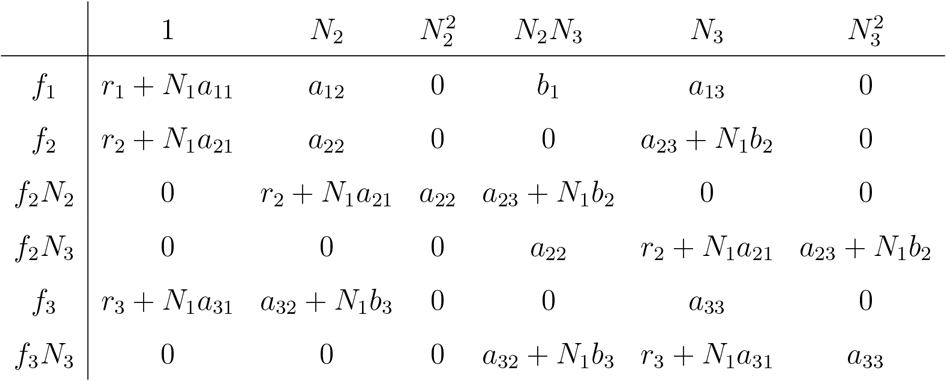

Next, form the matrix 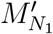 whose first column is *H*_1,row_ and its remaining columns are the remaining columns (i.e., columns 2 to 6) of the matrix 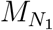 (i.e., replace the first column of 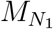 whose top header is 1 with the leftmost column which contains the *f*’s). From the formula of 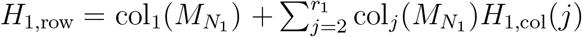, we can see that *H*_1,row_ is the first column of 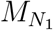 added to it a multiple of every other column of 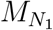, implying that 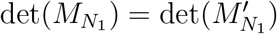. This determinant (i.e., 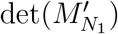) can be written as *T*_11_*f*_1_ + *T*_12_*f*_2_ + *T*_13_*f*_3_ which is shown below

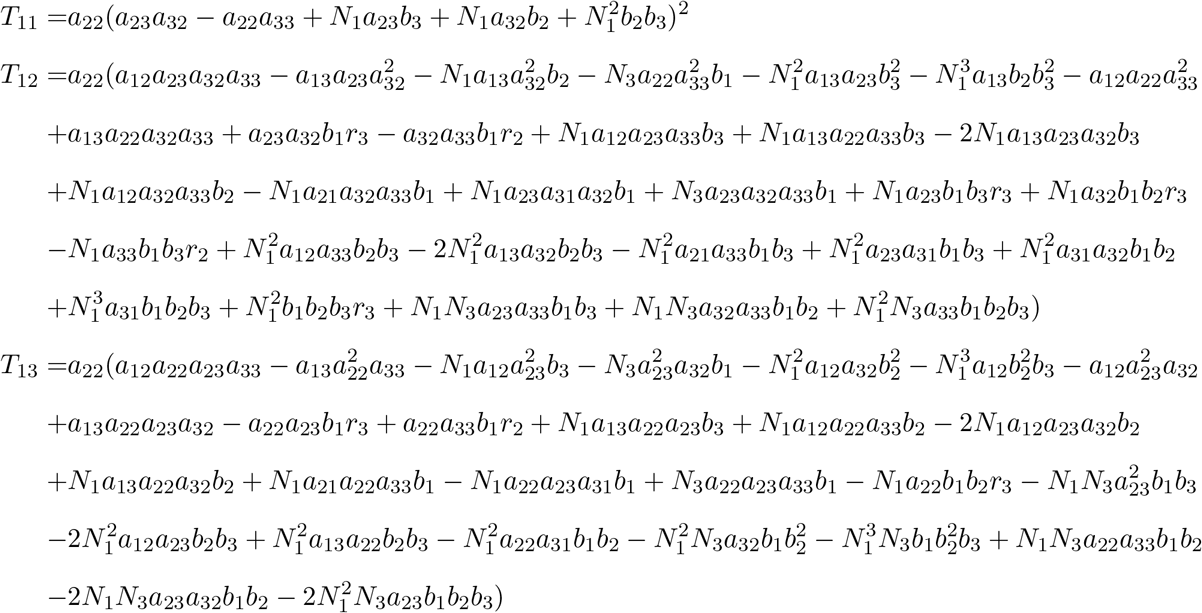

Upon substituting *f*_1_, *f*_2_ and *f*_3_ into *T*_11_*f*_1_ + *T*_12_*f*_2_ + *T*_13_*f*_3_ and simplify the expression, we have the formula of the resultant 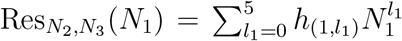 which is a polynomial of degree 5 in *N*_1_ and contains no *N*_2_’s nor *N*_3_’s. The six coefficients of the resultant *h*_(1,5)_, *h*_(1,4)_, …, *h*_(1,0)_ are shown below and none of them contain any of the *N*’s.

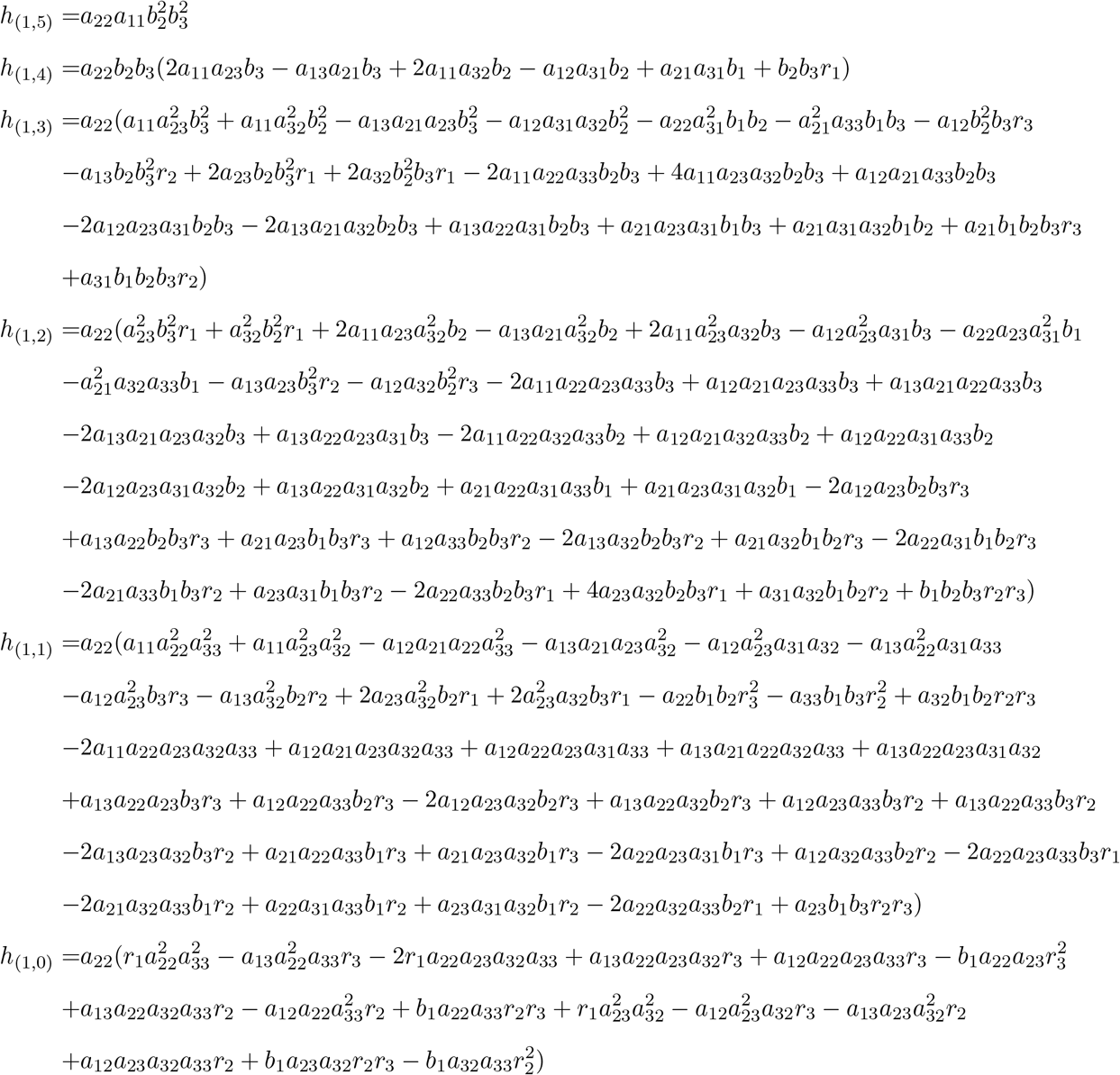

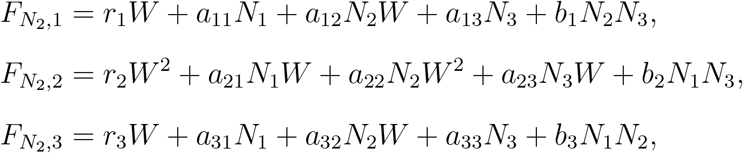

Note that the total degree of each of 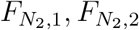 and 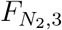 (or the total degree of *f*_1_, *f*_2_ and *f*_3_ assuming *N*_2_ is a constant) is *d*_2,1_ = 1, *d*_2,2_ = 2 and *d*_2,3_ = 1 respectively. From the *d*’s, we compute 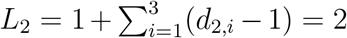. Now, we form the monomial set *H*_2_, which is a union of three disjoint monomials 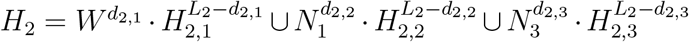 where none of these *H*’s involve *N*_2_ and each is indicated below:

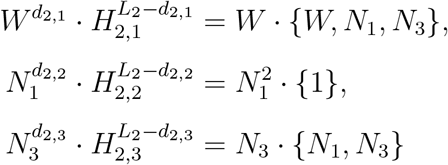

Next, form the monomial set 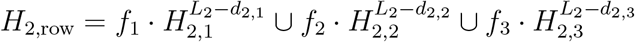 evaluated at *W* = 1 that is shown below. In addition, form the monomial set *H*_2,col_ which is simply *H*_2_ evaluated at *W* = 1 to get

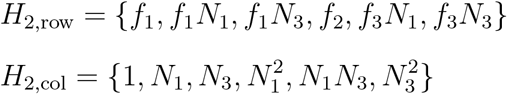

After that, form the Macaulay matrix 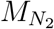 which is a square matrix whose size 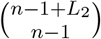 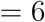. The (*i, j*) entry of the Macaulay matrix is the coefficient of *H*_2,col_(*j*) in the expression of *H*_2,row_(*i*) assuming that *N*_2_ is a constant. The matrix 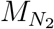 is shown below:

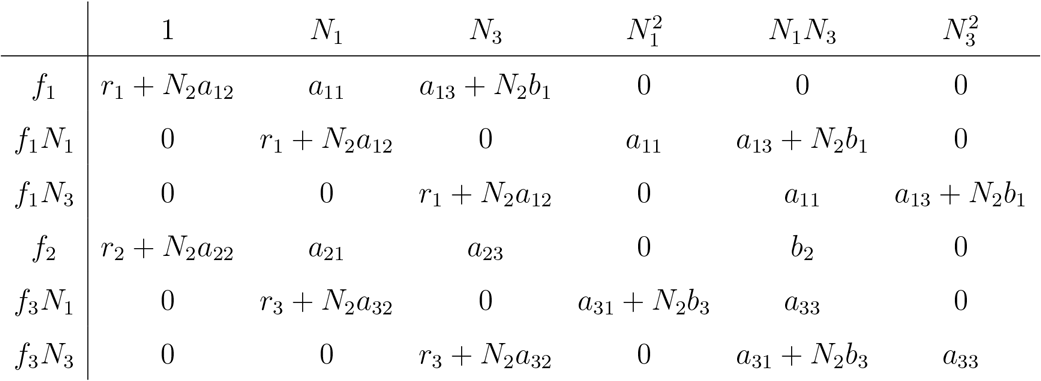

Next, form the matrix 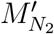 whose first column is *H*_2,row_ and its remaining columns are the remaining columns (i.e., columns 2 to 6) of the matrix 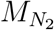 (i.e., replace the first column of 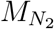 whose top header is 1 with the leftmost column which contains the *f*’s). Again, from the formula of 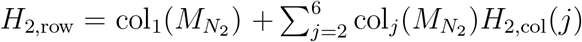, we can see that *H*_2,row_ is the first column of 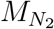 added to it a multiple of every other column of 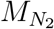, implying that 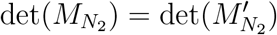. This determinant (i.e., 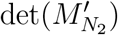) can be written as *T*_21_*f*_1_ + *T*_22_*f*_2_ + *T*_23_*f*_3_. The expressions of *T*_21_ and *T*_23_ are too large to be displayed here, however, their forms are shown below:

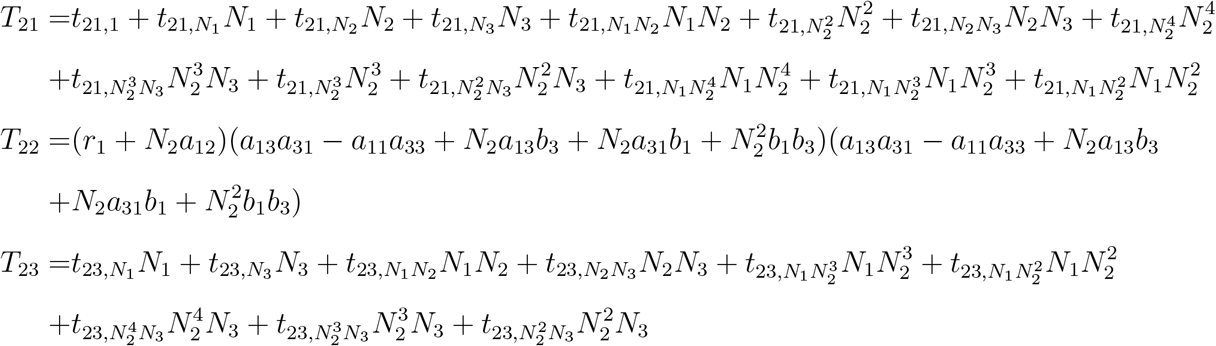

Again, the *t*’s are polynomials in model parameters (i.e., the *r*’s, *a*’s and *b*’s). For illustration purposes, closed form expressions for 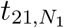 and 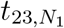 are shown below:

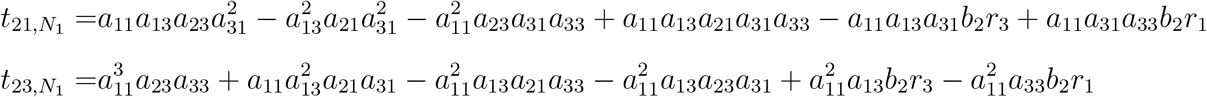

Upon substituting *f*_1_, *f*_2_ and *f*_3_ into *T*_21_*f*_1_ + *T*_22_*f*_2_ + *T*_23_*f*_3_ and simplify the expression, we have the formula of the resultant 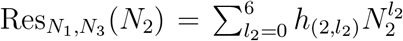 which is a polynomial of degree 6 in *N*_2_ and contains no *N*_1_’s nor *N*_3_’s. The seven coefficients of the resultant *h*_(2,6)_, *h*_(2,5)_, …, *h*_(2,0)_ are shown below and none of them contain any of the *N*’s.

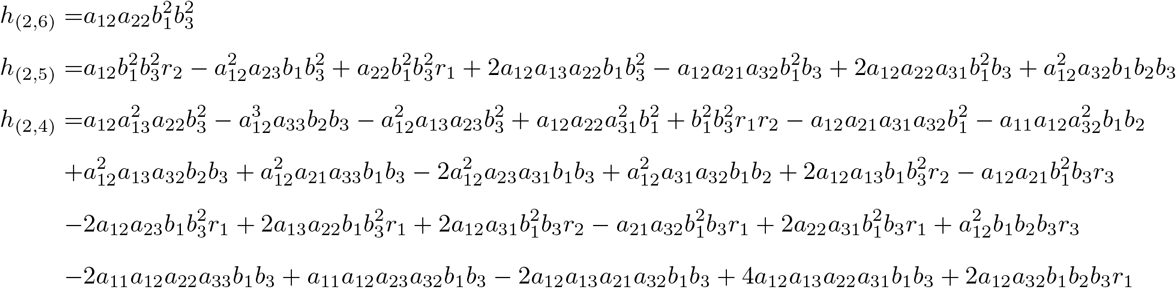

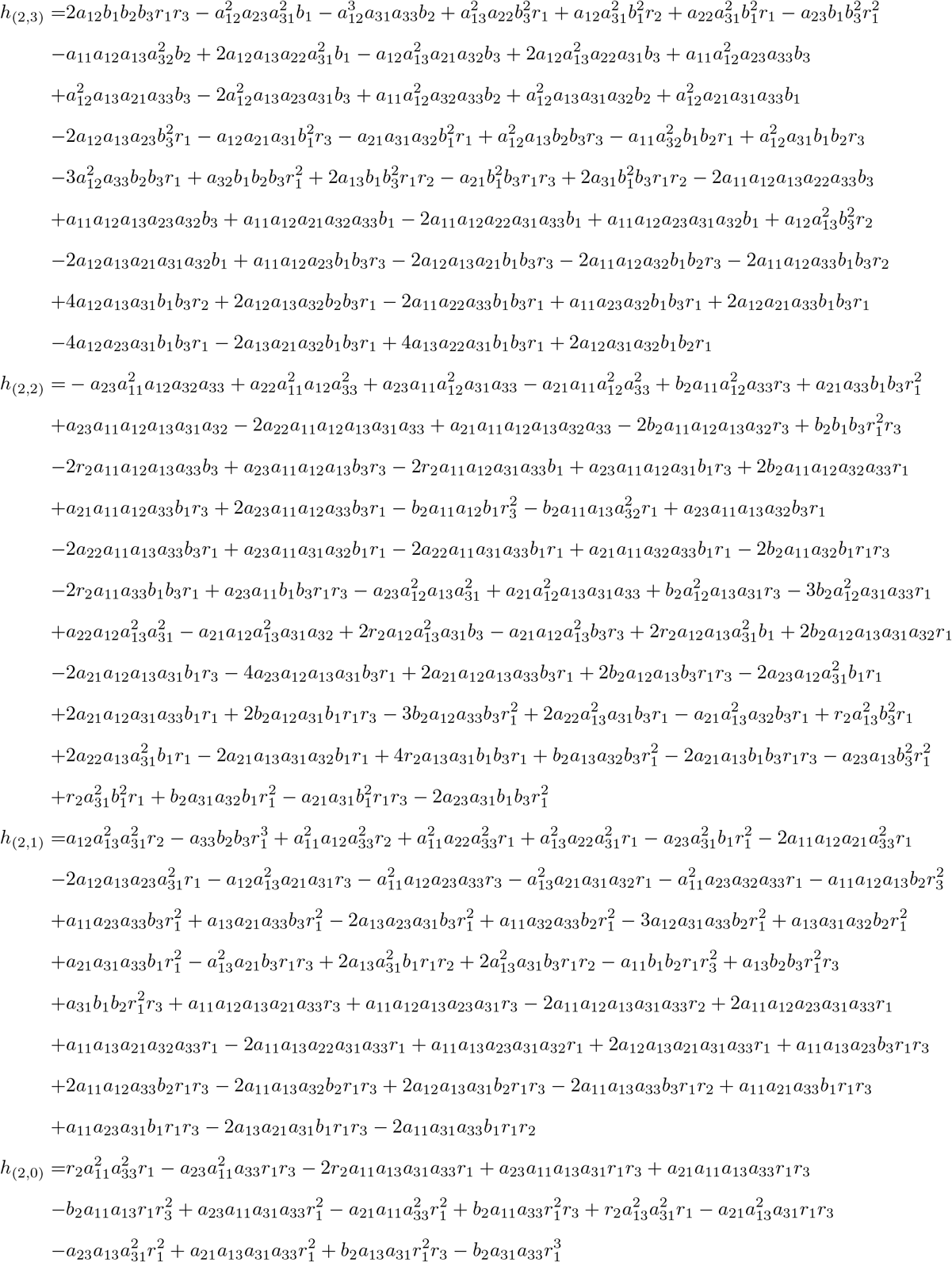

Next, assume that *N*_3_ is constant and homogenize *f*_1_, *f*_2_ and *f*_3_ with a forth variable *W* :

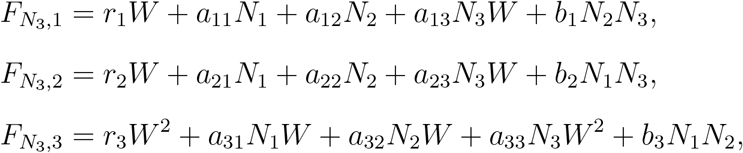

Note that the total degree of each of 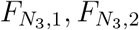 and 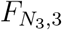 (or the total degree of *f*_1_, *f*_2_ and *f*_3_ assuming *N*_3_ is a constant) is *d*_3,1_ = 1, *d*_3,2_ = 1 and *d*_3,3_ = 2 respectively. From the *d*’s, we compute 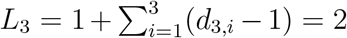. Now, we form the monomial set *H*_3_, which is a union of three disjoint monomials 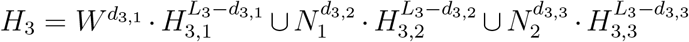 where none of these *H*’s involve *N*_3_ and each is indicated below:

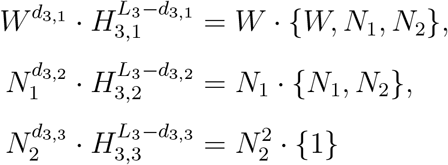

Next, form the monomial set 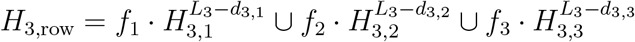 evaluated at *W* = 1 that is shown below. In addition, form the monomial set *H*_3,col_ which is simply *H*_3_ evaluated at *W* = 1 to get

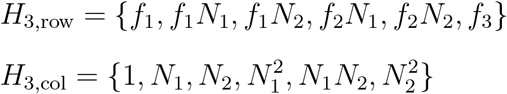

After that, form the Macaulay matrix 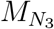 which is a square matrix whose size is 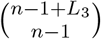 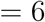. The (*i, j*) entry of the Macaulay matrix is the coefficient of *H*_3,col_(*j*) in the expression of *H*_3,row_(*i*) assuming that *N*_3_ is a constant. The matrix 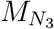 is shown below:

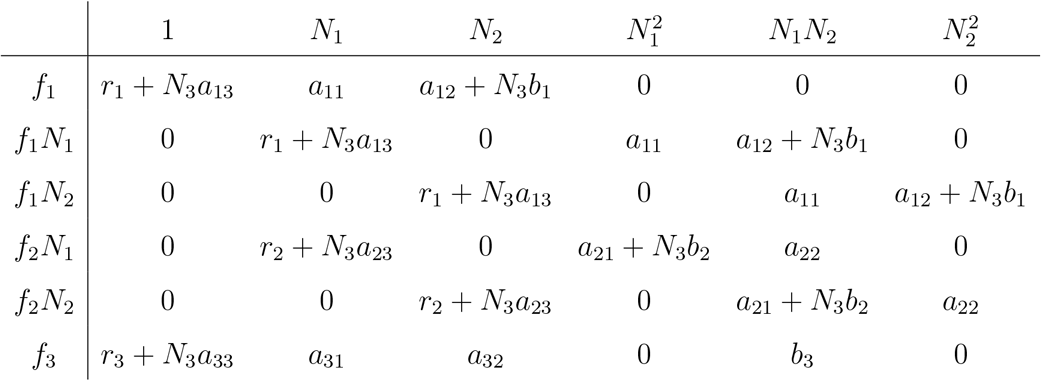

Next, form the matrix 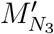 whose first column is *H*_3,row_ and its remaining columns are the remaining columns (i.e., columns 2 to 6) of the matrix 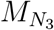 (i.e., replace the first column of 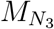 whose top header is 1 with the leftmost column which contains the *f*’s). Again, from the formula of 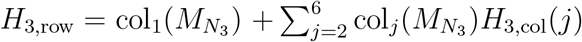, we can see that *H*_3,row_ is the first column of 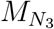 added to it a multiple of every other column of 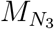, implying that 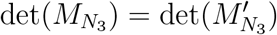. This determinant (i.e., 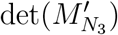) can be written as *T*_31_*f*_1_ + *T*_32_*f*_2_ + *T*_33_*f*_3_. The expressions of *T*_21_ and *T*_23_ are too large to be displayed here, however, their forms are shown below:

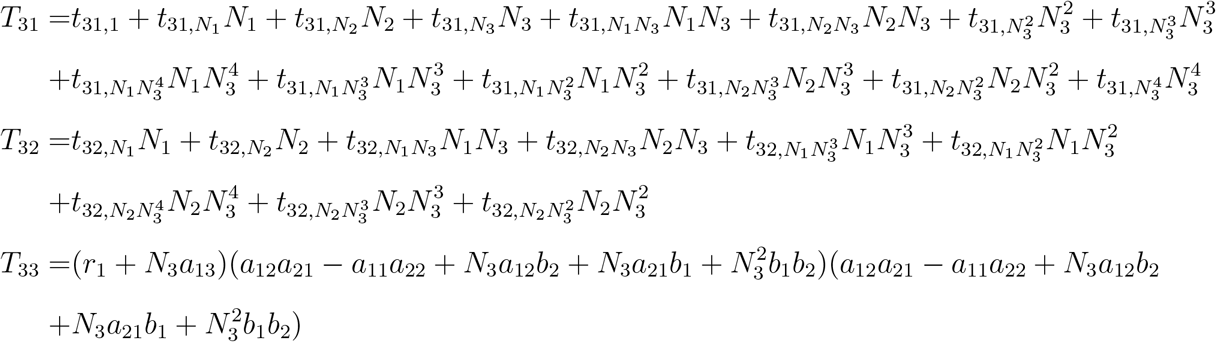

Again, the *t*’s are polynomials in model parameters (i.e., the *r*’s, *a*’s and *b*’s). For illustration purposes, closed form expressions for 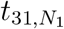 and 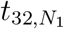 are shown below:

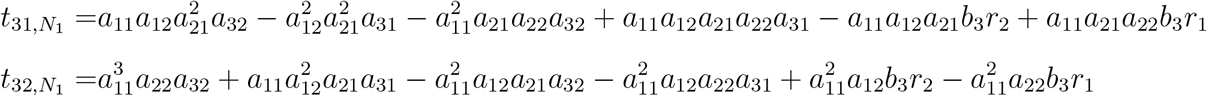

Upon substituting *f*_1_, *f*_2_ and *f*_3_ into *T*_31_*f*_1_ + *T*_32_*f*_2_ + *T*_33_*f*_3_ and simplifying the expression, we have the formula of the resultant 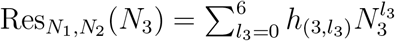 which is a polynomial of degree 6 in *N*_3_ and contains no *N*_1_’s nor *N*_2_’s. The seven coefficients are shown below

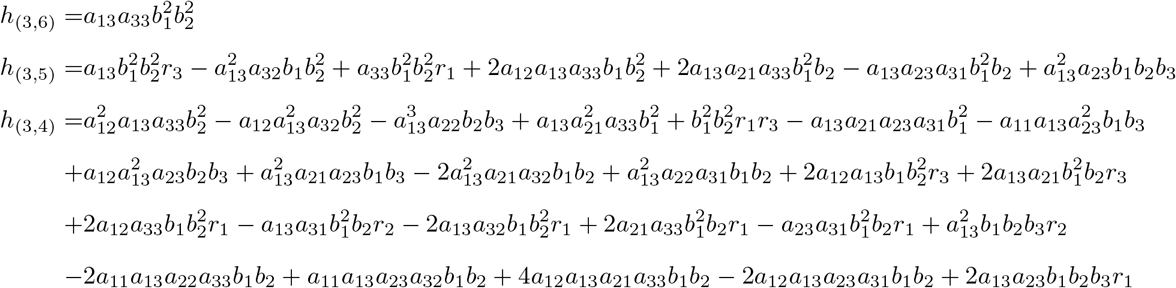

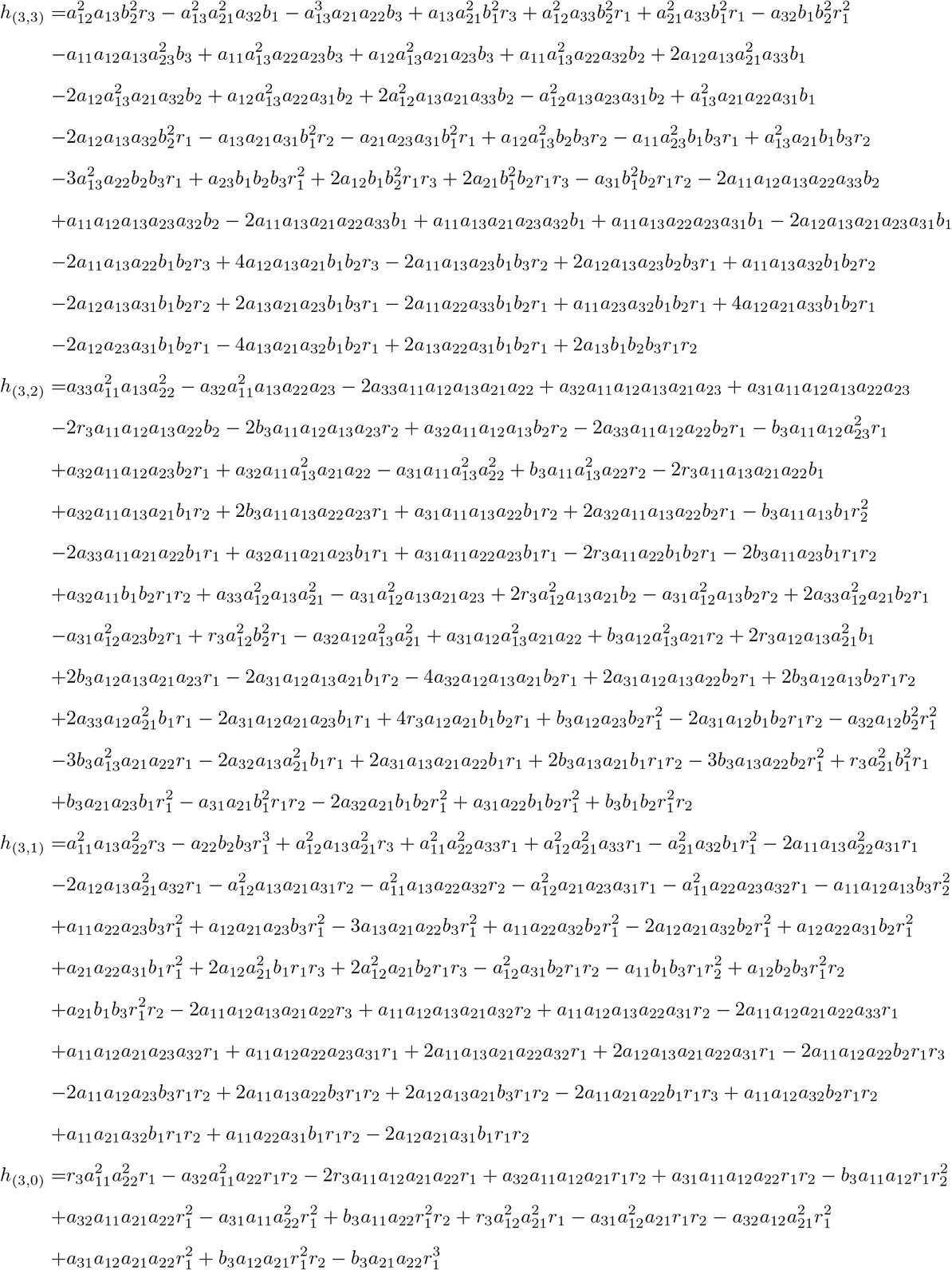

After finding the resultants, we evaluate *T* (*f*_1_, *f*_2_, *f*_3_) (i.e., the determinant of the eliminating matrix) as well as *J* (*f*_1_, *f*_2_, *f*_3_) (i.e., the determinant of the Jacobian of *f*_1_, *f*_2_ and *f*_3_) which are shown below:

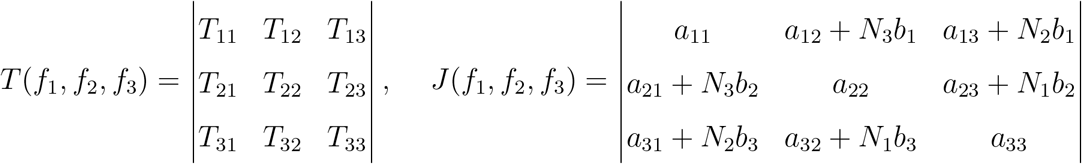

Obtain the series expansion of the reciprocal of each resultant individually then multiply the results by *T* (*f*_1_, *f*_2_, *f*_3_)*J* (*f*_1_, *f*_2_, *f*_3_) to obtain

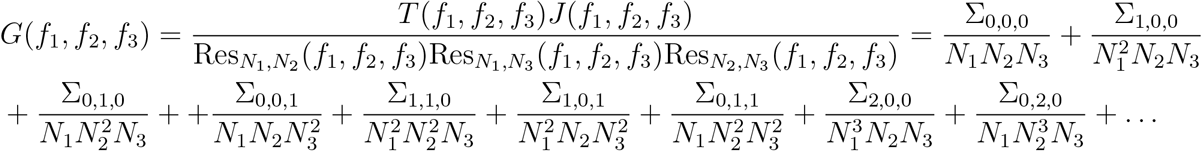

Without factorization, expressions of some of the Σ’s can extend to multiple pages. The expression for some of the lower Σ’s are shown below where 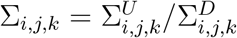 is written as a fraction of two polynomials.

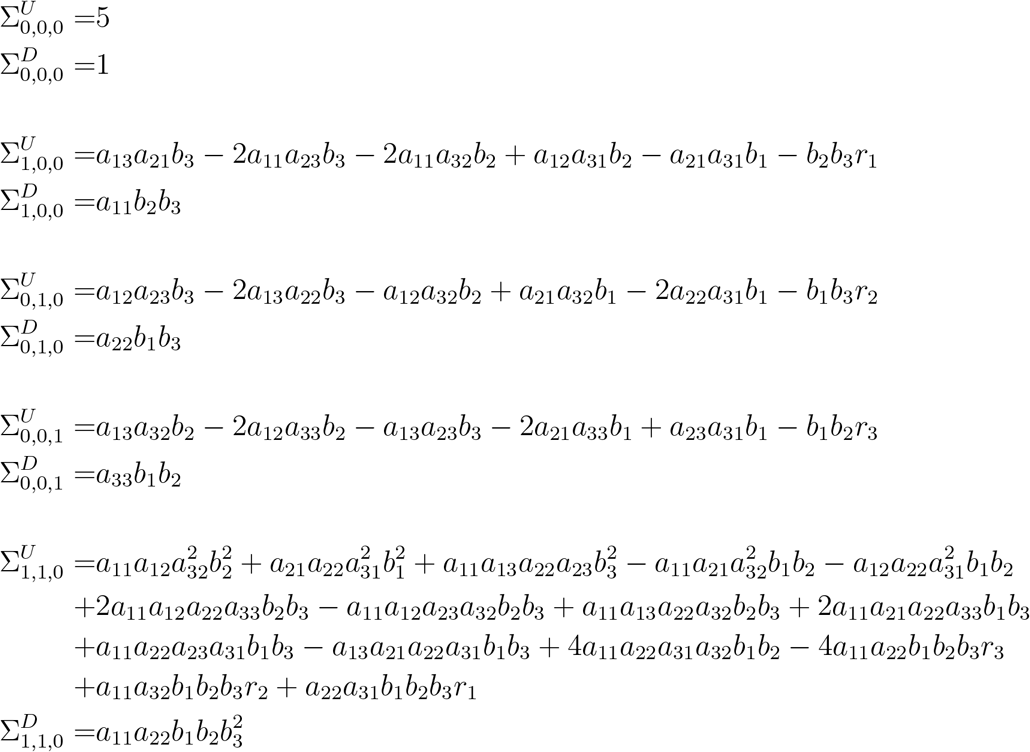

Observe that 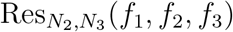 is a polynomial of degree 5 in *N*_1_ only and thus cannot be solved analytically. Similarly, 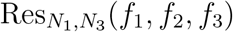 and 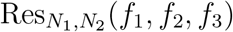 are polynomials of degree 6 i in *N*_2_ and *N*_3_. Note that the roots of the three resultants, upon appropriate pairing of roots of each of them, are the roots of the system *f*_*i*_(*N*_1_, *N*_2_, *N*_3_) = 0 for *i* = 1, 2, 3. From Abel’s impossibility theorem, since it is impossible to solve for the roots of a quintic or higher degree polynomials in terms of radicals, then the roots of any of the resultants are unattainable analytically which implies that the system *f*_*i*_(*N*_1_, *N*_2_, *N*_3_) = 0 cannot be solved analytically. Since Σ_0,0,0_ = 5, then the system *f*_*i*_(*N*_1_, *N*_2_, *N*_3_) = 0 for *i* = 1, 2, 3 has exactly 5 complex roots. Denote to them by *η*_**1**_ = [*η*_**1**,1_, *η*_**1**,2_, …, *η*_**1**,5_]^*T*^, ***η***_**2**_ = [*η*_**2**,1_, *η*_**2**,2_, …, *η*_**2**,5_]^*T*^ and ***η***_**3**_ = [*η*_**3**,1_, *η*_**3**,2_, …, *η*_**3**,5_]^*T*^. Choose a map *m*(*N*_1_, *N*_2_, *N*_3_) = [1, *N*_1_, *N*_1_*N*_2_, *N*_1_*N*_3_, *N*_1_*N*_2_*N*_3_]^*T*^. Note that if we choose a lower order map such as *m*(*N*_1_, *N*_2_, *N*_3_) = [1, *N*_1_, *N*_2_, *N*_3_, *N*_1_*N*_2_]^*T*^, one or more coefficients of the characteristic equation of *S* that will be shown in the following pages will vanish; thus a higher-order map is needed. Next, let *Q*(*N*_1_, *N*_2_, *N*_3_) = *N*_1_*N*_2_*N*_3_ and compute *S*(*s*_1_, *s*_2_, *s*_3_) = *W* Δ*W* ^*t*^ where *W*_*ij*_ = *m*_*i*_(*η*_**1***,j*_, *η*_**2***,j*_, *η*_**3***,j*_) and Δ_*ii*_ = *Q*(*η*_**1***,i*_ − *s*_1_, *η*_**2***,i*_ − *s*_2_, *η*_**3***,i*_ − *s*_3_) is a diagonal matrix as follows.

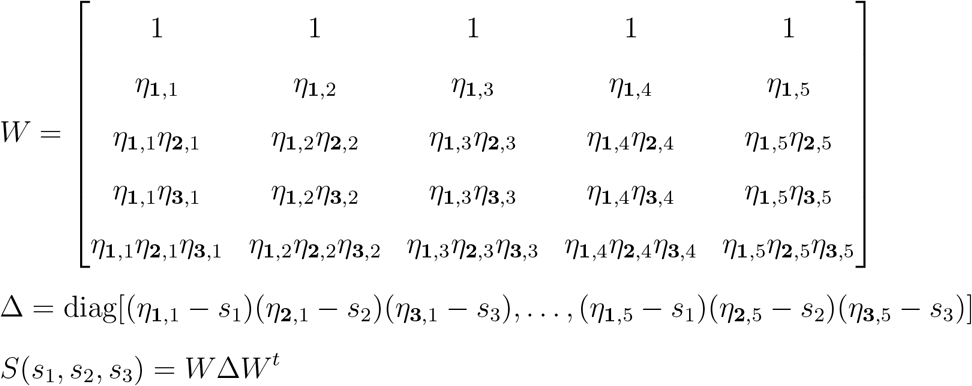

Note that 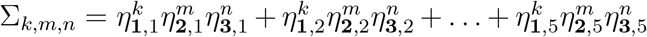 for *k, m, n* = 0, 1, 2, The components of the symmetric 5×5 matrix *S* are shown below:

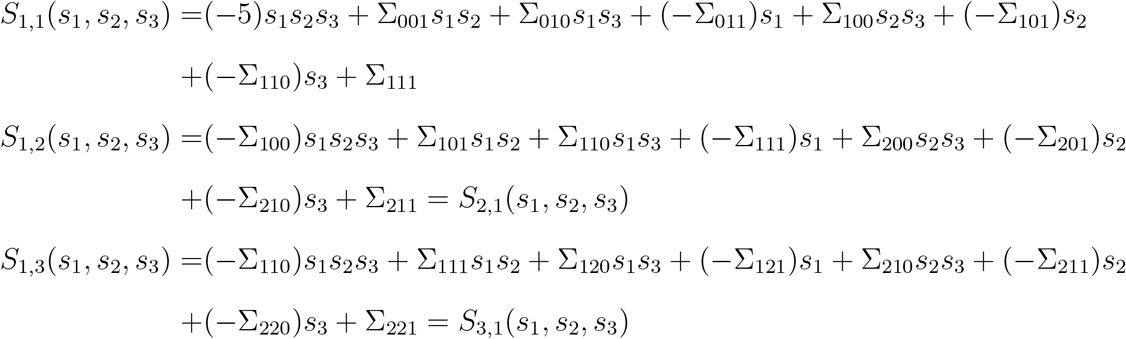

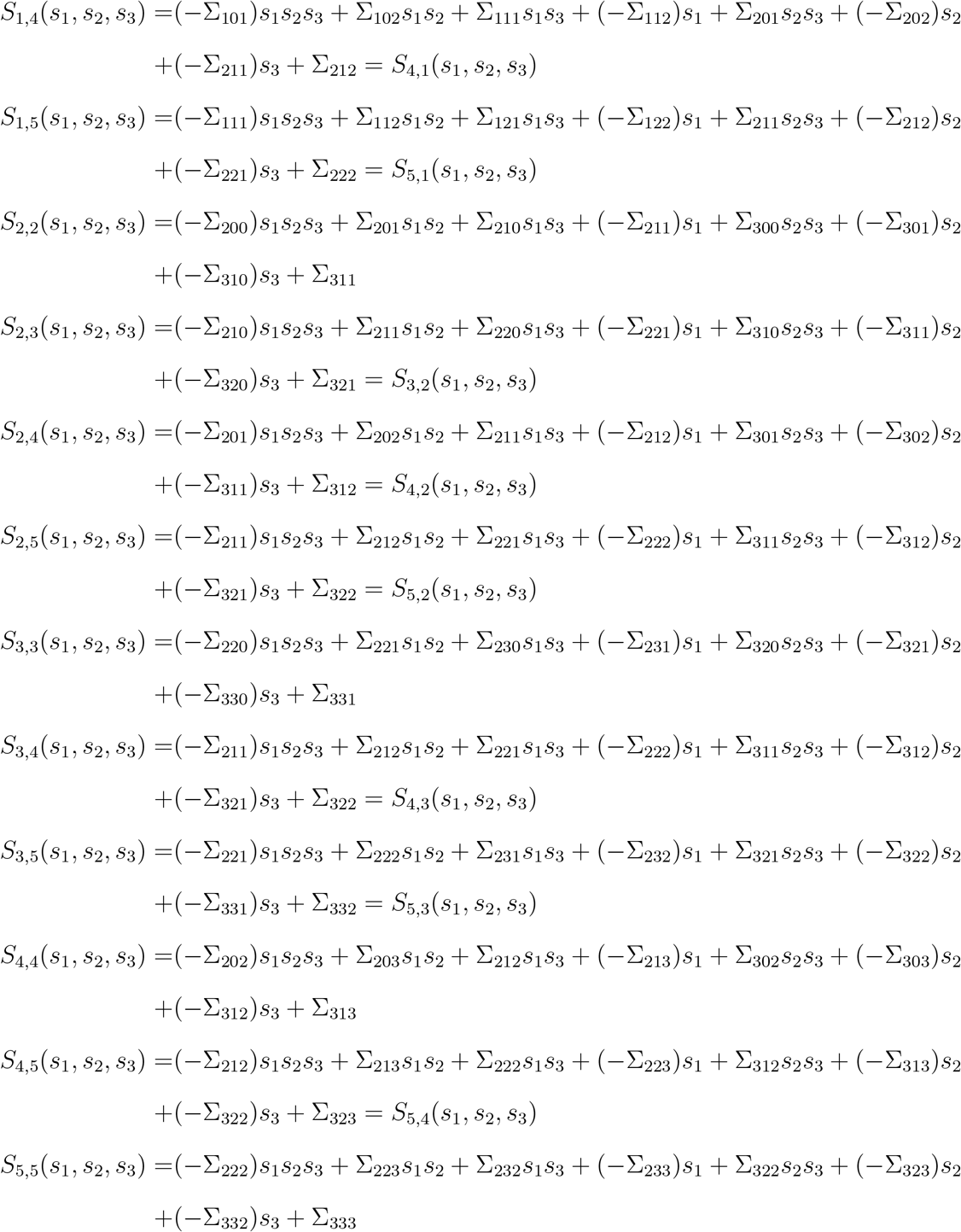

The characteristic equation of the matrix *S* is simply det(*S*(*s*_1_, *s*_2_, *s*_3_)) = *λ*^5^+*v*_4_(*s*_1_, *s*_2_, *s*_3_)*λ*^4^+ *v*_3_(*s*_1_, *s*_2_, *s*_3_)*λ*^3^ + *v*_2_(*s*_1_, *s*_2_, *s*_3_)*λ*^2^ + *v*_1_(*s*_1_, *s*_2_, *s*_3_)*λ + v*_0_(*s*_1_, *s*_2_, *s*_3_). The coefficients of the characteristic equation need to be evaluated at (*s*_1_, *s*_2_, *s*_3_) = {(0, 0, 0), (∞, 0, 0), (0, ∞, 0), (∞, ∞, 0), (0, 0, ∞), (∞, 0, ∞), (0, ∞, ∞), (∞, ∞, ∞)}. Note that *v*_*i*_(*m*_1_, *m*_2_, *m*_3_) where *m*_1_, *m*_2_, *m*_3_ ∈ {0, ∞} is the coefficient of 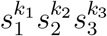 in *v*_*i*_(*s*_1_, *s*_2_, *s*_3_) where *k*_*j*_ = 0 if *m*_*j*_ = 0 and *k*_*j*_ = 5 − *i* if *m*_*j*_ = ∞ for *j* = 1, 2, 3. Since some of these 40 quantities are large, we will omit writing them. Next, let *V* (*a, b, c*) be the number of consecutive sign changes in [1, *v*_1_(*a, b, c*), *v*_0_(*a, b, c*)] where *a, b* and *c* are either 0 or ∞. The formula of *V* (*a, b, c*) is shown below

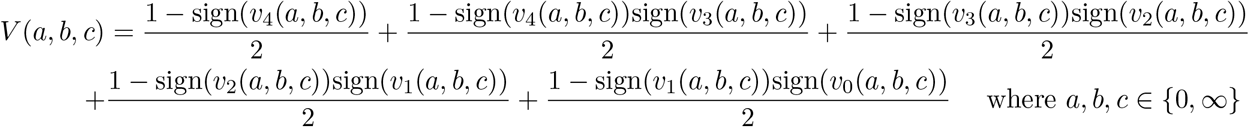

From the *V*’s, we can find the formula of the number of feasible roots of *f*_1_(*N*_1_, *N*_2_, *N*_3_), *f*_2_(*N*_1_, *N*_2_, *N*_3_) and *f*_3_(*N*_1_, *N*_2_, *N*_3_) which is given by *F* (**Ψ**) = (*V* (0, 0, 0) − *V* (∞, 0, 0) − *V* (0, ∞, 0) − *V* (0, 0, ∞) + *V* (∞, ∞, 0) + *V* (∞, 0, ∞) + *V* (0, ∞, ∞) − *V* (∞, ∞, ∞))/4. Let us consider the parameter **Ψ** = (*r*_1_, *r*_2_, *r*_3_, *a*_11_, *a*_12_, *a*_13_, *a*_21_, *a*_22_, *a*_23_, *a*_31_, *a*_32_, *a*_33_, *b*_1_, *b*_2_, *b*_3_) = (1.5, −1.5, −1.5, 2, −1.5, −1.5, *a*_21_, 2, −1.5, −1.5, −1, 1, 1, −1, *b*_3_) where the parameters *a*_21_ ∈ [1, 6] and *b*_3_ ∈ [2, 5] are restricted. We find that feasibility (i.e., *F* (**Ψ**) ≥ 1) is described by the signs of the *v*_0_’s. In particular, if any of the conditions below is satisfied, feasibility is guaranteed, which is evident from plotting any of the quantities below (except *v*_0_(0, 0, 0) < 0 (see below).

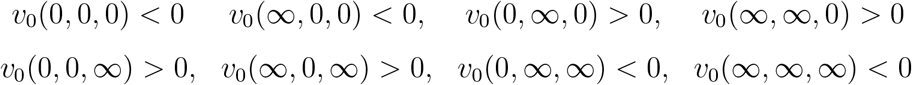

When we plot *F* (**Ψ**), we find that in some regions, some non-integer values between 0 and 2 are output due to numerical error or *m*(*N*_1_, *N*_2_, *N*_3_) having lower order monomial maps. However, we rectified the error quickly via assigning non-integer values to their closest integers. After the rectification process, we obtained a feasibility domain plot that matches the one obtained from simulations (i.e., counting the number of feasible equilibrium points via solving the isocline equations numerically). There was no need to perform any numerical corrections when we plot the sign of *v*_0_(0, ∞, 0) or any of the 8 inequalities above (except *v*_0_(0, 0, 0) < 0) and we see that it matches the feasibility domain as shown in the plots in the following page. When we plot *v*_0_(0, 0, 0) < 0, its shape has clearly the shape of the feasibility domain but has errors that are rectifiable.

**Supplementary Figure S2:**
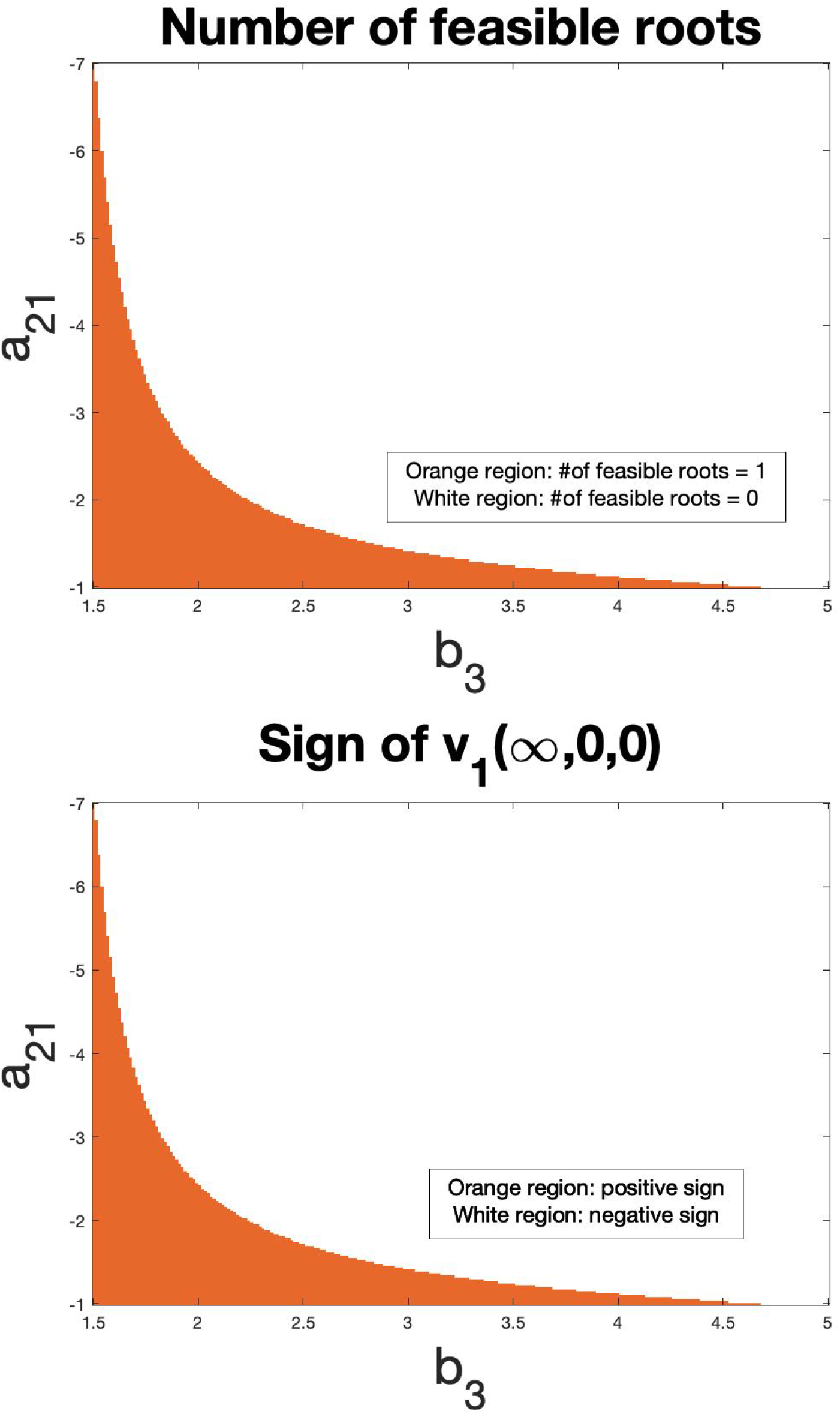
The top figure shows the number of feasible roots *F* in Lotka-Volterra model with simple higher-order terms where (*r*_1_, *r*_2_, *r*_3_, *a*_11_, *a*_12_, *a*_13_, *a*_22_, *a*_23_, *a*_31_, *a*_32_, *a*_33_, *b*_1_, *b*_2_) = (0.5, −1.5, −0.5, 0.5, −1.5, −0.5, 2.6, −5, −0.5, −10, 1, 0.2, −0.1,), *a*_21_ ∈ [−7, −1] and *b*_3_ ∈ [1.5, 5]. The bottom figure shows the sign of *v*_1_(∞, 0, 0) with the same model and parameter values and ranges. Both figures confirm that *F* > 0 when *v*_1_(, 0, 0) > 0. Simulations done via solving the isocline equations numerically and checking for the feasibility of roots match the two figures displayed here.

**Supplementary Figure S3:**
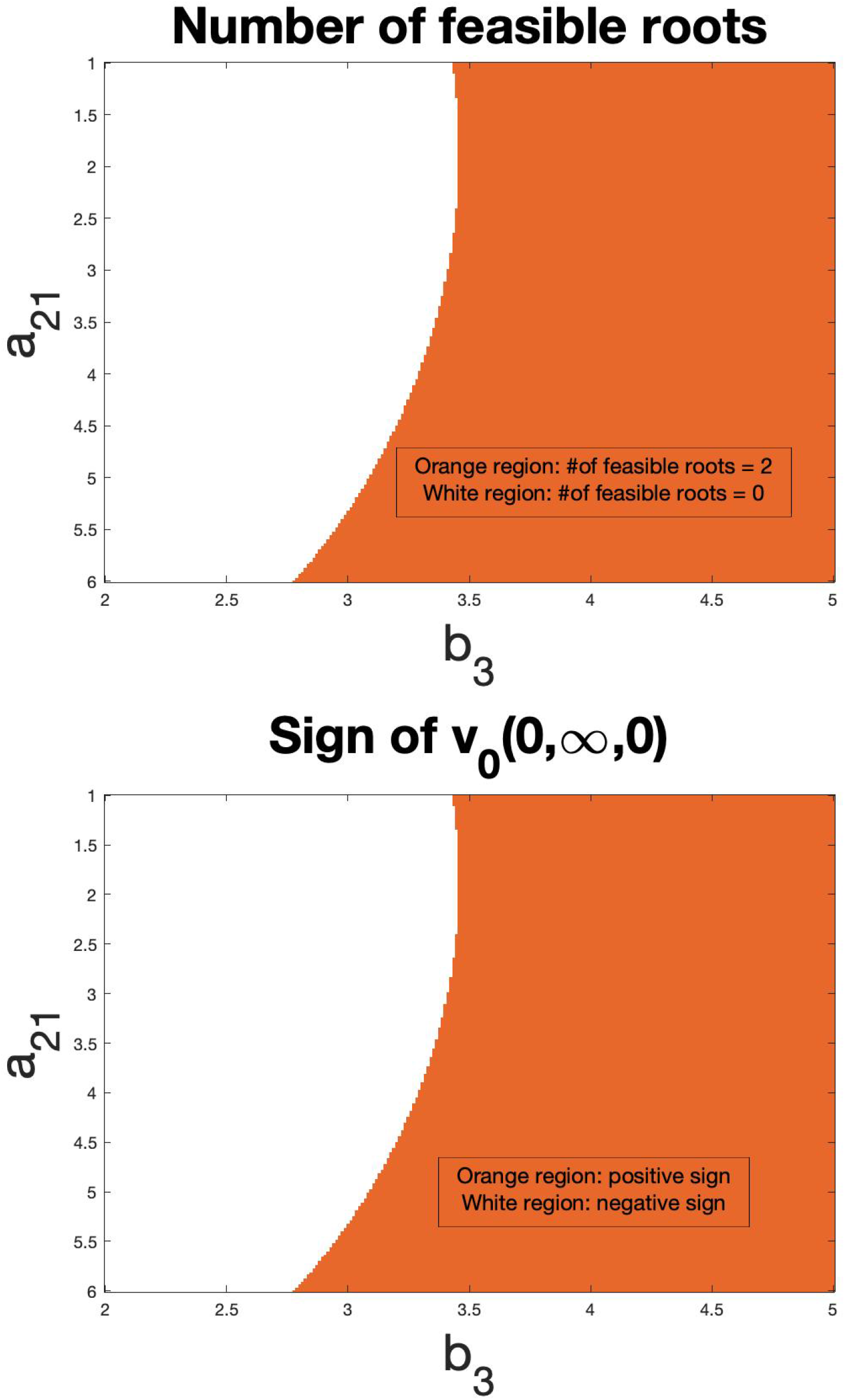
The top figure shows the number of feasible roots *F* in Lotka-Volterra model with higher-order interactions where (*r*_1_, *r*_2_, *r*_3_, *a*_11_, *a*_12_, *a*_13_, *a*_22_, *a*_23_, *a*_31_, *a*_32_, *a*_33_, *b*_1_, *b*_2_) = (1.5, −1.5, −1.5, 2, −1.5, −1.5, 2, −1.5, −1.5, −1, 1, 1, −1), *a*_21_ ∈ [1, 6] and *b*_3_ ∈ [2, 5]. The bottom figure shows the sign of *v*_0_(0, ∞, 0) with the same model and parameter values and ranges. Both figures confirm that *F* > 0 when *v*_0_(0, ∞, 0) > 0. Simulations done via solving the isocline equations numerically and checking for the feasibility of roots match the two figures displayed here.

**Supplementary Figure S4:**
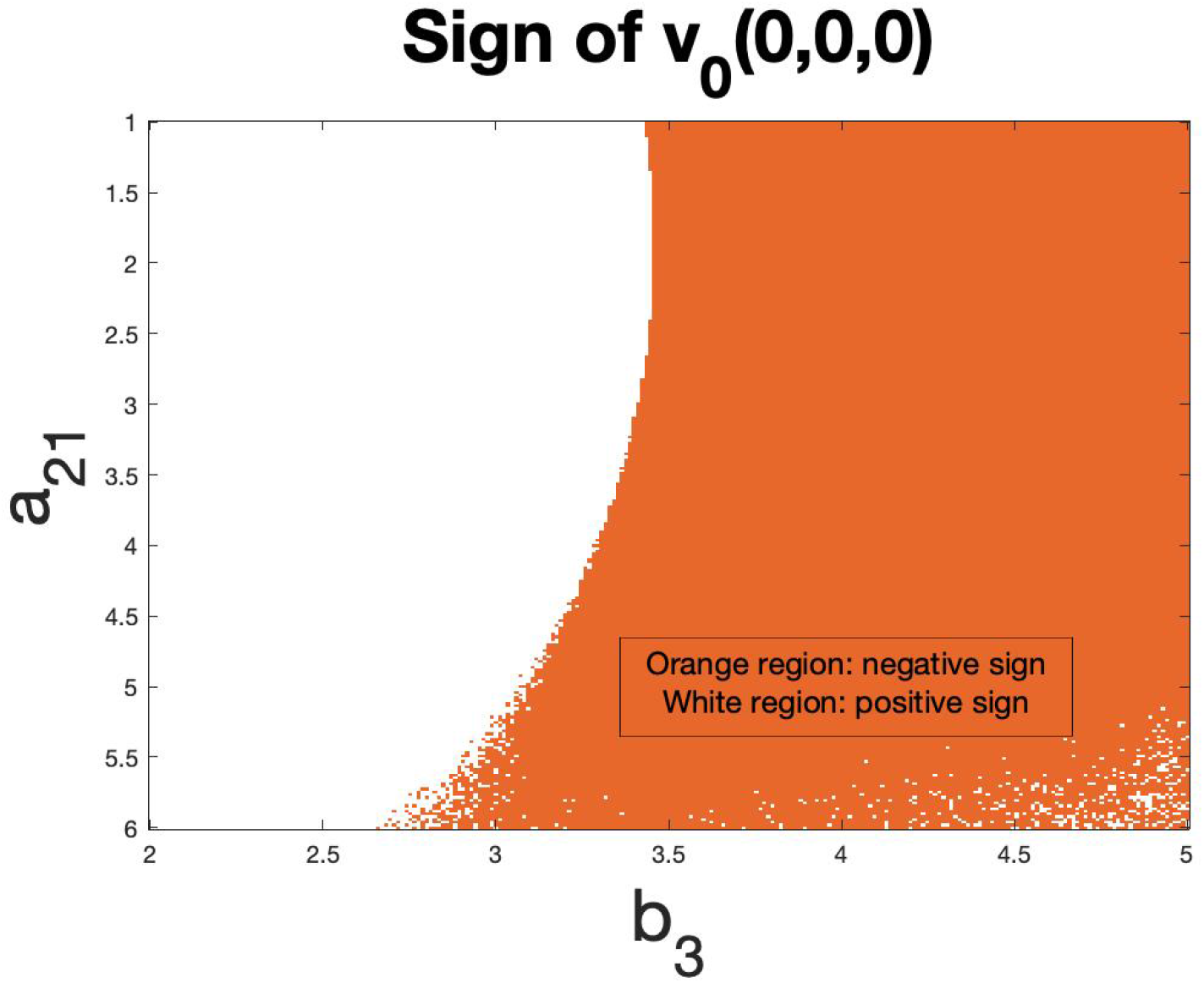
The figure plots the sign of *v*_0_(0, 0, 0) in Lotka-Volterra model with higher-order interactions when (*r*_1_, *r*_2_, *r*_3_, *a*_11_, *a*_12_, *a*_13_, *a*_22_, *a*_23_, *a*_31_, *a*_32_, *a*_33_, *b*_1_, *b*_2_) = (1.5, −1.5, −1.5, 2, −1.5, −1.5, 2, −1.5, −1.5, −1, 1, 1, −1), *a*_21_ ∈ [1, 6] and *b*_3_ ∈ [2, 5]. The shape of the figure matches the shape of the feasibility domain, yet suffers from errors.

